# Protein-coding repeat polymorphisms strongly shape diverse human phenotypes

**DOI:** 10.1101/2021.01.19.427332

**Authors:** Ronen E. Mukamel, Robert E. Handsaker, Maxwell A. Sherman, Alison R. Barton, Yiming Zheng, Steven A. McCarroll, Po-Ru Loh

## Abstract

Hundreds of the proteins encoded in human genomes contain domains that vary in size or copy number due to variable numbers of tandem repeats (VNTRs) in proteincoding exons. VNTRs have eluded analysis by the molecular methods—SNP arrays and high-throughput sequencing—used in large-scale human genetic studies to date; thus, the relationships of VNTRs to most human phenotypes are unknown. We developed ways to estimate VNTR lengths from whole-exome sequencing data, identify the SNP haplotypes on which VNTR alleles reside, and use imputation to project these haplotypes into abundant SNP data. We analyzed 118 protein-altering VNTRs in 415,280 UK Biobank participants for association with 791 phenotypes. Analysis revealed some of the strongest associations of common variants with human phenotypes including height, hair morphology, and biomarkers of human health; for example, a VNTR encoding 13-44 copies of a 19-amino-acid repeat in the chondroitin sulfate domain of aggrecan (ACAN) associated with height variation of 3.4 centimeters (s.e. 0.3 cm). Incorporating large-effect VNTRs into analysis also made it possible to map many additional effects at the same loci: for the blood biomarker lipoprotein(a), for example, analysis of the kringle IV-2 VNTR within the *LPA* gene revealed that 18 coding SNPs and the VNTR in *LPA* explained 90% of lipoprotein(a) heritability in Europeans, enabling insights about population differences and epidemiological significance of this clinical biomarker. These results point to strong, cryptic effects of highly polymorphic common structural variants that have largely eluded molecular analyses to date.

---

The human genome contains thousands of variable-number-of-tandem-repeat (VNTR) polymorphisms^1,2^, but the effects of these polymorphisms on human phenotypes are largely unknown. VNTRs are multi-allelic variants at which a nucleotide sequence (from seven to thousands of bp long) is repeated several to hundreds of times, with lengths varying among individuals. Extreme alleles of VNTRs have been implicated in diseases including progressive myoclonus epilepsy^3^ and facioscapulohumeral muscular dystrophy^4^; long alleles of shorter tandem repeats (STRs) are implicated in Huntington’s Disease^5^ and amyotrophic lateral sclerosis^6,7^. VNTRs received early attention in human genetics because the length variation they introduce is often ascertained on DNA and protein gels. However, because most VNTRs are invisible to SNP arrays and difficult to recognize or measure using short-read sequencing, VNTRs have not been considered in the large genotype-phenotype association studies that have been central to human genetics for the past 15 years. Their relationship to human trait variation is thus largely unknown, though recent work has begun to explore their relationship to gene expression^8–10^.

We hypothesized that abundant exome-sequence data might contain heretofore unappreciated information about VNTR lengths, and that VNTR alleles might also segregate on specific SNP haplotypes in ways that would make them amenable to analysis by statistical imputation^11^ in SNP-phenotype data sets from hundreds of thousands of people, such as UK Biobank (UKB)^12^.

## Exploring the phenotypic effects of coding VNTRs

We first identified candidate VNTRs by scanning the human reference genome for tandem repeat sequences^13^. We then estimated the lengths of these VNTRs in thousands of individuals’ genomes using available whole exome sequence data. Since most VNTRs are far longer than individual sequence reads, we first estimated VNTR length by measuring the numbers of sequencing reads that aligned to the tandemly repeated sequences. (We reduced technical effects on coverage estimates by normalizing each VNTR measurement to measurements from other genomes with similar exome-wide coverage profiles.) We thereby estimated “diploid VNTR content”—the sum of maternally- and paternally-derived allele lengths. In downstream analyses, we focused on 118 exon-overlapping repeats (in 118 unique genes) for which these measurements exhibited high estimates of cis-heritability, which we estimated from sibling pairs by utilizing SNP-based estimates of identity by descent (IBD) at each VNTR locus (Supplementary Table 1).

We then used these data—together with genotypes for surrounding SNPs—to infer the contribution of each haplotype to the diploid “VNTR content” measurement from each individual. Intuitively, extended SNP haplotypes provide information about which individuals in a cohort are likely to have inherited the same genomic segment from a recent common ancestor, thereby enabling resolution of diploid measurements across a cohort into allele-specific contributions; given the large SNP+exome sequence data set available from UK Biobank (*N*=488,377 SNP, *N*=49,959 exome), the accuracy of such inference is limited only by the VNTR mutation rate (mutations since recent shared ancestors) and VNTR measurement error. We developed a statistical algorithm to efficiently and accurately perform such analysis on tens of thousands of diploid VNTR measurements, using sibling IBD information to benchmark accuracy and optimize analysis parameters (Methods). Using allele-specific VNTR lengths estimated in this way for 49,959 UKB participants for whom whole exome sequence data were available^14^, we created reference haplotypes of SNP and VNTR alleles, to use as a reference panel for imputing VNTR lengths into SNP-array genotypes available for larger cohorts, including (for the current work) the remainder of the UKB cohort (*N*=437,612). The SNP+VNTR reference panel we created will be broadly available via UKB with publication of the current work (see Data availability).

We applied this approach to measure the relationship of coding VNTR alleles to 791 phenotypes in up to 415,280 unrelated UKB participants (depending on phenotype) of European ancestry. This analysis found 180 statistically significant associations (Supplementary Table 2). To determine whether such associations were driven by VNTR length variation, or by other, nearby genetic variation with which VNTR alleles are in linkage disequilibrium (LD), we performed fine-mapping analyses^15^ considering nearby genotyped and imputed variants^12,16^ as well as the VNTR. Because variation at most VNTRs arises from three or more alleles, VNTR variation was only partially correlated with individual SNPs, enabling analysis to distinguish VNTR from SNP effects in a way that has been challenging for di-allelic variants but recently possible for multi-allelic structural variants such as those at the complement component 4 (*C4A/C4B*) and haptoglobin (*HP*) loci^17–19^.

Nineteen phenotype associations, involving five distinct VNTRs (Fig. 1), exhibited strong evidence (posterior probability >0.95; Methods) that VNTR variation (rather than nearby SNPs) drove genotype-phenotype associations. These associations appeared to explain some of the largest known GWAS signals for diverse human phenotypes, including height, serum urea, and hair phenotypes (Table 1), with multiple associations exhibiting strength comparable to or exceeding that of any SNP in the genome. Four VNTRs—within exons of *ACAN, TENT5A, MUC1,* and *TCHH*—had not previously been implicated at these loci. Analysis also replicated a known association between the length of the KIV-2 repeat in *LPA* and lipoprotein(a) concentration^20^ (*P* = 4.4 x 10^-(25,121)^). All five VNTRs were genotyped and imputed accurately according to cross-validation benchmarks (Extended Data Fig. 1a and Supplementary Table 1), a conclusion that was further supported by analysis of whole-genome sequencing data (Extended Data Fig. 1b) and previous gel electrophoresis studies that reported allele-length distributions^21–23^ similar to those we inferred.

**Figure 1.**
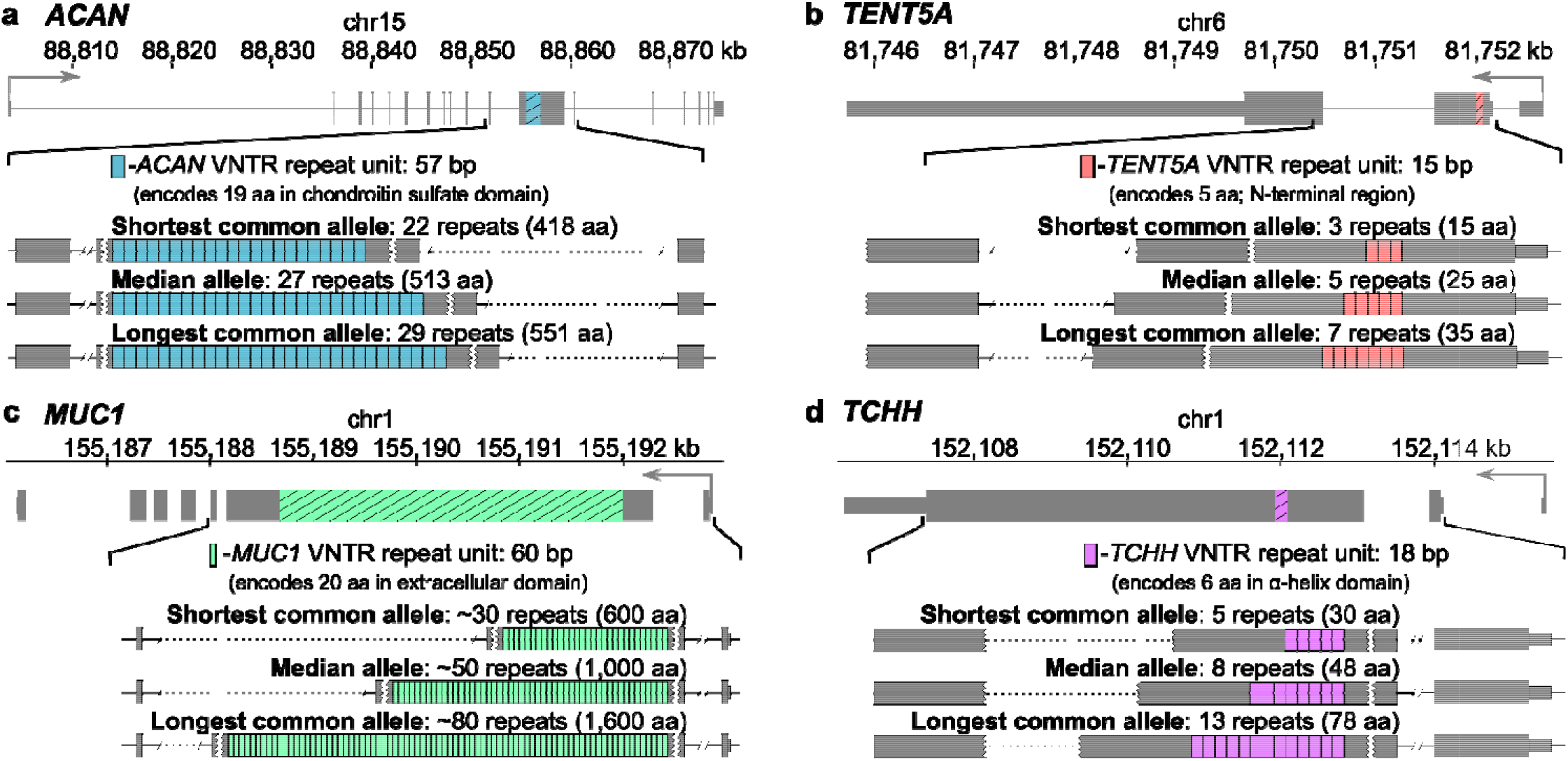
Exonic VNTRs create large-scale size polymorphisms of protein domains. VNTR alleles of varying lengths are depicted for four VNTRs (within *ACAN*, *TENT5A, MUC1,* and *TCHH*) for which we identified large-effect phenotype associations with strong implication of the VNTR by fine-mapping analysis. Gene diagrams indicate the position of the VNTR on the GRCh38 reference; callouts show examples of expanded and contracted alleles (the longest and shortest common (>1% AF) alleles identified among UKB participants of European ancestry).

**Table 1.**
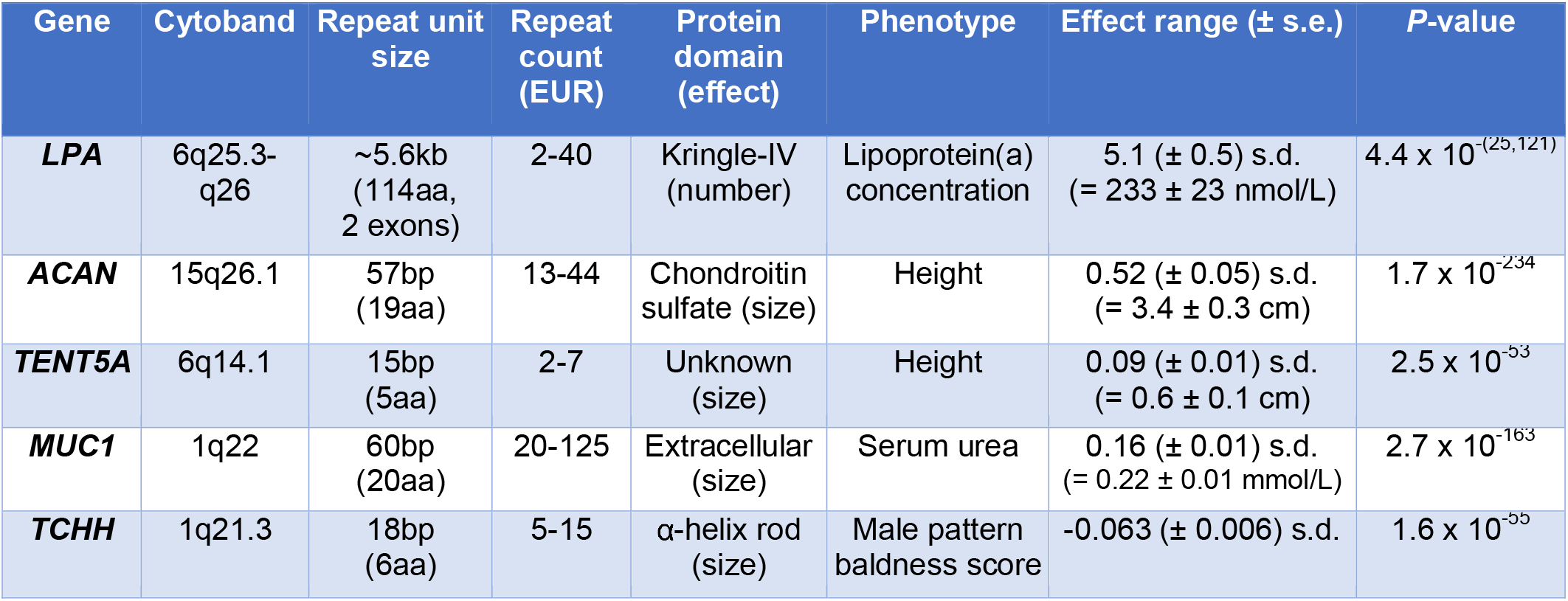
VNTRs within protein-coding sequences affect diverse human phenotypes. The table lists five protein-altering VNTRs that passed stringent fine-mapping criteria, in the sense that the VNTR (rather than nearby genomic variants) appears to be the primary driver of association signal at the locus. Here *P*-values (in linear mixed model analyses of *N*=415,280 unrelated UKB participants of European ancestry) and estimated effect-size ranges (across the longest and shortest alleles sufficiently common to be amenable to our computational analysis) are listed for the most-strongly associated phenotype; additional associations listed in Supplementary Table 2). Male pattern baldness scores were computed as in Yap et al. (2018)^59^; aa, amino acids.

| Gene | Cytoband | Repeat unit size | Repeat count (EUR) | Protein domain (effect) | Phenotype | Effect range (± s.e.) | P-value |
| --- | --- | --- | --- | --- | --- | --- | --- |
| **LPA** | 6q25.3-q26 | ~5.6kb (114aa, 2 exons) | 2-40 | Kringle-IV (number) | Lipoprotein(a) concentration | 5.1 (± 0.5) s.d. (= 233 ± 23 nmol/L) | $4.4 \times 10^{-(25,121)}$ |
| **ACAN** | 15q26.1 | 57bp (19aa) | 13-44 | Chondroitin sulfate (size) | Height | 0.52 (± 0.05) s.d. (= 3.4 ± 0.3 cm) | $1.7 \times 10^{-234}$ |
| **TENT5A** | 6q14.1 | 15bp (5aa) | 2-7 | Unknown (size) | Height | 0.09 (± 0.01) s.d. (= 0.6 ± 0.1 cm) | $2.5 \times 10^{-53}$ |
| **MUC1** | 1q22 | 60bp (20aa) | 20-125 | Extracellular (size) | Serum urea | 0.16 (± 0.01) s.d. (= 0.22 ± 0.01 mmol/L) | $2.7 \times 10^{-163}$ |
| **TCHH** | 1q21.3 | 18bp (6aa) | 5-15 | α-helix rod (size) | Male pattern baldness score | -0.063 (± 0.006) s.d. | $1.6 \times 10^{-55}$ |

## Deep fine-mapping of *LPA* variants influencing lipoprotein(a) concentration

In addition to identifying the strongest common-variant associations for several phenotypes, analysis of VNTRs made it possible to appreciate a complex interplay between VNTR length variation and other strong-effect coding variants within these genes—often (though not always) within the VNTR itself—in shaping human phenotypes.

Complex genetics involving VNTRs and SNPs at the same locus was clearly revealed in analysis of lipoprotein(a) concentration (Lp(a)), for which elevated levels are a major risk factor for coronary artery disease^24,25^. Lp(a) is almost completely heritable, with roughly half of its population variance explained by a VNTR-generated size polymorphism in the kringle-IV domain of apo(a)^20^. Each KIV-2 repeat unit (~5.6 kb) spans two exons of *LPA,* which together encode a 114-amino-acid copy of this domain. The large size of the VNTR, together with the presence of common DNA-sequence variation within the repeat units (which traced multiple distinct repeat expansions; Extended Data Fig. 2), enabled us to accurately estimate allele lengths from exome sequencing depth-of-coverage (RMSE=0.9 repeat units in cross-validation benchmarks; Extended Data Fig. 1 and Supplementary Note), recovering a multimodal distribution of KIV-2 VNTR alleles^21^ (Fig. 2). Longer alleles—with more copies of the encoded kringle repeat—are known to associate with lower Lp(a) levels^20,26^, reflecting retention of longer apo(a) isoforms in the endoplasmic reticulum^27^. As expected based on earlier work^28^, we found by sib-pair analysis that inheritance at the *LPA* locus explained most of the variance in Lp(a) measurements (*R*=0.93 in sib-pairs sharing both *LPA* alleles), with KIV-2 length explaining ~61% of this variance in a nonparametric model.

**Figure 2.**
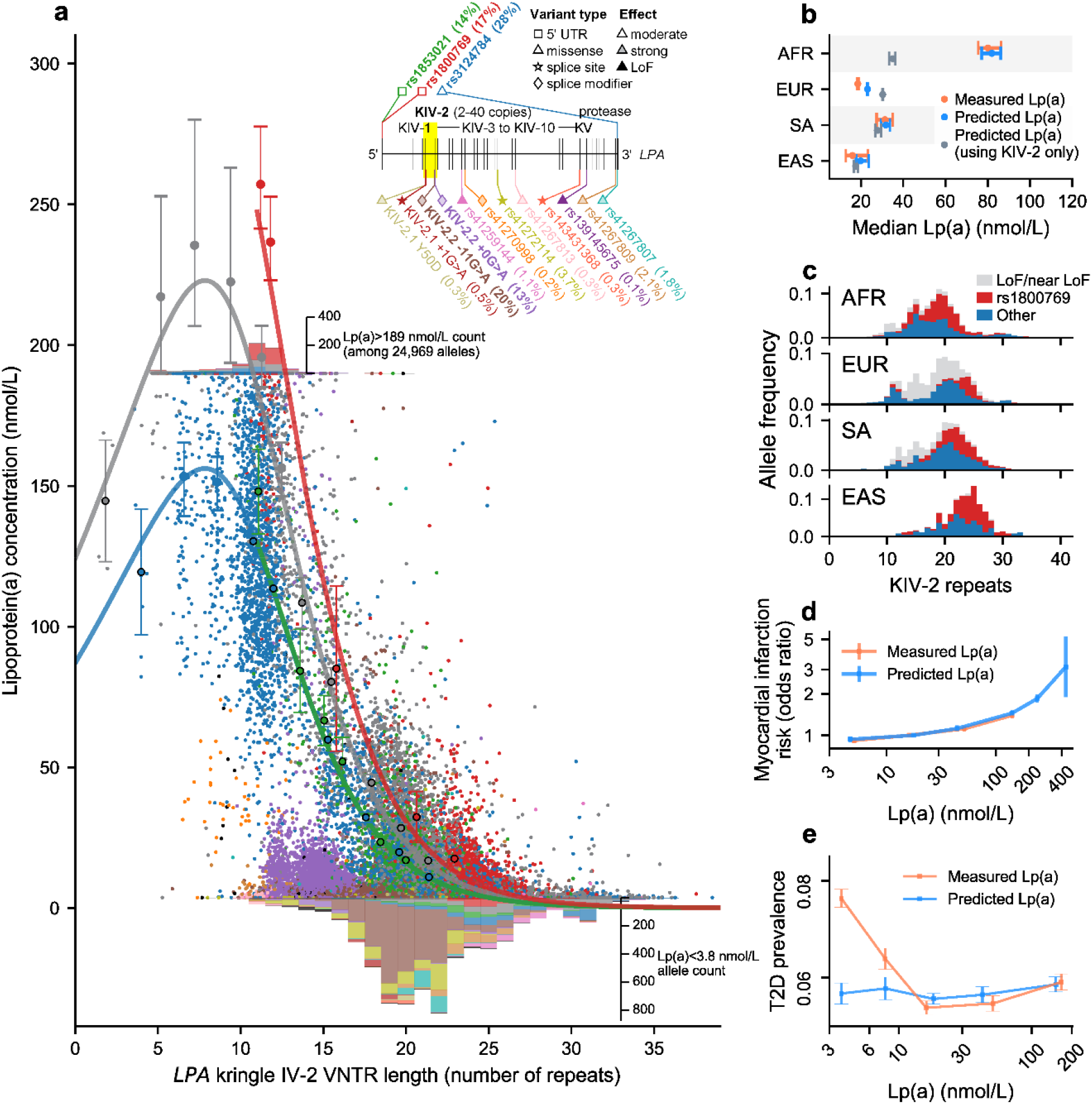
Kringle IV-2 repeat length variation and 23 *LPA* SNPs together explain ~90% of lipoprotein(a) heritability. **a,** Serum lipoprotein(a) concentration vs. KIV-2 VNTR length in an effective-haploid model of Lp(a), involving *N*=24,969 *LPA* alleles (in exome-sequenced UKB participants of European ancestry) for which the allele on the homologous chromosome was predicted to produce little or no Lp(a) (<4 nmol/L) based on KIV-2 length and/or presence of Lp(a)-reducing SNPs. The inset shows the locations of the 15 most common (unlinked) SNPs that we found (by fine-mapping analysis) to further affect Lp(a). The red, blue, green, and gray curves indicate parametric fits of Lp(a) to KIV-2 length (for noncarriers of any Lp(a)-modifying SNP in gray; carriers of rs1800769 in red; carriers of rs3124784 in blue; and carriers of rs1853021 in green) in an analysis controlling for the Lp(a)-modifying SNPs; the large points with error bars indicate mean Lp(a) for alleles in KIV-2 length bins. Small points on the plot correspond to individual *LPA* alleles, color-coded according to which fine-mapped Lp(a)-modifying SNPs they carry (black for carriers of additional rare SNPs not pictured in the inset). Histograms at top and bottom report counts of Lp(a) measurements outside the reportable range (<3.8 nmol/L or >189 nmol/L), with colors corresponding to Lp(a)-modifying SNPs carried by *LPA* alleles in these individuals (additional information in Methods). **b**, Observed and genetically predicted median Lp(a) among individuals of African (AFR; *N*=893), European (EUR; *N*=42,162), South Asian (SA; *N*=954), and East Asian (EAS; *N*=156) ancestry. **c**, *LPA* allele frequencies by ancestry. Counts for VNTR alleles in *cis* with a large-effect Lp(a)-reducing variant are colored grey; counts for VNTR alleles in *cis* with the Lp(a)-increasing 5’ UTR variant rs1800769 are colored red. **d**, Myocardial infarction risk vs. measured or genetically predicted Lp(a). **e**, Type 2 diabetes prevalence vs. measured or genetically predicted Lp(a). Error bars, 95% CIs.

To identify additional *LPA* variants that might more completely explain Lp(a), and to explore their interactions with KIV-2 length, we utilized individuals heterozygous for either of two coding variants (combined MAF=0.05) known to create null alleles that produce undetectable serum Lp(a). This approach created an effective haploid model for Lp(a), in which genetic variation on the haplotype not impaired by the common Lp(a)-null variant could be associated to Lp(a) measurements. This strategy made it possible to identify (and measure the effects of) alleles that produce unexpectedly low Lp(a), whose effects are usually made hard to measure by the much-larger contribution to Lp(a) from the allele on the homologous chromosome (Extended Data Fig. 3). We performed stepwise conditional analysis to identify *LPA* sequence variants that associated with low Lp(a) despite occurring on short or medium-length KIV-2 alleles that typically associate with higher Lp(a) levels (Methods).

These analyses identified 17 protein-altering variants which appeared to greatly reduce Lp(a) (*P*<1 x 10^−17^ for each variant); 43% of European haplotypes were affected by at least one of these variants. These variants included six variants predicted to partially or fully abolish constitutive splice sites and six missense variants that achieved the strongest associations in 12 consecutive stages of stepwise analysis; five additional rare (MAF<1%) coding variants exhibited top or near-top associations in further conditional analyses (Fig. 2a, Extended Data Fig. 4, and Supplementary Table 3). The two variants with the largest impacts on Lp(a) variation in the European population (owing to their high allele frequencies; MAF=13% and 21%) were variants within the KIV-2 region (i.e., within the VNTR) that were computationally predicted to impair splicing^29^ of KIV-2 exon 2; one of these splice variants had previously been experimentally validated^30^ (Fig. 2a and Extended Data Fig. 4). These variants appeared to reduce Lp(a) by 85% and 89%, respectively, when present within a single KIV-2 repeat unit; alleles carrying either variant on multiple repeat units within the VNTR produced nearly undetectable Lp(a) (Extended Data Fig. 5). Further fine-mapping analyses identified three other common variants (MAF=14-28%) – two in the 5’ untranslated region of *LPA* (previously observed to regulate translational activity^31,32^) and one missense variant—that associated with more modest effects on Lp(a) levels across a broad range of KIV-2 alleles (Fig. 2a and Supplementary Table 3).

The strong effects of the VNTR and SNPs at *LPA,* the large sample size of UK Biobank, and the ability to chromosomally phase all these variants accurately, made it possible to identify nonlinear and cis-epistatic effects at *LPA—*something that is often challenging to do in human genetics (because large-effect variants are often too rare to meaningfully analyze variant combinations with high statistical power and with knowledge about their chromosomal phase). For example, accounting for the effects of the 17 implicated coding variants at *LPA* showed that the well-documented inverse relationship between KIV-2 length and Lp(a)^20,27^ breaks down for very short (high-protein-level) alleles (Fig. 2a). Throughout most of the KIV-2 length range (12-24 repeats), each one-repeat-unit decrease in KIV-2 length resulted in a 37% increase in Lp(a) (Fig. 2a). However, this effect was attenuated for alleles with fewer than 12 repeats and appeared to invert around 8 repeats, with three extremely short, very rare alleles (4 or fewer repeats) exhibiting markedly lower Lp(a) levels (*P*=5.1 x 10^−19^) that were not explained by any sequence variants we identified (Fig. 2a).

Accounting for *LPA* sequence variants and the apparently nonlinear effect of KIV-2 length explained 90% of the heritable variance (83% of total variance) in Lp(a) (vs. at most ~60% of total variance in earlier work^20,33^). This model estimated allelic contributions to Lp(a) using a parametric fit of the nonlinear relationship of Lp(a) to KIV-2 copy number, with additional coding and splice variants acting multiplicatively on Lp(a) (Methods). Crucially, this analysis leveraged phase-resolved KIV-2 length estimates and SNP haplotypes to apply Lp(a)-modifying effects only to the KIV-2 allele in *cis* (i.e., on the same haplotype); these pervasive *cis*-epistatic effects were poorly modeled by the standard additive model that assumes linear effects of allele dosages (which explained only 61% of variance in Lp(a)).

Lp(a) is known to vary greatly across populations^20^, with median measurements 4-fold higher among Africans than among Europeans, but the reasons for this extensive cross-population variation have been unclear (since population differences in KIV-2 length are too small to explain them). We found that this variation was largely explained by population differences in the allele frequencies of the *LPA* sequence variants we identified (Fig. 2b). Elevated Lp(a) in UKB participants of African ancestry (median 80.1 nmol/L vs.18.5 nmol/L in Europeans) was primarily explained by the relative paucity of alleles carrying variants that greatly reduced Lp(a) (~13% of African alleles vs. ~43% of European alleles, despite ample discovery power in both populations) and the much higher frequency of an Lp(a)-increasing 5’ UTR variant among African alleles (MAF=46% vs. 17% in European alleles for rs1800769; Fig. 2c). These allele frequency differences also explained the apparent difference in shape of the Lp(a)-KIV-2 curve in different populations (Extended Data Fig. 6).

The high accuracy of genetically predicted Lp(a) (*R*^2^=0.83 in Europeans) made it possible to better understand epidemiological associations involving Lp(a). Higher genetically predicted Lp(a) has been found (by Mendelian randomization analysis) to increase cardiovascular risk^25,34^. We observed that this relationship extends to extreme Lp(a) levels: individuals with genetically predicted Lp(a)>400 nmol/L exhibited a three-fold increase in myocardial infarction (OR=3.15, 95% CI=1.9-5.2; Fig. 2d).

The much-stronger genetic prediction of Lp(a) also made it possible to identify when a known epidemiological correlation with Lp(a) was not due to Lp(a) itself. For example, low Lp(a) exhibits epidemiological association to type 2 diabetes (T2D) risk, an association with conflicting proposed interpretations^20,34^ analogous to the debate over whether HDL levels shape cardiovascular risk, or are shaped by other cardiovascular risk factors^35^. We found (by Mendelian randomization analysis in UKB) that genetically predicted Lp(a) did not associate with T2D risk, suggesting that either T2D itself, T2D-related liver comorbidities, or T2D medication use accounts for the 17% (s.e. 1%) lower levels of Lp(a) seen in T2D patients (Fig. 2e and Extended Data Fig. 7). Controlling for genetic effects almost completely by using our more accurate genetic predictor of Lp(a) levels also made it possible to identify several environmental exposures that further modified Lp(a), including alcoholic liver disease, oral contraceptives, and hormone replacement therapy (which all lowered Lp(a)) and anti-epileptic medications (which increased Lp(a); Supplementary Table 4).

## Human height strongly affected by VNTRs in *ACAN* and *TENT5A*

Human height associates with hundreds of common alleles^36^, generally with small effect sizes (individual common alleles generally explain <0.05 standard deviations). We found that size variation of a 57bp (19 amino acid) repeat in the *ACAN* gene strongly associated with height (*P*=1.7 x 10^−234^), with an effect size differential of 0.52 standard deviations (s.e. 0.05)— i.e., 3.4 centimeters—between the longest and shortest alleles sufficiently common to be amenable to our analysis (Fig. 3). SNPs at the *ACAN* locus were among the first variants in the genome to be associated with height^37^; however, no causal variants that could plausibly underlie these associations (which reached *P*=1.0 x 10^−188^ for individual SNPs vs. *P*=1.7 x 10^−234^ for the VNTR) had previously been identified.

**Figure 3.**
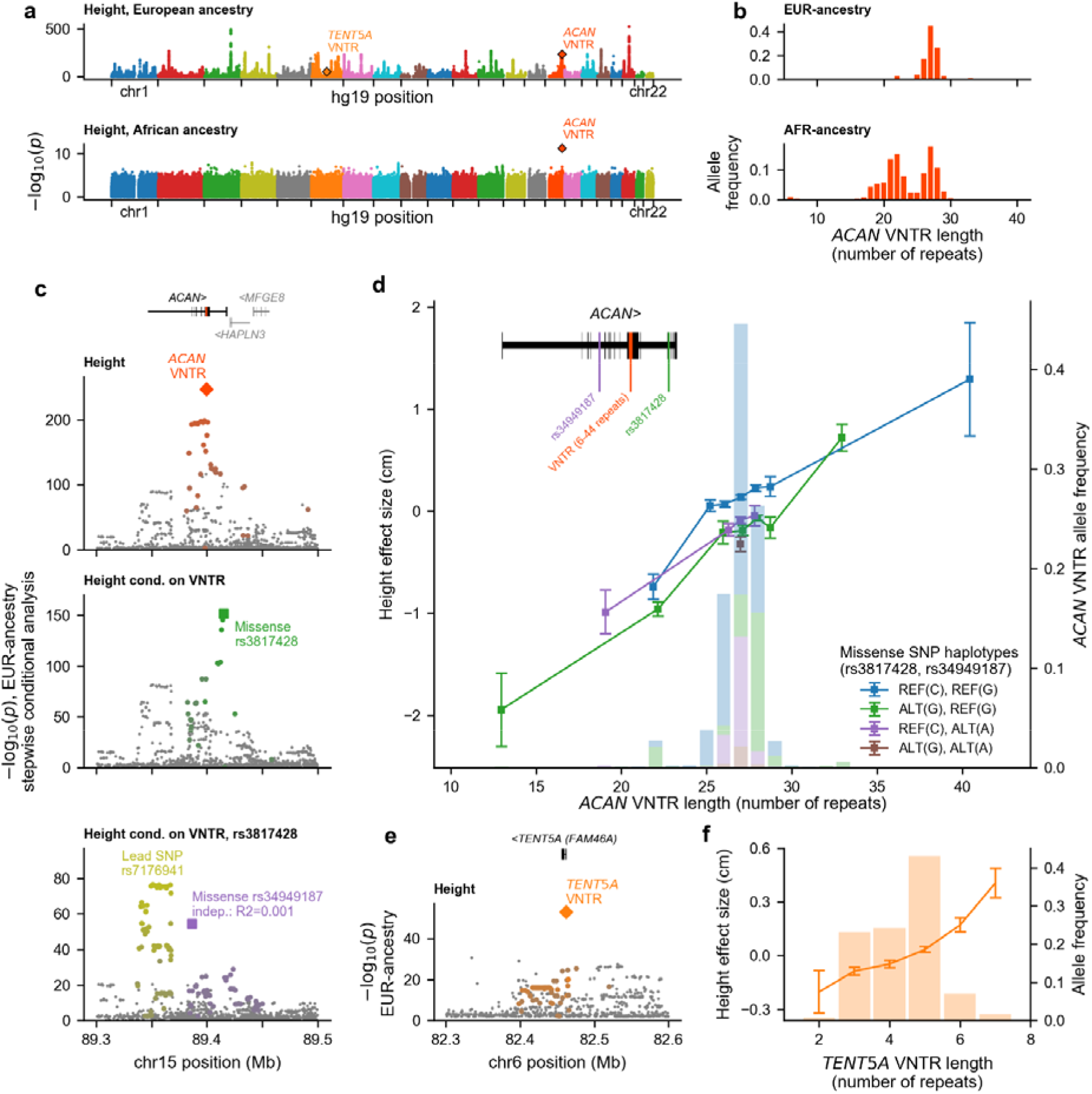
Lengths of protein-coding repeat polymorphisms in *ACAN* and *TENT5A* associate with large increases in human height. **a**, Genetic associations with height in UKB participants of European (top; EUR *N*=415,280) and African (bottom; AFR *N*=7,543) ancestry. **b**, *ACAN* VNTR allele length distributions by ancestry. **c**, Height association statistics at *ACAN* in three consecutive steps of stepwise conditional analysis (of *N*=415,280 EUR individuals). Coding mutations likely to influence height are indicated with large markers; variants in partial LD (*R*^2^>0.1) with these labeled variants are highlighted in the same color. To maximize power, height phenotypes were first adjusted for genetic predictions computed using the rest of the genome (Methods). **d**, Height effect sizes (lines, left axis) and allele frequencies (in EUR; histograms, right axis) of *ACAN* alleles defined by VNTR length and missense SNP haplotype; error bars, 95% CIs. Rare long alleles (40-42 repeats) were grouped into one bin. **e**, Height association statistics at *TENT5A.* Variants in LD (*R*^2^>0.1) with the *TENT5A* VNTR are highlighted in orange. **f**, Height effect sizes and allele frequencies of *TENT5A* VNTR alleles; error bars, 95% CIs.

To uncover this association, we estimated *ACAN* VNTR allele lengths in UK Biobank participants (RMSE ~0.85 repeat units; Extended Data Fig. 1) by analyzing counts of exome sequencing reads mapping within the VNTR, which had been sequenced to ~400x mean coverage due to the tiling of the VNTR by 12 capture probes. To maximize modeling accuracy and minimize capture bias, we independently phased and imputed copy numbers for repeat subtypes distinguished by common coding-sequence variation within the 57bp repeat unit (Extended Data Fig. 8 and Supplementary Note). Consistent with previous gel electrophoresis studies^22^, the inferred allele distribution contained common 26-, 27-, and 28-repeat alleles and low-frequency alleles with 13-33 repeats; we also identified a very short 6-repeat African allele and rare long European alleles of up to ~44 repeats (Fig. 3b,d).

Height exhibited an approximately linear relationship with length of the *ACAN* VNTR, with consistently increasing effects observed across a series of at least nine distinct VNTR allele lengths, resulting in an association signal (*P*=1.7 x 10^−234^) stronger than that of any nearby variant, explaining 0.19% of height variance among European-ancestry UKB participants (Fig. 3c,d). Moreover, among 7,543 UKB participants of African ancestry, the *ACAN* VNTR association was nearly 50% stronger than the association of any other variant in the genome (*P*=5.2 x 10^−12^ for the VNTR vs. *P*=1.4 x 10^−8^ for the strongest SNP association) and explained an even larger proportion of height variance (0.60%), primarily owing to greater VNTR length variation among UKB research participants with African ancestry (s.d.=3.7 repeats vs. 1.5 repeats in Europeans; Fig. 3b).

Aggrecan, the protein encoded by *ACAN,* is a prominent component of the extracellular matrix in growth plate cartilage and is required for normal growth plate cytoarchitecture^38^. The VNTR generates 2.4-fold size variation in aggrecan’s first chondroitin sulfate domain (CS1), a domain whose amino-acid residues are modified by long, charged polysaccharide chains that endow this extracellular matrix with key properties including the ability to hold large amounts of water. ^22^

As at *LPA,* incorporation of the *ACAN* VNTR into genetic association analysis (by stepwise conditional analysis) made it possible to identify additional genetic effects —driven *at ACAN* by two common missense SNPs (Fig. 3c and Supplementary Table 5). The two missense SNPs, which affect ACAN globular domains, had two of the top three predicted deleteriousness scores^39^ (CADD = 23.1 for rs3817428 and 27.6 for rs34949187) among common missense SNPs in *ACAN* and were corroborated by Bayesian fine-mapping^15^ analysis (posterior probability >0.99 of causality). A combined model including the VNTR and these SNPs explained 0.33% of height variance in Europeans.

Despite the strong effects of *ACAN* VNTR alleles on height, neither end of the allelic spectrum appeared to compromise ACAN function in any way detrimental to health. Whereas loss-of-function mutations in *ACAN* cause autosomal dominant skeletal disorders^40^, VNTR length variation did not associate at Bonferroni significance with any disease in UK Biobank (*P*>7 x 10^−4^), including lumbar disc degeneration, for which a previously reported association^41^ did not replicate despite ample power in UKB (*P*=0.22 in an analysis with *N*=18,982 cases). A participant homozygous for the short 6-repeat allele (AF=1.2% among participants with African ancestry) had no reported musculoskeletal disease phenotypes.

A distinct coding VNTR, in the *TENT5A* gene (previously named *FAM46A),* also associated with height. Recent work has demonstrated that TENT5A—a poly(A) polymerase in which multiple coding variants have been linked to autosomal recessive osteogenesis imperfecta (OI)^42^—polyadenylates and increases expression (in osteoblasts) of the collagen genes *COL1A1* and *COL1A2* and other genes mutated in OI^43^. The short VNTR in *TENT5A* consisted of two to seven repeats of a 15bp repeat unit (encoding a glycine-rich 5 amino acid sequence (GGDFG) in a functionally uncharacterized domain), which we genotyped in UK Biobank by adapting our methods to directly identify short alleles spanned by single sequencing reads (Supplementary Note). The *TENT5A* VNTR associated more strongly with height than did any other variant at this locus (*P*=2.5 x 10^−53^; Fig. 3e), and the six VNTR alleles exhibited monotonically increasing effects on height with increased VNTR length (with an effect range of ~0.1 s.d. (~0.6 centimeters); Fig. 3f).

## Kidney-function phenotypes shaped by a VNTR in *MUC1*

Like *ACAN,* the *MUC1* (mucin 1) gene encodes a secreted (cell-surface-associated) protein and has a VNTR that shapes the length of a heavily glycosylated extracellular domain. Ultra-rare frameshift mutations within the *MUC1* VNTR cause autosomal dominant tubulointerstitial kidney disease^44^; however, the phenotypic effects of the VNTR length polymorphism are largely unknown. In our analyses, length of the *MUC1* VNTR associated with several renal phenotypes (Fig. 4), including serum urea (*P*=2.7 x 10^−163^) and serum urate (*P*=4.7 x 10^−99^), two metabolic waste products normally removed from plasma by the kidney and excreted in urine. Longer VNTR alleles also associated with gout (*P*=3.6 x 10^−17^), a painful disease that occurs when excessive uric acid crystallizes and deposits in the joints. *MUC1* encodes a transmembrane protein (mucin 1) with cell-adhesive and anti-adhesive properties; the VNTR encodes approximately 20-125 repeats^23^ of a 60bp (20 amino acid) coding sequence that determines the length of the mucin 1 extracellular domain. *MUC1* VNTR allele lengths in UK Biobank exhibited a broadly bimodal allele distribution along with rare expanded alleles (Fig. 4c), consistent with previous work^23^. Benchmarking these allele length estimates against independent estimates from whole genome sequencing data indicated high accuracy (estimated *R*^2^ = 0.94; Extended Data Fig. 1 and Supplementary Note).

**Figure 4.**
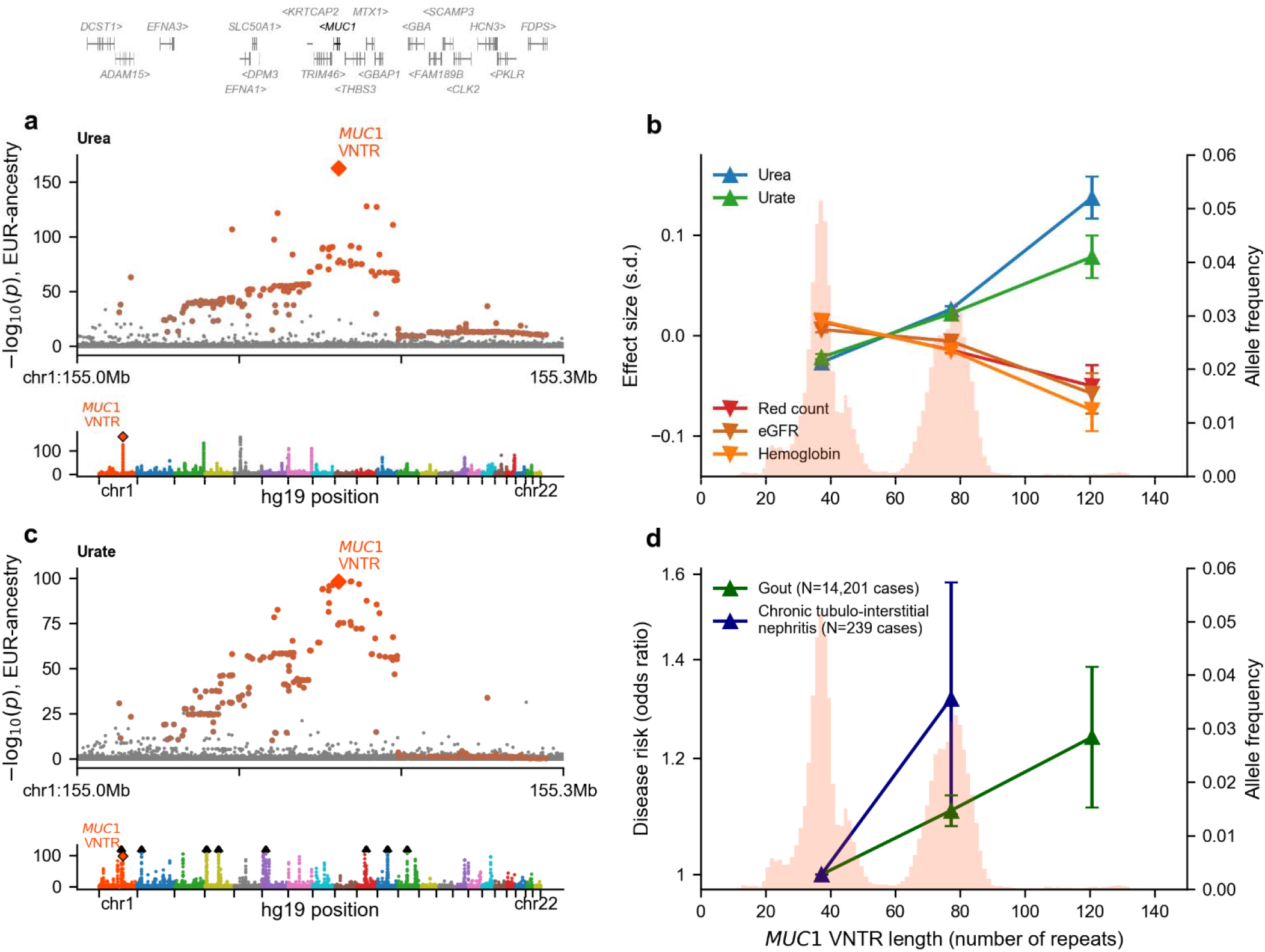
*MUC1* VNTR length associates with multiple renal phenotypes. **a** and **c**, Genetic associations with serum urea (**a**) and serum urate (**c**) at the *MUC1* locus (above in each panel) and genome-wide (below in each panel). Variants in LD with the *MUC1* VNTR (*R*^2^>0.1 with VNTR length) are highlighted in orange in the locus plots. **b** and **d**, Effect sizes (lines, left axis) and allele frequencies (histograms, right axis) of *MUC1* VNTR alleles on phenotypes related to kidney function (**b**, quantitative traits; **d**, disease traits). For estimating effect sizes, VNTR alleles were analyzed in three groups: short (<58 repeat units), long (58-100 repeat units), and very long (>100 repeat units). Error bars, 95% CIs; eGFR, estimated glomerular filtration rate. All analyses were performed on *N*=415,280 UKB participants of European ancestry.

The *MUC1* VNTR length polymorphism appeared to underlie some of the strongest, earliest reported SNP associations with serum urea and serum urate, two biomarkers of renal function that otherwise have somewhat independent genetics (genetic correlation = 0.25 (s.e. 0.01); Fig. 4a-c). For urea, the VNTR tied with a SNP on chromosome 5 for the strongest association genome-wide, explaining ~1% of heritable variance (~0.2% of total variance) in Europeans and accounting for nearly all of the association signal at the *MUC1* locus (previously reported as *MTX1-GBA*^45^; Fig. 4a). For urate, the VNTR also appeared to be the primary causal variant at a locus previously reported as *TRIM46*^46^ (Fig. 4b). Longer *MUC1* alleles associated with increasing levels of both serum urea and urate across the VNTR length spectrum, with an incompletely dominant effect on urea (*P*=2.3 x 10^−2^0 for interaction; Extended Data Fig. 9) but an additive effect on urate (*P*=0.56 for interaction).

Associations with several additional renal phenotypes indicated a complex relationship between *MUC1* VNTR length and kidney function (Fig. 4c,d). Long *MUC1* alleles (>58 repeat units) increased risk of gout (OR=1.10; 95% CI, [1.08-1.13], *P*=1.2 x 10^−16^) and chronic tubulointerstitial nephritis (OR=1.31 [1.09-1.57], *P*=3.4 x 10^−3^, which remained significant after correcting for 13 kidney diseases tested). However, *MUC1* VNTR allele length did not associate with chronic kidney disease (OR=1.01 [0.99-1.04], *P*=0.33) despite ample statistical power (*N*=14,573 cases) and only weakly influenced glomerular filtration rate as estimated from serum creatinine (beta=-0.19% [0.11-0.28%] for long vs. short alleles). Long *MUC1* alleles associated with modest reductions in red blood cell counts (beta=-0.029 s.d., s.e.=0.002, *P*=1.5 x 10^−39^) and hemoglobin levels (beta=-0.031 s.d., s.e.=0.002, *P*=9.9 x 10^−44^), possibly reflecting an impact of reduced kidney function on erythropoietin production.

## *TCHH* VNTR strongly associates with hair phenotypes

Repeat length variation in another coding VNTR—an 18bp repeat in *TCHH*—associated strongly with male pattern baldness (*P*=1.6 x 10^−55^). *TCHH* encodes trichohyalin, an intermediate filament protein that associates in regular arrays with keratin intermediate filaments and confers mechanical strength to the inner root sheath^47^. The 18bp VNTR encodes part of a highly-stabilized alpha-helix that forms an elongated rod structure^48^. A rare nonsense mutation in *TCHH* has been implicated in uncombable hair syndrome^49^, and a common haplotype containing the *TCHH* missense SNP rs11803731 (encoding a leucine to methionine substitution in TCHH) is by far the strongest genetic determinant of hair curl in individuals of European ancestry^50,51^. In UK Biobank, the *TCHH* VNTR and rs11803731 exhibited independent associations with male pattern baldness (Fig. 5a,b).

**Figure 5.**
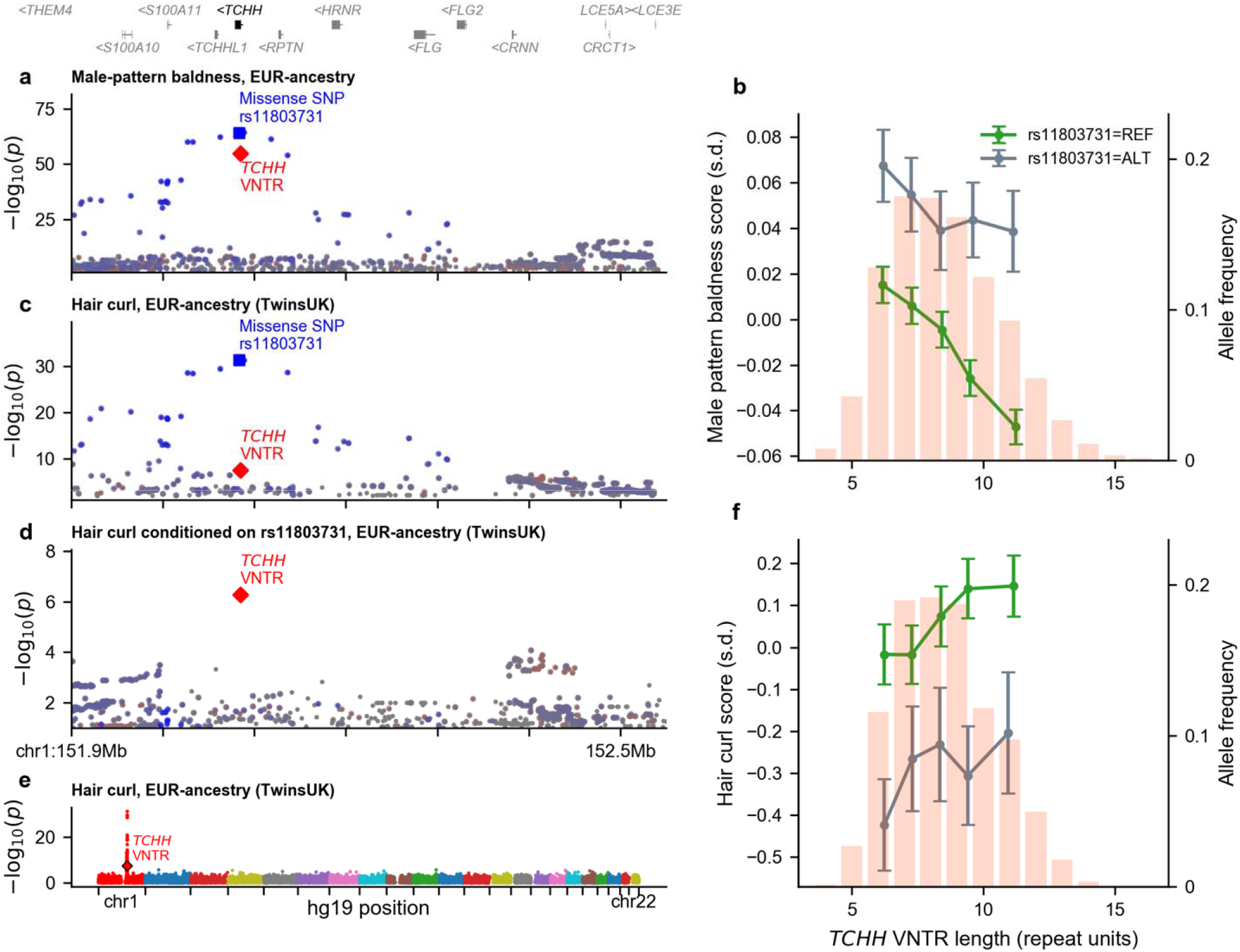
*TCHH* VNTR length and missense SNP rs11803731 associate independently with hair phenotypes. **a**, Genetic associations with male pattern baldness at *TCHH* in *N*=189,537 male UKB participants of European ancestry. Colors indicate partial LD (*R* > 0.1) with missense SNP rs11803731 (blue), the *TCHH* VNTR (red), or both rs11803731 and VNTR length (purple). **b**, Male pattern baldness effect sizes and allele frequencies of *TCHH* alleles. *TCHH* alleles were binned by VNTR length quintile and missense SNP rs11803731 status. **c**, Genetic associations with hair curl at *TCHH* in *N*=3,334 TwinsUK participants, with variants colored as in panel a. **d**, Hair curl associations in TwinsUK conditioned on rs11803731. **e**, Genome-wide genetic associations with hair curl in TwinsUK. **f**, Hair curl effect sizes of *TCHH* alleles (grouped as in panel b).

Intriguingly, the *TCHH* VNTR appeared to be hypermutable and was poorly tagged by all nearby individual SNPs (*R*^2^<0.1), leading us to wonder whether it might also contribute to hair curl in a way invisible to genome-wide association studies of this phenotype. The large size of the UKB cohort enabled accurate imputation (*R*^2^~0.7) of *TCHH* VNTR alleles into the TwinsUK cohort^52^ (*N*=3,334 genotyped individuals with hair curl phenotypes), which revealed that indeed, the *TCHH* VNTR appeared to be the human genome’s second-largest contributor to hair curl variation genome-wide (explaining ~1% of variance; *P*=3.6 x 10^−8^) after the missense SNP rs11803731 in *TCHH* (which explained ~4% of variance; Fig. 5c-f). Linkage disequilibrium with the VNTR and rs11803731 further explained a distant association previously reported near *LCE3E* (450kb upstream of *TCHH*) and previously thought to be independent of *TCHH*^51,53^ (Fig. 5b,c).

## Discussion

These results identify many strong effects of protein-coding VNTRs on human phenotypes. Most were among the strongest effects of all common variants identified for these phenotypes to date. VNTRs appeared at multiple loci to be the causal variants explaining the strongest, earliest-reported—but previously mysterious—genetic associations for multiple traits. Incorporation of multi-allelic VNTRs into fine-mapping analyses also helped identify many more functional variants at the same loci, revealing allelic series of SNP and VNTR alleles that will be powerful tools for functional studies and epidemiological research.

We described the phenotypic relationships of five VNTRs that showed the strongest evidence of causality from among 118 autosomal coding VNTRs accessible to our analytical approach. (Further work will be required to determine which of the other VNTR-phenotype associations we identified—for which fine-mapping analysis was less definitive due to reduced power or the presence of fewer VNTR alleles—are causal.)

These results are likely just the leading edge of a far-larger set of VNTR-phenotype associations that will become visible as VNTR imputation methods and reference haplotypes are applied to other SNP data sets, and then ultimately as long-read sequencing of large casecontrol cohorts comes to allow direct measurement of far more VNTRs in future long-read sequencing studies of human diseases. We explored here the kinds of common phenotypes and quantitative traits that exhibit common variation among UK Biobank participants; imputation of our reference VNTR-SNP haplotypes (see Data availability statement) could enable human genetics to analyze these VNTRs in a far-larger set of common and rare diseases. The strong associations identified motivate future studies that expand analysis to additional phenotypes and to tens of thousands more VNTRs^1,2^ that we were unable to analyze accurately—either because they exist in noncoding sequences, or are too short for depth-of-coverage to accurately measure their length variation, or are too mutable to maintain a consistent relationship to SNP haplotypes across generations. Ongoing and imminent reductions in the cost of long-read sequencing are an exciting development in this regard: future studies will yield more insights into the impact of VNTRs on human phenotypes and the mutational and evolutionary processes^54^ that produced the VNTR allelic spectra now present in human populations.

A longstanding frustration in human genetics has been that the overwhelming majority of reported genetic associations involve associations to haplotypes of noncoding and missense SNPs whose potential phenotypic contributions are challenging to dis-entangle from one another, and whose first-order molecular effects are opaque. VNTRs have several important attributes that help us overcome this longstanding challenge. First, multi-allelic VNTRs make it possible to dis-entangle the phenotypic contributions of VNTR alleles from those of nearby di-allelic variants such as SNPs with which they are in just partial LD. Second, associations to protein-coding VNTRs implicate the size and copy number of specific protein domains, leading to specific, testable hypotheses about the effects of protein domains in biological systems. Third, VNTRs provide clear directions of association, revealing whether risk is generated by having more or less of a domain. Finally, VNTRs generate natural allelic series of many common, functionally distinct alleles that can be used for dose-response studies in human tissues and cellular models. We hope that these attributes lead to many insights about the mechanisms by which gene and protein variation generate variation in human biology.

## Acknowledgments

The authors are grateful to R. Gupta, J. Hirschhorn, M. Hujoel, S. Raychaudhuri, and M. Warman for helpful discussions. This research was conducted using the UK Biobank Resource under application #40709. R.E.M. was supported by NSF grant DMS-1939015 and US National Institutes of Health (NIH) grant K25 HL150334. R.E.H. and S.A.M. were supported by R01 HG006855. M.A.S. was supported by the MIT John W. Jarve (1978) Seed Fund for Science Innovation and NIH fellowship MH124393. A.R.B. was supported by NIH fellowship F31 HL154537 and training grant T32 HG 2295-16. P.-R.L. was supported by US NIH grant DP2 ES030554, a Burroughs Wellcome Fund Career Award at the Scientific Interfaces, the Next Generation Fund at the Broad Institute of MIT and Harvard, and a Sloan Research Fellowship. TwinsUK is funded by the Wellcome Trust, Medical Research Council, European Union, Chronic Disease Research Foundation (CDRF), Zoe Global Ltd and the National Institute for Health Research (NIHR)-funded BioResource, Clinical Research Facility and Biomedical Research Centre based at Guy’s and St Thomas’ NHS Foundation Trust in partnership with King’s College London. Computational analyses were performed on the O2 High Performance Compute Cluster, supported by the Research Computing Group, at Harvard Medical School (http://rc.hms.harvard.edu).

## Methods

### UK Biobank genetic data

The UK Biobank resource contains extensive genetic and phenotypic data for ~500,000 participants recruited from across the UK^55^. We analyzed SNP and indel genotypes available from blood-derived SNP-array genotyping of 805,426 variants in 488,377 participants and subsequent imputation to 93,095,623 autosomal variants (using the Haplotype Reference Consortium, UK10K, and 1000 Genomes Phase 3 reference panels) in a subset of 487,409 participants^12^. We further analyzed exome-sequencing read alignments and genotype calls available from whole-exome sequencing (WES) of 49,960 participants^14^ (which achieved >20x coverage by 76bp paired-end reads for an average of 94.6% of targeted sites). We augmented the SNP-imputation data set with 4.9 million (predominantly rare) autosomal variants from the WES genotype call set that we previously imputed into the full cohort^16^.

### Sample filters for ancestry and relatedness

We applied strict filters to avoid confounding from population stratification and relatedness among individuals in genetic association analyses. We performed initial analyses on a stringently-filtered set of 337,466 unrelated, White British individuals identified by UKB^12^ (based on self-report and analysis of genetic principal components (PCs)) who had not since withdrawn from the study. In follow-up analyses of VNTRs exhibiting potentially causal phenotype associations, we expanded this sample set to a larger set of 415,280 participants that we identified using less-extreme filtering on ancestry and relatedness to maximize power to fine-map associations. Specifically, starting with the set of individuals who reported White ethnicity, we (i) removed PC outliers (more than six standard deviations away from the mean in any of the first 10 PCs); and (ii) removed one individual from each ≤2^nd^-degree related pair (kinship coefficient > 0.0884) previously identified by UKB^12^, prioritizing retaining individuals for whom height measurements were available. In secondary analyses of cross-population variation, we further analyzed smaller subsets of UKB participants who self-reported African, South Asian, or East Asian ancestry (comprising 1.6%, 1.9%, and 0.3% of the UKB cohort, respectively).

### UK Biobank phenotype data

We performed initial analyses on a set of 791 phenotypes that we curated from the UK Biobank “core” data set. This set of phenotypes consisted of: (i) 637 diseases with >250 reported cases (as of Oct 10, 2019) collated by UKB from several sources (self-report and accruing linked records from primary care, hospitalizations, and death registries) into single “first occurrence” data fields indexed by ICD-10 diagnosis codes; and (ii) 154 continuous and categorical traits selected based on high heritability or common inclusion in genome-wide association studies. Phenotypes in the latter set were derived from physical measurements and touchscreen interviews; blood count, lipid and biomarker panels of biological samples; and follow-up online questionnaires. For continuous traits, we performed quality control and normalization (outlier removal, covariate adjustment, and inverse normal transformation) as previously described^16,56^.

### Protein-coding VNTR ascertainment and genotyping pipeline

We identified and genotyped VNTR allele length variation from exome-sequencing data using an analysis pipeline consisting of three main steps (detailed in the Supplementary Note):

1. *Identify potential VNTR loci from repeat sequences in the human reference.* We identified approximate tandem repeats in the GRCh38 reference using two approaches: (i) Tandem Repeats Finder^13^ v4.09 (using its suggested parameters 2 5 7 80 10 50 2000 -l 6 -h to detect repeated patterns of up to 2kb); and (ii) a separate algorithm we developed to identify large, multi-kilobase repeats such as the KIV-2 VNTR in *LPA* (Supplementary Note). We filtered to autosomal loci with a repeat unit at least 9bp long that overlapped at least one exon (of any transcript) and overlapped the set of WES targets.
2. *Estimate VNTR lengths from exome-sequencing depth-of-coverage.* At each potential VNTR, we estimated diploid VNTR content (i.e., the sum of VNTR allele lengths across an individual’s two alleles) for exome-sequenced UKB participants by counting aligned reads overlapping the VNTR. To reduce technical noise in these measurements, we normalized depth-of-coverage estimates in each individual against corresponding estimates in the 200 other individuals in the cohort with closest-matching exome-wide sequencing profiles (Supplementary Note). To gauge the precision of these estimates, we computed the correlation coefficient between estimates in pairs of “IBD2” siblings who shared both haplotypes identical-by-descent (i.e., had inherited the same allele from their mother and inherited the same allele from their father).
3. *Phase and impute VNTR allele length estimates by modeling haplotype sharing.* We performed statistical phasing on estimates of diploid VNTR content to estimate haploid allele lengths, which we then imputed from the exome-sequenced participants into the remainder of the UKB cohort. To do so, we developed a computational algorithm to efficiently phase multiallelic VNTR variants (with real-valued length estimates) using surrounding SNP-haplotype information and simultaneously compute cross-validation-based benchmarks of phasing and imputation accuracy (Supplementary Note).

The first step of this pipeline (based solely on analysis of the human reference sequence) produced a list of 8,186 exon-overlapping tandem repeat sequences in the human genome. However, we expected that the large majority of these tandem repeats did not represent true protein-coding VNTRs (either because they did not exhibit allele length variation or because they did not overlap coding sequence due to imprecise endpoint-calling) or were too short to accurately genotype from sequencing depth-of-coverage data. We therefore developed a stringent filtering pipeline to create a high-confidence subset of protein-coding VNTRs suitable for analysis, ensuring that estimated lengths of these VNTRs (based on WES depth-of-coverage) exhibited heritable variation that was not the result of being contained within a larger CNV (Supplementary Note). These filters produced a final set of 118 high-confidence proteincoding VNTRs modifying 118 distinct genes (Supplementary Table 1).

### Genotyping paralogous sequence variation within VNTRs

We also sought to identify and genotype intra-allelic variation within repeat units—i.e., paralogous sequence variants (PSVs) —within the *LPA* and *ACAN* VNTRs, both to improve the accuracy of VNTR length estimates (Supplementary Note) and for downstream fine-mapping of the *LPA* locus for Lp(a). To do so, we catalogued within-repeat variation observed in exome-sequenced individuals and then adapted our genotyping, phasing, and imputation pipeline to analyze PSV copy number estimates derived from counting numbers of reference vs. alternate base calls at each such variant (Supplementary Note).

### Initial VNTR-phenotype association and fine-mapping analyses

We performed initial association tests between the 118 curated coding VNTRs and 791 phenotypes in the stringently-filtered subset of 337,466 unrelated White British participants identified by UKB. We first computed the Pearson correlation and *P*-value for each VNTR-phenotype pair. (All *P*-values reported throughout the manuscript are two-sided.) For associations passing a significance threshold of *P*<5 x 10^−8^ (commonly used in genome-wide association studies, and slightly conservative here), we recomputed association statistics using linear regression including 20 PCs as covariates, retaining 180 VNTR-phenotype pairs that remained significant at *P*<5 x 10^−8^.

To determine which of these VNTR-phenotype associations were likely to represent causal effects of VNTR allele length variation (vs. tagging of nearby causal SNPs), we first used BOLT-LMM^57^ v2.3.4 to compute linear regression association statistics for both the VNTR and all nearby SNPs and indels imputed by UKB (within 500kb of the VNTR) using a standard set of covariates (assessment center, genotype array, age, age squared, sex, and 20 PCs). We then applied the Bayesian fine-mapping software FINEMAP^15^ v1.3.1 (options --corr-config 0.999 --sss --n-causal-snps 5) to estimate the likelihood of causality for the VNTR, accounting for linkage disequilibrium with up to 2,000 of the most strongly associated nearby variants. The results of these analyses are summarized in Supplementary Table 2.

### Follow-up analyses of potentially causal VNTR-phenotype associations using refined genotypes and phenotypes

For each of six distinct VNTRs involved in 20 VNTR-phenotype associations that were assigned a high probability of causality (>0.95), we further optimized estimates of VNTR allele lengths by refining VNTR boundaries, carefully modeling biases in exome-sequencing read-capture induced by variation within repeat units, and (for the *TENT5A* VNTR) incorporating read-level information from single sequencing reads that spanned short alleles (Supplementary Note). These genotyping optimizations either increased or maintained the evidence for causality of all associations except one, between height and a VNTR in *RRBP1,* at which the posterior probability of causality dropped to 0.33 (Supplementary Note); we therefore dropped this association from further analysis, leaving nineteen VNTR-phenotype associations involving five VNTRs (*LPA*, *ACAN, TENT5A, MUC1,* and *TCHH).*

We curated three derived phenotypes for follow-up analyses based on relevance to the associations that fine-mapped to these VNTRs. For serum lipoprotein(a), the phenotype coding provided by UKB had coded 17% of Lp(a) measurements as missing due to falling outside the reportable range (3.8-189 nmol/L). To enable analysis of individuals with such measurements, we incorporated binary information available about whether Lp(a) had been below vs. above the reportable range by creating a “cropped Lp(a)” phenotype in which we assigned Lp(a) values of 3.7 or 190 nmol/L to such individuals; we carefully modeled the effect of this cropping in subsequent association and fine-mapping analyses (Supplementary Note). We also computed an estimated glomerular filtration rate (eGFR) phenotype (relevant to *MUC1*) from serum creatinine measurements using the MDRD study equation^58^, and we computed a male pattern baldness score phenotype (relevant to *TCHH*) following previous work^59^.

We performed follow-up analyses using the improved VNTR genotypes on an expanded set of 415,280 PC-filtered, unrelated European-ancestry participants (described above) and further optimized statistical power^60^ by performing linear mixed model association analysis of top phenotype associations using BOLT-LMM (Table 1 and European-ancestry genome-wide Manhattan plots in Fig. 3a and Fig. 4a,c; for African-ancestry genome-wide association analysis of height in Fig. 3a, we ran linear regression using our standard covariates). To emulate mixed model association power in analyses exploring the effects of specific VNTR alleles and SNPs at the *ACAN* and *TENT5A* loci for height (Fig. 3c-f), we performed analyses on height residualized for polygenic predictions of height from array-typed SNPs (omitting those within 2Mb of each VNTR) that we generated using BOLT-LMM (--predBetasFile) in 10-fold cross-validation^61^.

### Logistic regression analyses of VNTR associations with disease outcomes

We performed logistic regression to further investigate associations between genetically predicted Lp(a) and myocardial infarction (I21) and type-2 diabetes (E11), between *ACAN* VNTR length and intervertebral disk disorders (M51), and between *MUC1* VNTR length and 13 kidney diseases with at least 100 cases (I12, M10 = gout, N00, N02, N03, N04, N05, N11 = chronic tubulo-interstitial nephritis, N17, N18 = chronic kidney disease, N19, N25, and Q61, using the ICD-10 coded disease phenotypes collated by UKB). For *LPA* and *MUC1,* we binned data by the variable of interest (genetically predicted Lp(a) or *MUC1* VNTR length), included an indicator variable for each bin (omitting the reference bin; Lp(a) between 10-30 nmol/L in Fig. 2d and *MUC1* VNTR copy number <58 in Fig. 3d), and reported the effect size estimated for each bin. For *ACAN,* we analyzed VNTR length as a quantitative variable. We included age, age squared, and sex as covariates in these analyses.

### Estimating effects of VNTR alleles on quantitative traits

To estimate effect sizes of VNTR alleles or groups of alleles in analyses of quantitative traits, continuous phenotypes effects for highlighted VNTR-phenotype relationships, we binned alleles by VNTR length and SNP-haplotypes (when additional likely-causal SNPs had been identified), plotting the phenotypic mean among individuals carrying an allele in the bin (counting homozygous carriers twice) against the bin-wise median VNTR length (Figures 2a, 3d,f, 4b, and 5b,f). In Fig. 2a, we extrapolated mean Lp(a) in bins containing a large fraction (>15%) of Lp(a) measurements that exceeded the reportable range (>189 nmol/L) and had been cropped. We performed this extrapolation based on the median Lp(a) value in the and the assumption that Lp(a) within a bin was log-normally distributed with a *σ* = 0.304 (which appeared empirically to fit well across a broad cross-section of bins with fewer cropped values). In Fig. 3d (visualizing the effects of *ACAN* alleles on height), we rounded allele length estimates to the nearest integer and plotted integer bins with MAF>0.5% as well as extreme allele bins (for the rare 13- and 19-repeat alleles and very long alleles containing 40-42 repeats).

### Modeling lipoprotein(a) concentration from KIV-2 VNTR allele lengths and *LPA* sequence variants

Even though Lp(a) is almost completely determined by allelic variation at *LPA,* the specific *LPA* sequence variants that influence Lp(a) and the way that they interact to determine lipoprotein(a) concentration have remained elusive^20^. Part of the challenge of fine-mapping the *LPA* locus is the need for accurate genotyping of both KIV-2 length variation and SNP variation within the KIV-2 repeat (i.e., PSVs), which has not been available to previous large-scale studies. A further challenge is the multiple forms of nonlinearity that complicate the relationship between *LPA* sequence variation and Lp(a): (i) the nonlinear relationship of KIV-2 length with Lp(a) (even controlling for other *LPA* sequence variants), and (ii) the nonlinearities induced by the allele-specific nature of apo(a) production from individual *LPA* alleles (such that Lp(a)-modifying sequence variants on one chromosome exhibit effects that depend on the length of the KIV-2 repeat on that chromosome, while having no effect on the apo(a) production of the *LPA* allele on the other (homologous) chromosome). For these reasons, although well-powered fine-mapping studies that have applied standard stepwise conditional analyses adjusted for KIV-2 length have identified dozens of conditionally independent SNP associations at *LPA,* these lists have mostly contained noncoding variants unlikely to have causal effects^62,63^.

To fully leverage our comprehensive genotyping and imputation of phased KIV-2 allele lengths, PSVs in and near KIV-2 exons, and SNPs and indels at *LPA* in ~500,000 UKB participants— and to accurately model Lp(a) measurements that had been cropped to the range 3.8-189 nmol/L—we developed novel statistical methods to fine-map the complex association pattern at *LPA* to causal SNPs, and then to perform predictive modeling of Lp(a) from genotypes of these SNPs together with KIV-2 repeat lengths. Briefly, we first performed fine-mapping within an effective-haploid model of Lp(a) created by carriers of alleles producing little or no Lp(a). This framework isolated contributions of individual *LPA* alleles, greatly elucidating the effects of variants that substantially reduced Lp(a) (Extended Data Fig. 3). Stepwise conditional analyses within this framework identified an allelic series including 18 protein-altering variants and 3 variants in the 5’ UTR of *LPA* that each appeared likely to causally influence Lp(a) levels (based on achieving top or near-top association strengths in successive steps of analysis); 2 additional protein-truncating variants within KIV-2 exons had effects mostly masked by linkage disequilibrium with a canonical splice site variant^64^ (Supplementary Table 3 and Supplementary Note). Second, we created an intuitive model that accurately predicted Lp(a) as a sum of allelic contributions determined by KIV-2 length and the combination of alleles of the 23 likely-causal *LPA* SNP and indel variants carried on each haplotype. This model consisted of a lowdimensional parametrization of the “baseline curve” relating KIV-2 length to Lp(a) (in the absence of other Lp(a)-modifying variants) on top of which SNP modifiers exerted multiplicative effects (Supplementary Note).

We compared the above model of Lp(a) to two simpler models corresponding to standard analyses: a model predicting Lp(a) from KIV-2 length alone (used in Fig. 2b) and a linear model using KIV-2 length together with the 23 likely-causal *LPA* variants we identified from fine-mapping. To model Lp(a) from KIV-2 length alone, we first estimated contributions of KIV-2 alleles to Lp(a) in a SNP-unaware manner: we binned alleles solely by KIV-2 length in 2-repeat-unit windows, and we then averaged Lp(a) measurements for the alleles in each bin carried by individuals with a low-Lp(a) allele on the homologous chromosome (i.e., alleles plotted in Fig. 2a). As in the model above, we then predicted a given individual’s Lp(a) by summing the contribution of the two alleles (and cropping values outside the reportable range when performing comparisons against measured Lp(a)). In the linear model, we modeled Lp(a) as a linear combination of diploid genotypes (for the 23 *LPA* SNP and indel variants) and diploid KIV-2 content, fitting this model using linear regression against measured Lp(a).

### Association analysis of medication use and liver diseases with altered Lp(a) levels

To investigate potential effects of exposures including medication use and liver disease on lipoprotein(a) levels, we tested these exposures for association with differences between observed and genetically-predicted Lp(a) in UK Biobank. These analyses were well-powered because genetically-predicted Lp(a) explained ~80% of Lp(a) variance in participants of European ancestry (including those not in the exome-sequenced cohort, for whom we imputed KIV-2 length and *LPA* variants), thus serving as a proxy for baseline Lp(a) (prior to the exposure). We computed the log ratio of observed vs. predicted Lp(a), adjusted for age, sex, and 20 PCs, in 210,755 UK Biobank participants with predicted Lp(a) between 10-100 nmol/L (to avoid bias due to cropping of Lp(a) measurements to 3.8-189 nmol/L) in the unrelated, PC-filtered European-ancestry cohort. For the subsets of individuals who reported taking each of 1,314 medications (taken by at least 10 individuals analyzed), we computed a *z*-test to determine whether or not medication use associated (at Bonferroni significance, *P*<4 x 10^−5^) with a change in the log-ratio of observed vs. predicted Lp(a); if so, we exponentiated this change (and its 95% CI) and subtracted 1 to obtain non-log-scale changes in Lp(a). We performed analogous analyses for liver diseases (K70-K77) with at least 100 cases and for type 2 diabetes for reference. We note that these association analyses do not prove causality, which appears plausible or has been reported for many of the associations but is less clear for others; e.g., both T2D medications and T2D associate with similar reductions in Lp(a), leaving uncertain whether the associations are driven by medication use, T2D itself, or a T2D-related comorbidity (Supplementary Table 4).

### Association analysis of *TCHH* VNTR length variation with hair curl in TwinsUK

To explore the potential association of the *TCHH* 18bp (6 amino acid) repeat with hair curl (which was not phenotyped in UK Biobank), we analyzed 3,334 TwinsUK participants for whom both SNP-array genotypes and hair curl phenotypes were available and who did not report non-White ancestry. Hair curl phenotypes on a 4-point scale had previously been collected from two questionnaires (Q18_10, available for 3,015 of the 3,334 individuals we analyzed, and Q19_36, available for 1,689 of the individuals we analyzed). We normalized each hair curl phenotype separately in males and females by regressing out age, mean-centering, and dividing by the standard deviation; for individuals with both hair curl phenotypes available, we then averaged the two normalized phenotypes.

SNP-array genotyping had previously been performed using either an Illumina 610K or 317K array, which had low overlap with the Affymetrix arrays used by UK Biobank. To enable imputation of the *TCHH* VNTR from our reference panel of UK Biobank exome-sequenced participants, we first imputed SNPs in the region of chromosome 1 surrounding *TCHH* (chr1:140-165Mb) using the TOPMed imputation server^65,66^. (Prior to imputation, we excluded a small fraction of A/T or C/G SNPs to avoid potential strand-flipping.) After imputation, we improved the phasing of the TOPMed-imputed TwinsUK haplotypes at UK Biobank-typed SNPs by setting imputed genotypes that were <95% confident (i.e., 0.05<HDS<0.95 for either haploid dosage) to missing and then rephasing non-missing SNP genotypes using Eagle^67^ v2.4.1 --Kpbwt=100000, using all UK Biobank phased haplotypes^68^ as a reference panel. We then imputed *TCHH* VNTR allele lengths into these rephased SNP-haplotypes using the exome-sequence UKB participants as a reference panel as in our other analyses (Supplementary Note), with only the slight change of ignoring genotypes that had been set to missing when computing identity-by-state (IBS) sharing between reference and target haplotypes.

We performed linear mixed model association tests between the normalized, merged hair curl phenotype and the *TCHH* VNTR, imputed SNPs, and array-typed SNPs (with missingness <0.1) using BOLT-LMM with genotyping array as a covariate. Mixed model association analysis was necessary to account for substantial relatedness among TwinsUK participants (~1,000 monozygotic or dizygotic twin pairs among the 3,334 individuals we analyzed).

### Data availability

Individual-level VNTR allele length estimates (resolved to phased SNP-haplotypes) and genetically predicted Lp(a) values will be returned to the UK Biobank resource for download by approved researchers. Access to the following data resources is available by application: UK Biobank (http://www.ukbiobank.ac.uk/); Twins UK (https://twinsuk.ac.uk/); the Haplotype Reference Consortium imputation panel (http://www.haplotype-reference-consortium.org/).

### Code availability

The following publicly available software packages were used to perform analyses in this work: Eagle2 (v2.3.5), https://data.broadinstitute.org/alkesgroup/Eagle/; Minimac4 (v1.0.1), https://genome.sph.umich.edu/wiki/Minimac4; BOLT-LMM (v2.3.4), https://data.broadinstitute.org/alkesgroup/BOLT-LMM/; FINEMAP (v1.3.1), http://www.christianbenner.com/; plink (v1.9 and v2.0), https://www.cog-genomics.org/plink2/; Tandem Repeats Finder (v4.09.1), tandem.bu.edu/trf/trf.html; the TOPMed Imputation Server, https://imputation.biodatacatalyst.nhlbi.nih.gov/; BLAT (v35), http://hgdownload.soe.ucsc.edu/admin/exe/. Code and scripts used to perform the downstream analyses described above are available from the authors upon request.

