## Extended Data Figures for "Protein-coding repeat polymorphisms strongly shape diverse human phenotypes"

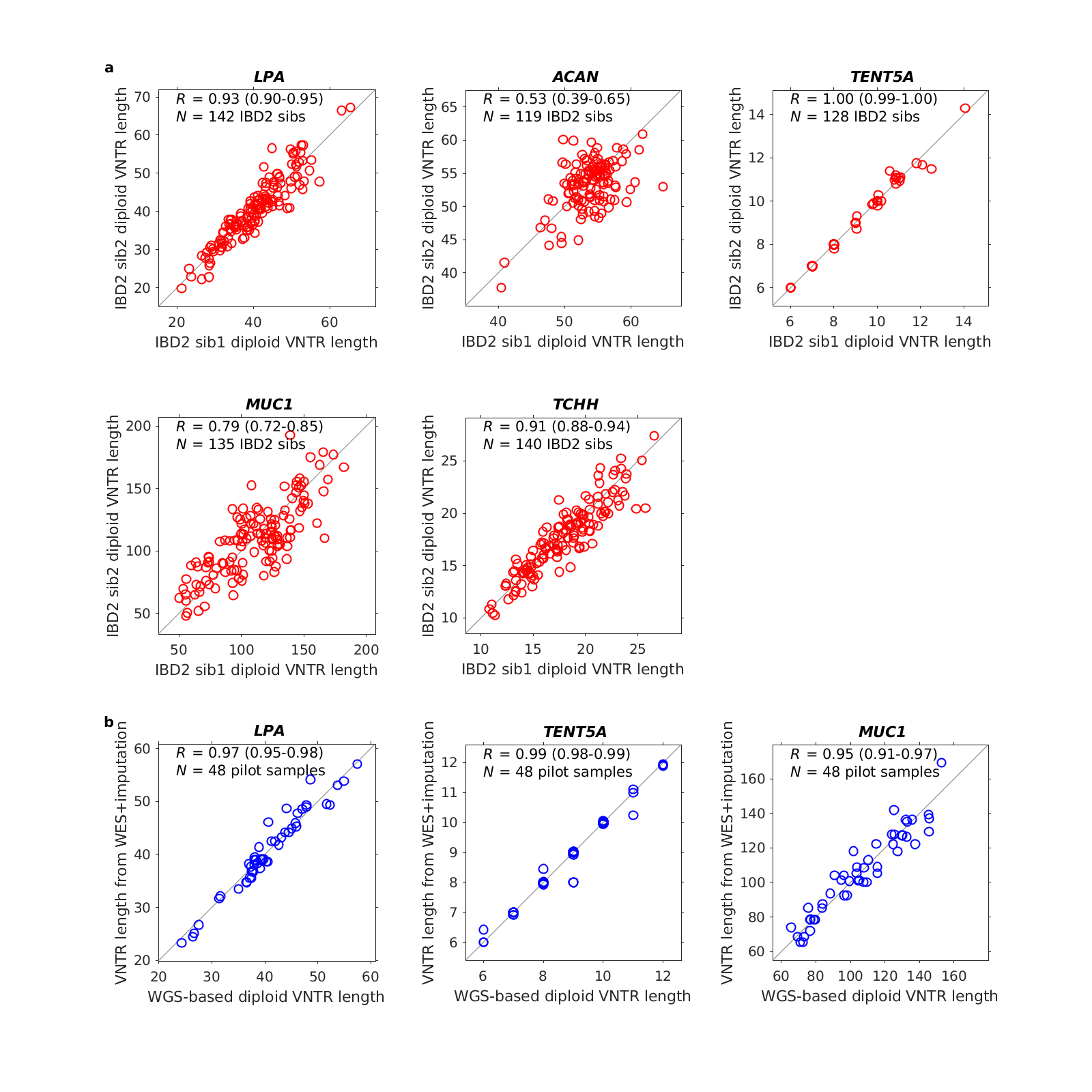

**Extended Data Figure 1. Benchmarks of VNTR genotyping and imputation accuracy. a**, Consistency of diploid VNTR length estimates (before phasing, which simultaneously refined allele length estimates) between pairs of siblings sharing both VNTR alleles (i.e., IBD2). **b**, Consistency of VNTR lengths estimated from exome sequencing vs. whole-genome sequencing (available from a pilot WGS analysis of *N*=48 UK Biobank participants, of which 6 had been exome-sequenced; imputed VNTR allele length estimates are plotted for the remaining 42 participants). WGS data enabled accurate diploid VNTR length estimation at *LPA* and *MUC1* (which have high allele length variance) and at *TENT5A* (because all alleles were spanned by 151bp sequencing reads). *ACAN* and *TCHH* allele lengths could not be accurately estimated from WGS.

**a**	🡨 Direction of transcription

Human reference coordinates 🡪

|  | **Intron position (… +1)** | **KIV-2 exon 1 position (160 … 1; minus strand)** | | | **Intron position**  **(-1 …)** | | | |
| --- | --- | --- | --- | --- | --- | --- | --- | --- |
| **Repeat**  **Type** | **+119** | **86** | **41** | **14** | **-16** | **-17** | **-23** | **-26** |
| **A** | T (**A1**)  C (**A2**) | T | A | T | G | G | C | T |
| **B** |  | A | G | C | A | A | - | A |
| **C** |  | T | G | C |  |  |  |  |
|  | PSV distinguishing A1/A2 subtypes | Synonymous PSVs within KIV-2 exon 1 distinguishing A/B/C repeat types | | | Additional intronic PSVs distinguishing A/B (C sequence not in ref) | | | |

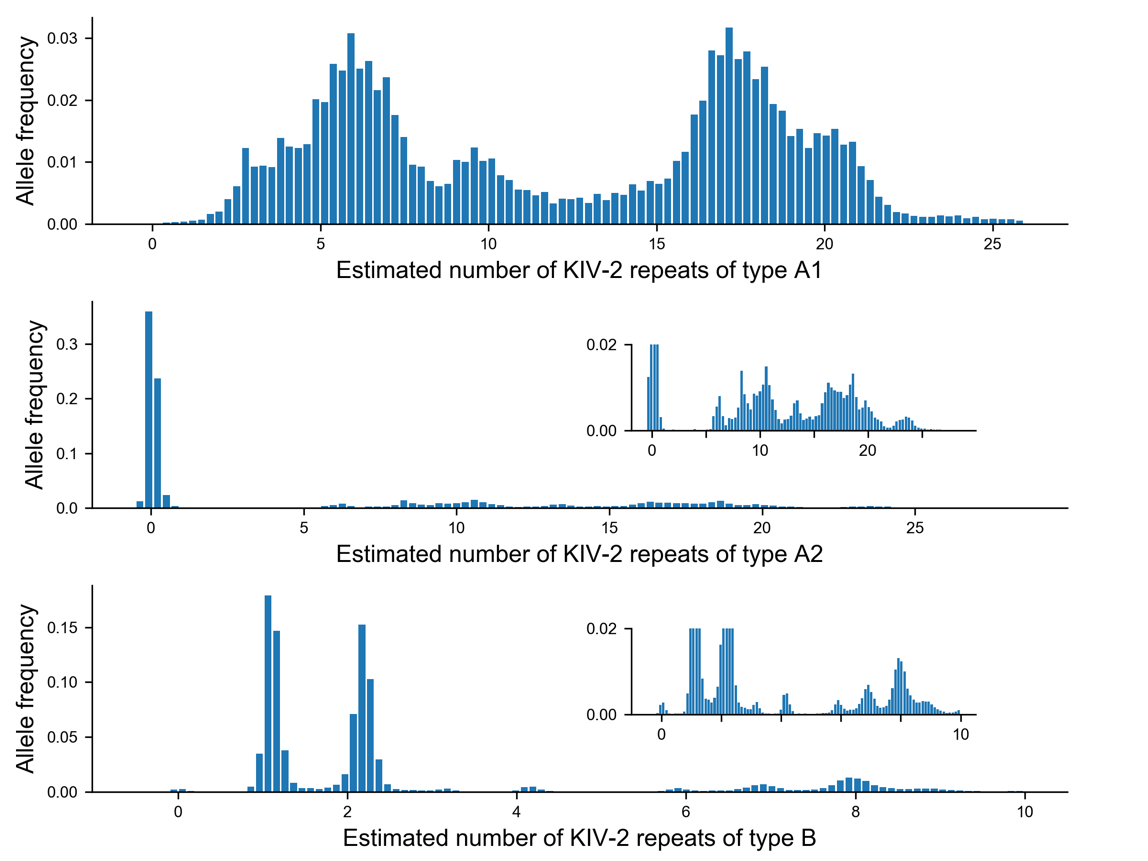
**b**

**Extended Data Figure 2. Genotyping *LPA* VNTR repeat subtypes. a**, Repeats within the *LPA* VNTR have previously been classified into three repeat types (A, B, C) based on sequence variation at three synonymous paralogous sequence variants (PSVs) within KIV-2 exon 1. A nearby PSV located 119bp into the downstream intron further subdivides repeat type A into two common subtypes, which we called A1 and A2. We separately estimated allelic copy numbers for repeat types A1, A2, and B by counting sequencing reads that mapped uniquely to one of these three repeat sequences. (We did not specially treat repeat type C, which only comprises ~1% of all repeats (Coassin et al. 2018); reads generated from type-C repeats therefore contributed to a mixture of the other repeat type counts depending on which PSVs they spanned, which determined whether they aligned best to repeat A1, A2, or B.) **b**, Separately genotyping and phasing each repeat type produced multimodal copy number distributions with some haplotypes carrying large expansions of repeat types A2 and B. The distribution of type-B repeat copy number estimates exhibited clear near-integer modes. Allele distributions for *N*=34,418 exome-sequenced, unrelated British UKB participants shown.

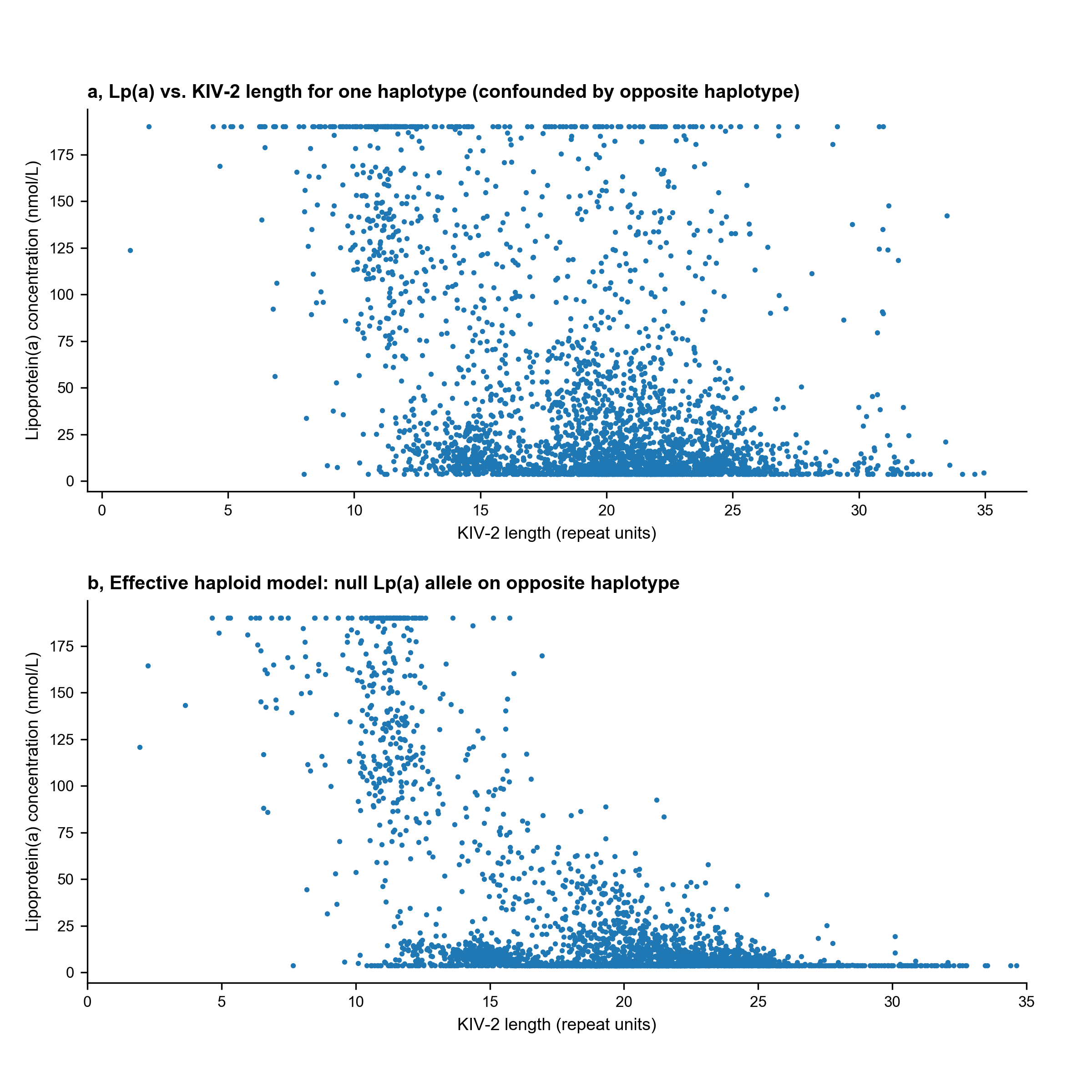

**Extended Data Figure 3. An effective-haploid model of Lp(a) created by carriers of null-Lp(a) alleles isolates contributions of individual *LPA* alleles. a,** Scatter plot of Lp(a) vs. estimated kringle IV-2 length (of one allele) in exome-sequenced UK Biobank participants. The relationship between KIV-2 length and Lp(a) is blurred by the contribution of the *LPA* allele on the “opposite” chromosome (i.e., the homologous chromosome inherited from the other parent). **b,** Scatter plot of Lp(a) vs. KIV-2 length restricted to *N*=3,343 alleles for which the opposite haplotype carries a low-frequency variant known to produce a null-Lp(a) allele (rs41272114 (Ogorelkova et al. 1999) or rs41259144 (Morgan et al. 2020)). In both panels, alleles are drawn from exome-sequenced, unrelated, British UKB participants; in panel **a**, alleles are down-sampled to match the count in panel **b**.

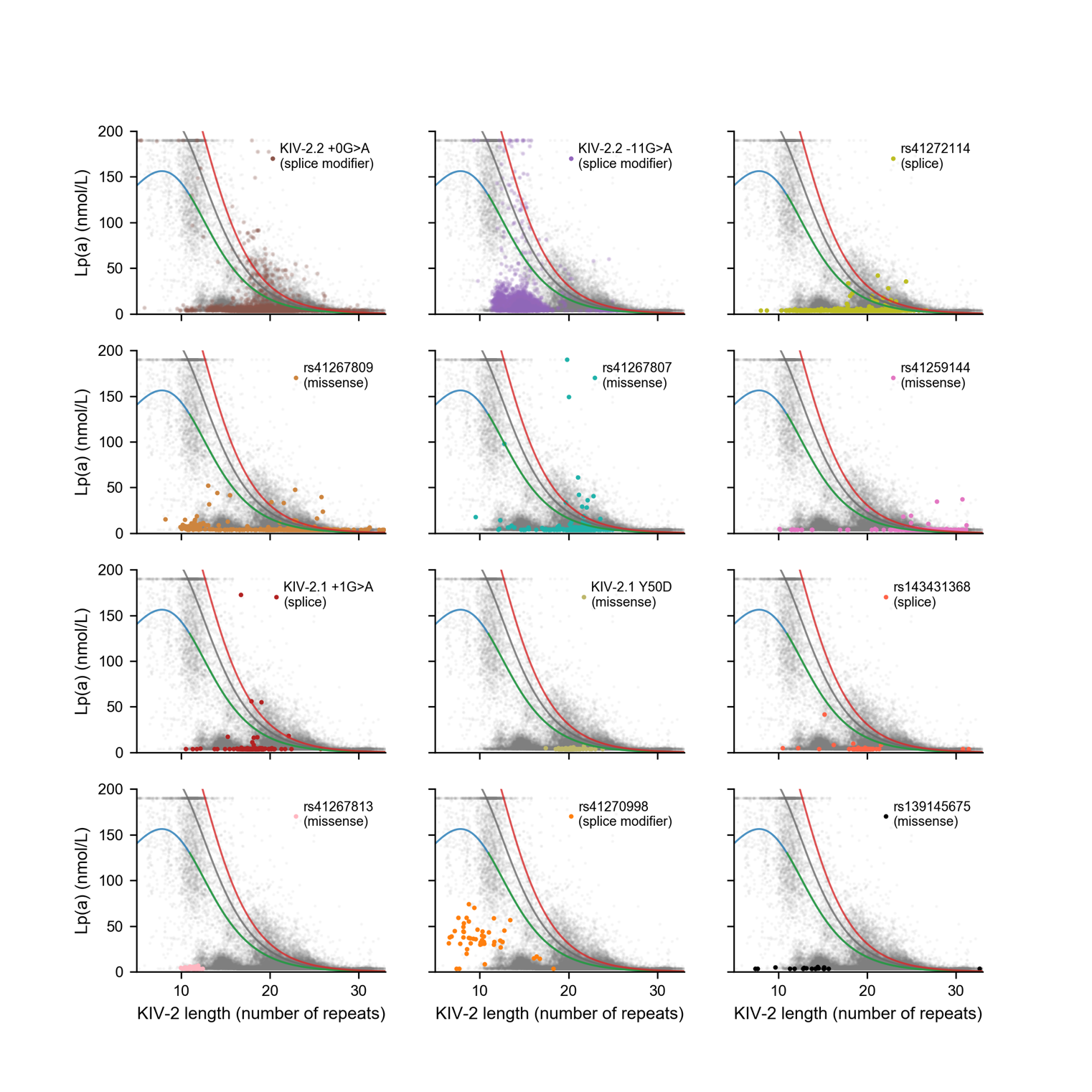

**Extended Data Figure 4. Lp(a)-reducing effects of 12 *LPA* coding or splice variants.** Scatter plots of Lp(a) vs. KIV-2 length, with carriers of a single Lp(a)-modifying variant identified in one of the first 12 steps of conditional fine-mapping analyses (Supplementary Table 3) highlighted in each panel. Points in each panel correspond to *LPA* alleles in *N*=24,969 exome-sequenced UKB participants of European ancestry for which the allele on the homologous chromosome was predicted to produce little or no Lp(a) (i.e., <4 nmol/L).

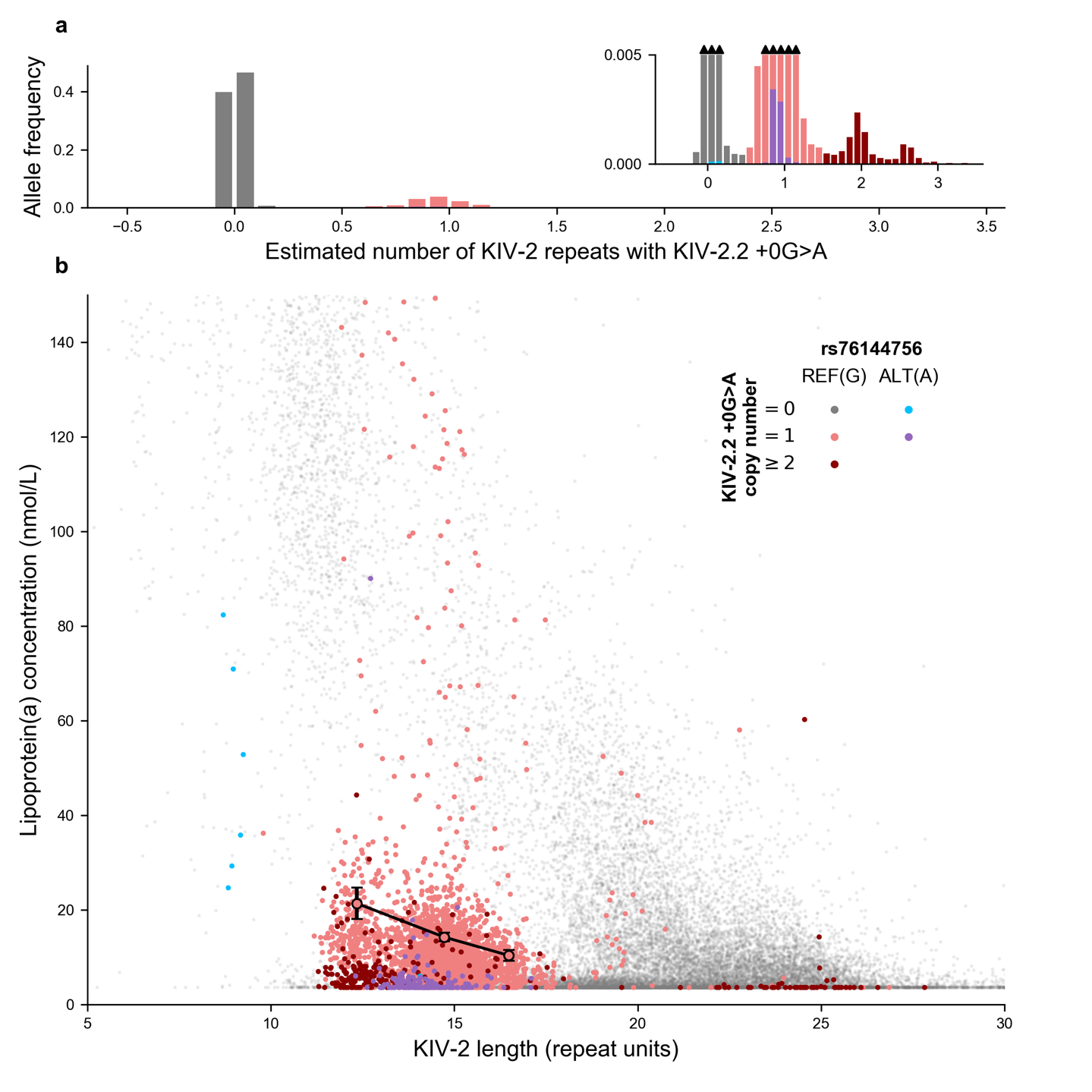

**Extended Data Figure 5. Epistatic effects of splice modifier variant**

**KIV-2.2 +0G>A (G4925A) dosage, KIV-2 length, and *LPA* missense SNP rs76144756 on Lp(a). a,** Distribution (across *N*=68,836 European *LPA* alleles) of the estimated number of KIV-2 repeats carrying a common splice variant in the last base pair of KIV-2 exon 2 that impairs splicing at the adjacent splice junction (G4925A in the nomenclature of Coassin et al. 2017)**. b,** Lp(a) vs. KIV-2 length for alleles in a near-haploid model as in Fig. 2a, with color indicating KIV-2.2 +0G>A dosage and rs76144756 status. Alleles carrying a single copy of the +0G>A splice variant exhibit substantially reduced Lp(a); alleles that additionally carry either another copy of the splice variant (on an additional KIV-2 repeat within the same allele) or the missense SNP rs76144756 exhibit further reduction in Lp(a). The inverse relationship between KIV-2 length and Lp(a) is still evident among alleles with impaired splicing due to carrying one copy of the KIV-2.2 +0G>A splice variant: large markers indicate mean Lp(a) among such alleles carrying the reference allele of rs76144756, binned by KIV-2 length (error bars, 95% CIs).

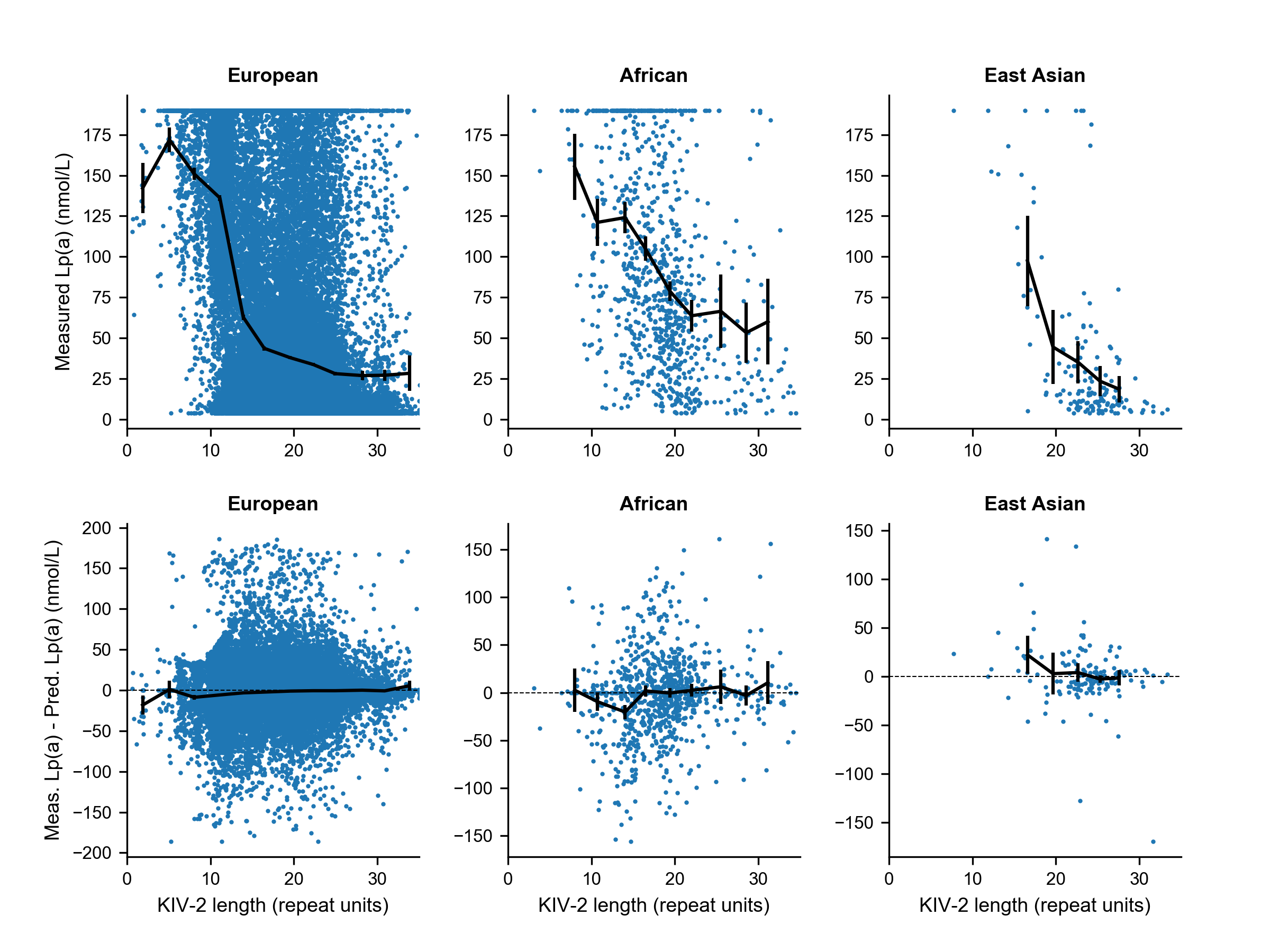

**Extended Data Figure 6. Allele frequency differences in KIV-2 VNTR length and *LPA* coding and 5’ UTR SNPs explain cross-population differences in the relationship between KIV-2 length and Lp(a). Top,** Measured Lp(a) vs. KIV-2 allele length (of one haplotype) in exome-sequenced UKB participants of self-reported European ancestry (left; *N*=46,472), African ancestry (middle; *N*=1,008), and East Asian ancestry (right; *N*=173). Mean Lp(a) across KIV-2 length bins is shown in black (error bars, 95% CIs). **Bottom**, Same as in **Top** but with measured Lp(a) residualized by genetically predicted Lp(a). The genetic architecture of Lp(a) appears to be consistent across populations, with predictions (based on a model we fit using European-ancestry samples; Methods) accounting for the KIV-2 length association in all populations.

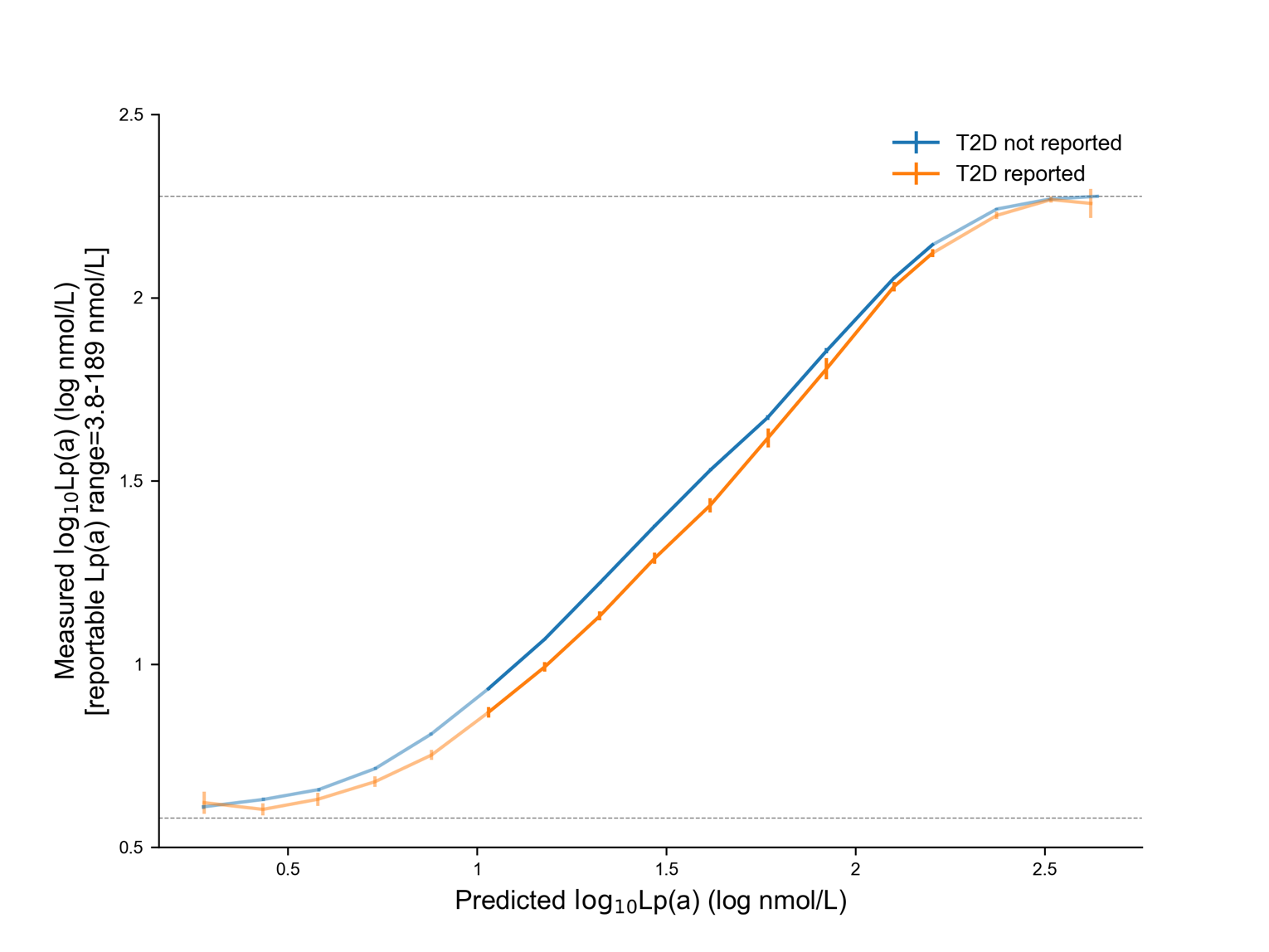

**Extended Data Figure 7. Type 2 diabetes patients exhibit reduced Lp(a) across the range of genetically predicted Lp(a).** Log-log plot of measured Lp(a) vs. genetically predicted Lp(a), stratified by type 2 diabetes (T2D) status, in *N*= 311,854 unrelated, British UKB participants. Means are plotted for bins of predicted Lp(a); error bars, 95% CIs. The reportable range of Lp(a) measurements was 3.8-189 nmol/L (horizontal lines), such that the cropping of reported measurements to this range tended to caused bias toward the mean in Lp(a) measurements of individuals with very low or very high Lp(a).

| **Repeat type** | **Repeat unit nucleotide sequence** | **Amino acid sequence** |
| --- | --- | --- |
| 1 | GGGCTTCCTTCTGGAGAAGTTCTAGAGACCACTGCCCCTGGAGTAGAGGACATCAGC | GLPSGEVLETTAPGVEDIS |
| 2 | GGGCTTCCTTCTGGAGAAGTTCTAGAGACCGCTGCCCCTGGAGTAGAGGACATCAGC | GLPSGEVLETAAPGVEDIS |
| 3 | GGGCTTCCTTCTGGAGAAGTTCTAGAGACTGCTGCCCCTGGAGTAGAGGACATCAGC | GLPSGEVLETAAPGVEDIS |
| 4 00x  4 01x  4 1x0  4 1x1 | GGGCTTCCTTCTGGAGAAGTTCTAGAGACTACTGCCCCTGGAGTAGAGGACATCAGC  GGGCTTCCTTCTGGAGAAGTTCTAGAGACTACTGCCCCTGGAGTAGAGGAGATCAGC  GGGCTTCCTTCTGGAGAAGTTCTAGAGACTACTGCCCCTGGAGTAGATGAGATCAGC  GGGCTTCCTTCTGGAGAAGTTCTAGAGACTACTGCCCCTGGAGTAGATGAGATCAGT | GLPSGEVLETTAPGVEDIS GLPSGEVLETTAPGVEEIS GLPSGEVLETTAPGVDEIS GLPSGEVLETTAPGVDEIS |
| E | GGGCTTCCTTCTGGAGAAGTTCTAGAGACTTCTACCTCTGCGGTAGGGGACCTCAGT | GLPSGEVLETSTSAVGDLS |
|  | **** * * ***  2 variants in 3 variants  “diagnostic codons” distinguish  -> types 1-4 repeat 4 types |  |

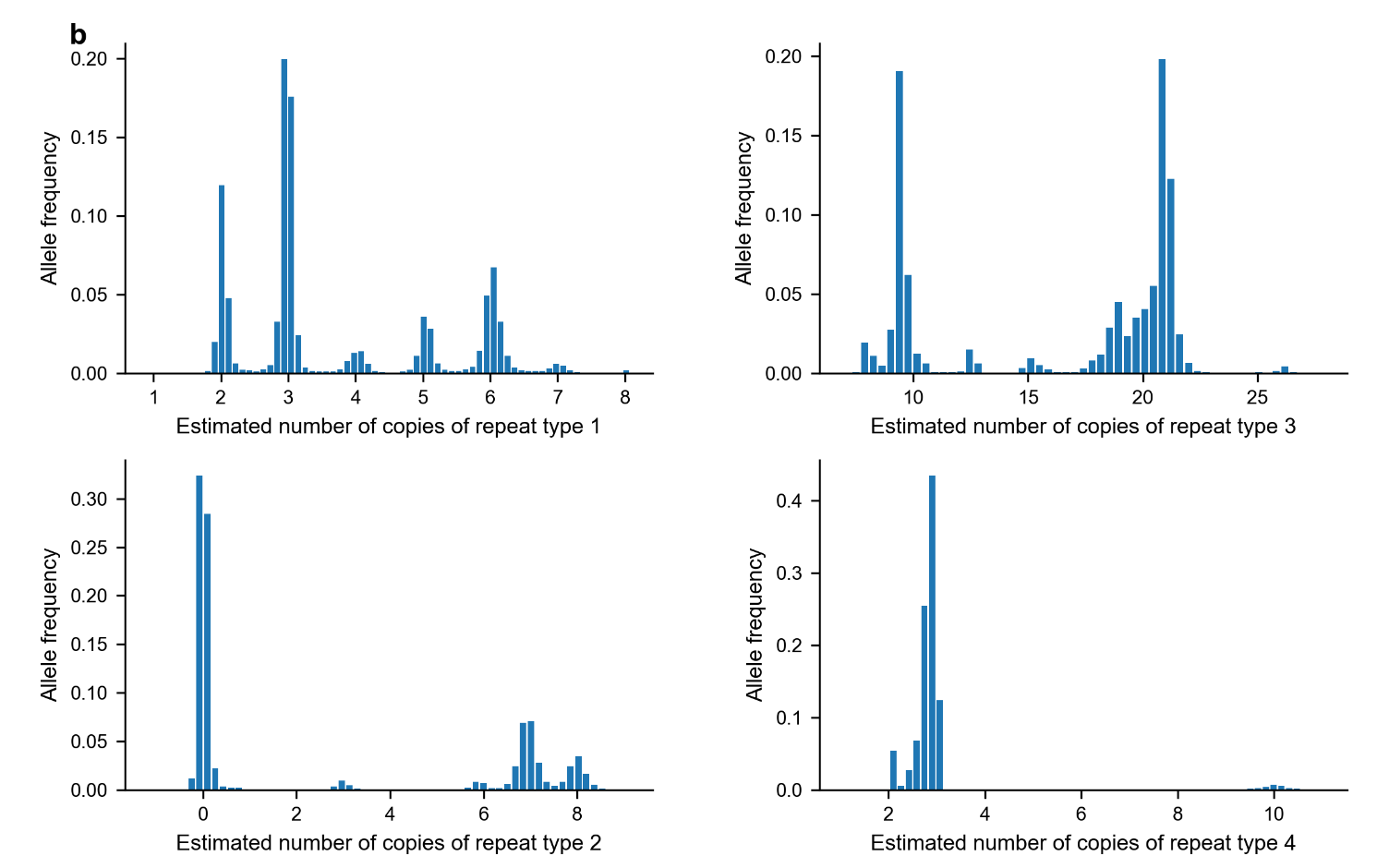

**Extended Data Figure 8. Genotyping *ACAN* VNTR repeat subtypes. a**, Repeats within the *ACAN* VNTR can be broadly classified into four repeat types according to sequence variation at two consecutive “diagnostic” base pairs (Doege et al. 1997). The fourth repeat type can be further subdivided based on three additional paralogous sequence variants (PSVs). The final, partially-diverged repeat unit (“repeat E”) of the *ACAN* VNTR is lost in a few rare alleles, creating unique sequence that we identified directly from exome sequencing reads. **b**, Separately genotyping and phasing each repeat type based on counts of sequencing reads containing the corresponding PSVs (Methods) enabled accurate, integer-mode estimates of repeat type counts within *ACAN* VNTR alleles. Allele distributions among *N*=46,472 exome-sequenced UKB participants of European ancestry are shown.

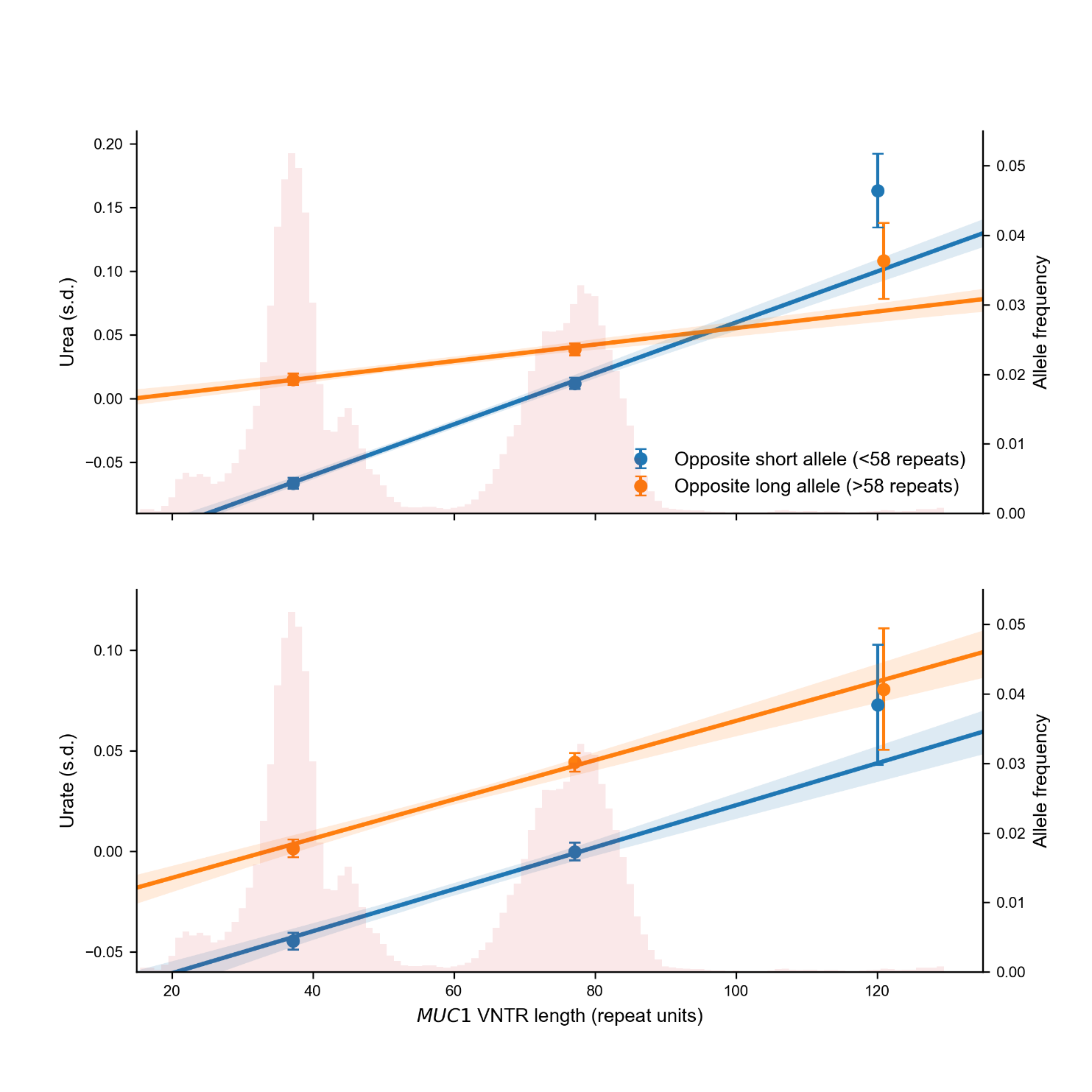

**Extended Data Figure 9. Incomplete dominance in the association of *MUC1* VNTR length with serum urea. a,** Serum urea vs. *MUC1* VNTR length, stratified by length of the VNTR allele on the homologous chromosome, in *N*=415,280 unrelated UKB participants of European ancestry. Plot markers indicate mean urea for alleles binned by *MUC1* VNTR length; least-squares linear fit (solid) also shown. **b,** Serum urate vs. *MUC1* VNTR length, stratified and plotted as in panel **a**. Error bars, 95% CIs. We formally tested for a dominance effect using linear regression in which we included an interaction term (the product of maternally- and paternally-derived allele lengths) in addition to summed allele lengths, age, age squared, sex, and 20 PCs, obtaining *P*=2.3 x 10^-20^ for urea and *P*=0.56 for urate.
