## Supplementary Note for "Protein-coding repeat polymorphisms strongly shape diverse human phenotypes"

**Supplementary Information for “Protein-coding repeat polymorphisms strongly shape diverse human phenotypes”**

### 1. Identifying protein-coding variable number tandem repeats

The analysis pipeline we used to identify protein-coding VNTRs (outlined in Methods) consisted of (i) scanning the GRCh38 reference genome for exon-overlapping repeat sequences; (ii) identifying the subset of such regions that appeared to exhibit heritable length variation (as estimated by exome sequencing read-depth); and (iii) applying a series of filters to reach a final, high-confidence set of VNTRs likely to affect protein length. The primary repeat ascertainment pipeline we used for (i) (based on Tandem Repeats Finder^1^) is described in Methods, as is the read-depth-based length estimation procedure we used for (ii). In this note we describe an additional method we developed to identify larger VNTRs (augmenting results from Tandem Repeats Finder) in (i), and we detail the filtering procedure we used for (iii).

**Scanning the genome for longer repeat regions**

To find larger and more highly-diverged candidate VNTR loci than those identified by Tandem Repeats Finder, we developed an additional, more permissive method to scan the reference genome for repeat regions that were likely to be enriched for polymorphic VNTR loci. This approach was motivated by our observation that known polymorphic VNTRs often correspond to sequences in the reference genome that are in tandem repeated segments but sometimes with high divergence. The method is comprised of two components: “VNTR Scanner” and “VNTR Partitioner.”

*VNTR Scanner.*

Step 1: Alignment scanning

Given the reference genome (GRCh38) and a fixed kmer size (k=30) and maximum genomic distance (maxDistance=1Mb), we find all kmer alignments based on the following criteria.

A kmer alignment is a list of L kmers where:

a) each of the L kmers occurs in the reference genome T times (exact matches)
b) all of the T alignments for this kmer occur within the same genomic window with maximum size maxDistance
c) the alignments of each kmer in the list correspond to consecutive positions along the genome in each of the T occurrences and
d) T >= minAlignments (=3) and T <= maxAlignments (=1000)

To make such scanning efficient, we utilized a pre-computed index of the reference genome based on the Burrows-Wheeler Transform as implemented in the bwa aligner^2^.

Step 2: Alignment grouping

Given the set of kmer alignments from step 1, we compute a set of (often larger) genome segments that are enriched for containing such kmer alignments to evaluate as candidate VNTRs.

For each kmer alignment, we define the encompassing interval as the maximum extent of the union of the kmer alignment locations. Each base within the encompassing interval is either aligned (i.e. lies within an aligned kmer) or not. Each candidate segment S is defined by a set of kmer alignments and the encompassing interval of S is the union of the encompassing intervals of the kmer alignments. We define the (alignment) density of a segment S as the number of aligned positions in the kmer alignments of S divided by the length of the encompassing interval of S.

To group alignments into segments, we first form an initial set of candidate segments by merging together all kmer alignments whose encompassing intervals transitively overlap. Given a density threshold minDensity (=0.75), we recursively subdivide each candidate segment S until the density of S is at or above minDensity. Subdividing a segment is performed by finding the largest gap (largest contiguous run of non-aligned positions) within the encompassing interval of S, splitting the kmer alignments of S into two subsets that are (fully) on the left or right of the gap, and then regrouping the kmer alignments in each subset into smaller candidate segments (again by transitive overlap of encompassing intervals).

*VNTR Partitioner.*

Given a genomic segment (e.g. from VNTR Scanner), VNTR Partitioner uses heuristic methods to analyze the segment for repeating VNTR-like self-alignments. The partitioner estimates an approximate "period" of the repetitive structure of the segment (i.e. the length of the repeating unit), then estimates a set of breaks at which to partition the input segment based on the period and finally creates a multiple sequence alignment of these subsegments of the original segment, from which we derive the consensus repeat unit and final period estimate. (We used these estimates only to perform initial filtering on candidate VNTRs and later re-examined VNTR repeat units and periods manually for VNTRs of particular interest; as such, we only briefly outline the main ideas of this method below.)

The partitioner uses multiple strategies to estimate the period of the segment.

The first is an approach similar to the VNTR Scanner. Given a kmer size (k=30) a set of kmer alignments are computed within the input segment. Each base position in the segment is assigned the period of the kmer alignment containing the base and the estimated period is the mode of the period distribution across all bases in the segment.

The second strategy aligns the input segment against a linearly-shifted copy of itself scoring each position as either +1 for a match or -1 for a mismatch. The shift may be up to 75% of the length of the segment, although shifts above 50% of the length of the segment are penalized by a score of -0.5 for each base over 50%. The estimated period is the smallest shift that maximizes this alignment score.

The third strategy takes the estimated period from method 1 or method 2 (whichever is larger, but capped at 200bp) and extracts a "query" sequence of this length from the start of the input segment. This query sequence is then aligned to the input segment in sliding target windows using the Smith-Waterman-Gotoh algorithm (with parameters match=2, mismatch=1, open=2.5, extend=0.5). Each target window is padded on the left and the right by 30% of the estimated period or 20bp, whichever is less. For each target window, the best SW alignment and alignment score is obtained as well as the predicted start of this query sequence alignment within the full input segment. Consecutive target windows will often have the same start position for the alignment and often the same alignment score.

Alignment start positions are then computed that represent local maxima in the vector of the alignment scores from each target window. Several heuristics are used to reduce noise in the estimation of the local maxima: Local maxima are computed over a small window with width equal to 20bp or the estimated period, whichever is less. To be selected as a local maximum, the alignment score for this position must be within a certain fraction (50%) of the global maximum alignment score. If potential local maxima have equal alignment scores, the alignment position corresponding to the largest number of target windows is used. Local maxima must be at least 3bp apart.

After the local maxima are determined, they are used to re-estimate the period (by taking the mode) and the sliding window alignment procedure is iterated one additional time with the updated period estimate.

Finally, a multiple-sequence alignment is constructed by extracting segments bounded by the local maxima of the sliding window alignments and aligning them using MUSCLE^3^ v3.8.425 with default parameters. From the MUSCLE alignment, a consensus repeat sequence is calculated based on the most common value in each column of the alignment and a final period estimate is determined based on the length of the consensus repeat sequence.

**Filtering candidate VNTR loci to a high-confidence set of protein-coding VNTRs**

Our analyses of the GRCh38 reference sequence identified 8,186 potential autosomal, exon-overlapping VNTRs (8,048 from Tandem Repeats Finder and 138 from the method above) that also overlapped the exome-capture targets used by UK Biobank. However, we expected that the large majority of these tandem repeats did not represent true protein-coding VNTRs (either because they did not vary in length in the population or because they did not overlap coding sequence due to imprecise endpoint-calling) or were too short to accurately genotype from sequencing depth-of-coverage data. We therefore developed a stringent filtering pipeline to create a high-confidence subset of protein-coding VNTRs for downstream analysis, ensuring that estimated lengths of these VNTRs (based on WES depth-of-coverage) exhibited heritable variation that was not the result of being contained within a larger CNV. This filtering pipeline proceeded in two steps.

*Step 1: Initial filters.*

- Removed regions with short repeat units (<9bp).
- Removed regions with fewer than 3 repeat units in the GRCh38 reference.
- Removed regions that did not appear to exhibit heritable length variation (as estimated by exome-sequencing depth-of-coverage), i.e., IBD2 sibling correlation ($IBD2 R$) < 0.25 or estimated imputation accuracy $\hat{R^{2}}<0.25$ (see later Supplementary Note sections for detailed definitions).
- Removed duplicated regions, choosing randomly among pairs of regions whose endpoints were within 100bp of one another.
- Removed regions whose flanking sequences had paralogs elsewhere in the genome (indicating potential containment in a larger CNV). To identify matches, we ran BLAT^4^ v35 on the 500bp sequence 50bp upstream and downstream of the putative VNTR, filtering it if a >97%-identity match was found for either flanking sequence.
- Removed regions that were not contained within a single gene transcript.

These initial filters produced a list of 254 exon-overlapping potential VNTR regions in 190 genes (where the number of regions exceeded the number of unique genes because some VNTR regions were still represented by multiple repeat sequence calls with different boundaries).

*Step 2: Stringent filters.*

- Required regions to either contain at least three entire exons, or be spanned by a single exon (of any transcript, allowing the region’s end points to move by up to 5bp to satisfy this requirement).
- Eliminated pairs of non-overlapping regions whose length estimates (imputed into the full UKB cohort) were correlated (*R*^2^>0.5) (indicating potential containment in a larger CNV).
- Eliminated regions whose depth-of-coverage length estimates in 30x-coverage WGS data (on 3,202 samples from the 1000 Genomes Project^5^) were correlated with depth-of-coverage on either flanking region (*R*>0.4), indicating potential containment in a larger CNV. (These data were generated at the New York Genome Center with funds provided by NHGRI Grant 3UM1HG008901-03S1.) Depth of coverage was measured within the putative VNTR region and in 1kb flanking segments starting 100bp outside of the VNTR region using Genome STRiP^6^ with default parameters but removing the default genome mask that masks out non-unique portions of the reference genome.
- Eliminated regions for which an overlapping region with correlated length estimates (*R*^2^>0.5) appeared to produce higher-quality read-depth-based length estimates (higher IBD2 sibling correlation).
- Eliminated regions entirely contained within another region.
- Eliminated regions that did not overlap canonical coding DNA sequence (CCDS from the UCSC hg38 downloads database).

These stringent filters eliminated roughly half of the 254 potential coding VNTRs that passed our initial filters. We then manually reviewed the excluded VNTRs, adding back in a few VNTRs that appeared to have been over-conservatively filtered (based on known VNTR variation at these loci from previous literature or examination of VNTR sequence together with transcript annotations on the UCSC Genome Browser^7^). We also manually reviewed VNTRs that passed the above filters but had estimated period lengths that were not multiples of 3, in most cases determining that the VNTR either fully spanned one or more exons or that its period had been incorrectly estimated; in other cases, we excluded the VNTR. Finally, we manually examined two remaining instances in which multiple VNTR regions were called in the same gene, selecting a representative VNTR in each case. These final filters produced our main analysis set containing 118 high-confidence protein-coding VNTRs (in 118 distinct genes).

### 2. Estimating diploid VNTR content from exome sequencing read depth

Depth-of-coverage is commonly used to identify copy-number variation from short-read sequencing^8,9^ and has more recently been used to estimate VNTR allele lengths (combined across maternally- and paternally-inherited haplotypes) from whole-genome sequencing data^6,10,11^. Here we applied this framework to estimate VNTR lengths from 76bp paired-end exome-sequencing reads (aligned to the GRCh38 reference using SPB pipeline used to process the 49,959-sample UK Biobank exome sequencing release^12^). For each repeat region ascertained in the GRCh38 reference that passed our filters for analysis (Methods), we began by counting, for each sample, the number of unique reads with a SAM flag^13^ of 0x53, 0x63, 0x93, 0xA3, 0x51, 0x61, 0x91, or 0xA1 that aligned to the region (allowing reads with any mapping quality including zero). We considered two possible definitions of “aligning to the region”: (i) having at least one mapped base fall within the region, or (ii) having all mapped bases fall within the region, choosing whichever approach produced higher IBD2 sib-pair correlation in a pilot analysis.

Counts of reads generated from polymorphic repeat regions are expected to scale linearly with total (maternal plus paternal) repeat length and with sequencing coverage, such that naively, dividing read counts by overall sequencing coverage (per-sample) produces estimates of “diploid VNTR content” (up to a suitable affine transformation). However, read counts can be strongly influenced by technical biases that vary across samples and typically contribute much more variance than Poisson noise. To mitigate this problem, computational methods that estimate depth-of-coverage usually model and correct for the effects of genomic features such as GC content that tend to influence technical biases^8,9^.

**Normalizing read depth against samples with similar exome-wide coverage profiles**

We developed a different normalization approach that leveraged the very large number of exome sequences (49,959) available for analysis: instead of explicitly identifying features that correlated with biases in sequencing depth, we identified samples that exhibited similar exome-wide coverage profiles, and we normalized VNTR-aligned read counts from each sample against corresponding read counts from the 300 samples with most similar coverage profiles (after adjusting all samples for overall coverage). The intuition behind this approach was that in the majority of the genome that does not vary in copy number between two individuals, depth-of-coverage is influenced primarily by technical biases, such that the coverage profile of each sample provides a “barcode” that identifies which other samples were influenced by the same technical biases (e.g., due to sharing the same batch effects).

The main challenge in effectively implementing this approach is deciding which genomic regions to include in the profiles. We found that restricting to "batch-informative regions" in which read depth was especially variable among samples was helpful. Understanding the properties of outlier regions in each stage of the analysis was also important: on one hand, we needed to prevent outliers from overly influencing the results; on the other hand, outliers could contain useful information about patterns of technical artifacts.

In detail, our normalization pipeline proceeded as follows:

- Compute read depth in 1kb windows of the genome (using mosdepth^14^ v0.2.5).
- Restrict to regions with 20-100x mean read depth across samples (81,564 1kb regions).
- Exclude 1kb regions overlapping repeat regions identified by an initial run of our VNTR scanner (77,003 1kb regions left).
- Normalize each sample to the same mean read depth across remaining 1kb regions.
- Normalize each region:
  -- subtract mean across individuals
  -- divide by sqrt(mean across individuals) (proportional to Poisson standard deviation).
- Rescale all normalized read depths so that median region has std dev = 1.
- Restrict to regions with top-10% variance (the intuition being that most regions just have variation due to random noise, but a small subset are informative of batch effects).
- Crop normalized read depths to (-2, 2) to reduce the effect of outliers.
- Compute pairwise distances among 49,959 samples (where distance is defined as the sum of squared differences of final transformed read depths at remaining 1kb regions).
- For each VNTR, normalize (aligned read count / exome-wide depth) in each sample by dividing this value by the mean (aligned read count / exome-wide depth) across the 300 other samples with closest-matching read-depth profiles.

We estimated (based on benchmarks assessing correlation of VNTR length estimates across IBD2 sib-pairs) that this normalization procedure tended to reduce noise variance by slightly more than half compared to naïve estimates (aligned read count / exome-wide depth) that did not make any attempt to correct for technical biases.

We implemented a few refinements of this pipeline for our final analyses of the five VNTRs that appeared to drive strong phenotypic associations. First, instead of normalizing each sample against 300 closest-matching samples chosen from all 49,958 other samples, we normalized against 200 closets-matching samples chosen from the subset of other samples who reported European ancestry (to prevent ancestry bias among the set of “reference samples” chosen for normalization). Second, for the *LPA* and *ACAN* VNTRs, we leveraged variation within these VNTRs to separately estimate length variation of specific subtypes of repeats (by filtering to reads distinguishing the subtypes). Third, we rescaled allele length estimates (which the above pipeline normalizes to a mean of ~1 across all diploid estimates) to absolute numbers of repeats per allele (by comparison to literature and/or identification of discrete peaks in phased allele distributions). These latter two refinements are detailed for each VNTR (see subsequent Supplementary Note).

### 3. Phasing and imputing VNTR lengths using surrounding SNPs

Our read-depth-based VNTR length estimates (“diploid VNTR content”) for 49,959 exome-sequenced UK Biobank participants enabled haplotype-sharing analyses to phase (and simultaneously refine) VNTR length estimates within the exome-sequenced cohort and to impute allele lengths into the remainder of the UK Biobank cohort. We employed an iterative algorithm in which haploid allele lengths of each individual in turn were updated according to a probabilistic haplotype-copying model using all other haplotypes as a reference panel, prioritizing copying from haplotypes closely matching the individual’s SNP-haplotypes. This overall structure is similar in spirit to the approach applied by PHASE v2^15^ and many subsequent algorithms developed to phase biallelic SNPs; however, we adapted the approach in several ways to better suit the task of phasing a single multiallelic VNTR variant (with real-valued length estimates) in a very large cohort for which accurately-phased haplotypes of surrounding SNPs had previously been generated^16^. (We performed phasing analyses on the subset of 49,796 exome-sequenced individuals with phased SNP-array haplotypes.)

**Haplotype-copying model based on identity-by-state (IBS) sharing**

Most of the commonly-used statistical phasing algorithms that have been developed to date have aimed to phase many SNPs across a locus or a chromosome. In this setting, hidden Markov model (HMM)-based approaches that model haplotypes as mosaics of sub-haplotypes copied (imperfectly) from a reference panel of other haplotypes (e.g., according to the Li-Stephens model^17^) have been a natural and very effective approach, as HMMs facilitate efficient computation of posterior probabilities across many SNPs via dynamic programming algorithms.

Here, we faced a very different phasing task: at a typical locus, we only needed to phase a single VNTR and could assume that SNP-haplotypes had already been phased to chromosome-scale accuracy^16^. As such, while a standard Li-Stephens HMM would produce a reasonable haplotype-copying model that could be used within a phasing routine, the HMM framework was unnecessary here, and a much simpler haplotype-copying model based on lengths of identity-by-state (IBS) sharing could produce equivalent results with less computation. Moreover, the IBS-based approach facilitated rapid tuning of the parameters of the haplotype-copying model, ultimately enabling improved phasing accuracy.

The typical length of a genomic segment shared identical-by-descent (IBD) between two haplotypes scales inversely with the time to the most recent common ancestor (TMRCA) because the TMRCA determines the number of opportunities for recombination in the coalescent tree connecting the haplotypes. In general, IBD segments are not expected to be identical-by-state (IBS)—i.e., carry exactly the same alleles for all SNPs within—due to gene conversion, recent mutation, and genotyping and phasing error. Conversely, IBS-sharing may extend beyond the ends of IBD segments due to sharing of common haplotypes. However, when considering only IBS across accurately-genotyped, accurately-phased SNPs from a modern SNP-array platform (which is unlikely to “notice” most gene conversions or recent mutations), the biological and technical complexities that break up IBS-sharing within an IBD segment tend to be quite rare (i.e., occur only once every few megabases), and portions of longer IBS segments not too near their ends tend to consistently represent true IBD.

We therefore quantified lengths of haplotype-sharing (in genetic map coordinates, i.e., centimorgans) with an IBS-like measure designed to be more robust to these complexities:

$$\mathcal{l}_{\mathrm{dir}}=a\cdot\left( IBS length in direction \left( left/right \right) \right)+\left( 1-a \right)\cdot\left( IBS length in direction, allowing 1 error \right)$$

$\mathcal{l=}b\cdot\left( \mathcal{l}_{\mathrm{left}}+ \mathcal{l}_{\mathrm{right}} \right)+\left( 1-b \right)\cdot\min\left( \mathcal{l}_{\mathrm{left}},\mathcal{l}_{\mathrm{right}} \right)$.

We set $a=0.5$ and $b=0.25$ based on empirical performance in phasing benchmarks (which tended to be insensitive to the choice of these parameters) and found that IBS-like lengths defined in this way served as a reasonable proxy for IBD lengths for the purpose of identifying haplotypes likely to share a recent common ancestor (and carry the same allele of a VNTR).

We then defined a haplotype-copying model in which the probability of a given haplotype having been copied from each of a set of reference haplotypes scaled roughly with:

$$exp(-\mathcal{l}_{0}\mathcal{/l)}$$

where $\mathcal{l}_{0}$ denotes a constant factor (to be tuned from the data) and $\mathcal{l}$ denotes the IBS-like length computed above. For computational convenience, we restricted the set of potential reference haplotypes to the top $K_{\mathrm{top}}$ haplotypes with longest $\mathcal{l}$, and to add more flexibility to the functional form of the relationship between haplotype-sharing length and copying probability, we adjusted copying probabilities by including a regularization probability $p_{\mathrm{reg}}$ of randomly sampling from the top $K_{\mathrm{top}}$ longest-matching reference haplotypes independent of $\mathcal{l}$. Together, these features produced copying probabilities:

$$P\left( copy from ref. hap. i \right)=\frac{\exp\left( -\frac{\mathcal{l}_{0}}{\mathcal{l}_{i}} \right)}{\sum_{k \mathrm{in} K_{\mathrm{top}}} \exp\left( -\frac{\mathcal{l}_{0}}{\mathcal{l}_{k}} \right)}\cdot\left( 1-p_{\mathrm{reg}} \right) + \frac{p_{\mathrm{reg}}}{K_{\mathrm{top}}}$$

We tuned the parameters $\mathcal{l}_{0}$, $K_{\mathrm{top}}$, and $p_{\mathrm{reg}}$ independently for each VNTR as described at the end of this section.

This haplotype-copying model immediately provided a means to impute VNTR allele lengths from a phased panel of reference SNP+VNTR length haplotypes into a target SNP haplotype (as the weighted average of copied haplotypes, weighting according to the above probabilities) and provided a probabilistic framework within which to perform phasing.

**Model for estimated diploid VNTR content**

Assuming an individual’s two haplotypes (carrying VNTR alleles of length $x_{1}$ and $x_{2}$) had been copied from reference haplotypes $i$ and $j$ with (phased) VNTR allele lengths $x_{i}$ and $x_{j}$, we modeled the observed “diploid VNTR content” estimate obtained from read-depth as being generated under the following model:

$$x_{1}= x_{i}+\epsilon_{\mathrm{mut}}^{\left( 1 \right)}, x_{2}= x_{j}+\epsilon_{\mathrm{mut}}^{\left( 2 \right)}$$

$$diploid VNTR estimate=x_{1}+x_{2}+\epsilon_{\mathrm{err}}=\left( x_{i}+\epsilon_{\mathrm{mut}}^{\left( 1 \right)} \right)+\left( x_{j}+\epsilon_{\mathrm{mut}}^{\left( 2 \right)} \right)+\epsilon_{\mathrm{err}}$$

where each $\epsilon_{\mathrm{mut}} \sim N(0, \sigma_{\mathrm{mut}}^{2})$ roughly models “mutation” along the branch of the coalescent tree connecting the individual’s haplotype and the copied reference haplotype (but also incorporates potential error in the reference haplotype’s VNTR length) and $\epsilon_{\mathrm{err}} \sim N(0, \sigma_{\mathrm{err}}^{2})$ models noise in our read-depth-based estimate of diploid VNTR content. We also tuned $\sigma_{\mathrm{mut}}$ and $\sigma_{\mathrm{err}}$ independently for each VNTR.

**Iterative update of haploid allele length estimates**

Within this probabilistic framework, we performed phasing by iteratively leaving out each individual in turn and updating the left-out-individual’s (phased) VNTR allele lengths to their posterior means (assuming each of the individual’s two haplotypes had been “copied with noise” under the above model from a reference panel containing all other individuals’ haplotypes). Explicitly, the posterior probability of copying the individual’s haplotype 1 from reference haplotype $i$ and copying haplotype 2 from reference haplotype $j$ satisfies:

$$P\left( hap1 copied from ref. hap. i, hap2 copied from ref. hap. j \right| diploid VNTR estimate=\hat{x_{1+2}})$$

$$\propto P\left( \mathrm{SNP} hap1 from ref. hap. i \right)\cdot P\left( SNP hap2 from ref. hap. j \right)\cdot\exp\left( -\frac{\left( \hat{x_{1+2}}-x_{i}-x_{j} \right)^{2}}{2\cdot\left( 2\sigma_{\mathrm{mut}}^{2}+\sigma_{\mathrm{err}}^{2} \right)} \right)$$

which provides weights from which the posterior mean can be computed as the weighted sum of posterior means assuming each copying configuration (using the formula for the conditional expectation of a bivariate Gaussian):

$$E\left[ x_{1} \right| hap1 from ref. hap. i, hap2 from ref. hap. j, diploid VNTR estimate=\hat{x_{1+2}}]$$

$$=x_{i}+\frac{\epsilon_{\mathrm{mut}}^{2}}{2\cdot\left( 2\sigma_{\mathrm{mut}}^{2}+\sigma_{\mathrm{err}}^{2} \right)}\cdot\left( \hat{x_{1+2}}-x_{i}-x_{j} \right)$$

and likewise for the posterior mean of $x_{2}$.

We slightly modified the above computation to disallow copying both haplotypes from the same individual; this safeguard is commonly implemented in SNP-phasing methods to prevent circular updates in which two individuals who share both haplotypes IBD iteratively copy each other’s phased allele estimates, impeding progress. We implemented this modification by zeroing out probabilities of copying from reference haplotypes $i$ and $j$ in the same reference individual (and renormalizing probabilities).

**Continuous (real-valued) versus discrete allele length estimates**

Our probabilistic framework modeled VNTR allele lengths as real-valued quantities rather than integer-valued copy numbers. This framework was sensible for modeling read-depth-based (continuous) estimates of diploid VNTR content and enabled efficient computation of posterior probabilities using Gaussian conditional expectations. In downstream analyses (e.g., association tests), we continued to analyze the real-valued VNTR allele lengths produced by our phasing and imputation pipeline, as these “dosage”-type estimates appropriately reflect uncertainty in allele lengths, maximizing power in regression analyses. However, to improve clarity of presentation, we displayed allele histograms using discrete bins in our main figures (equivalent to rounding allele lengths to the nearest integer).

**Benchmarking of phasing and imputation accuracy**

To benchmark the accuracy of phased VNTR length estimates, we adapted a trio-based approach commonly used for SNP-phasing benchmarks. The basic idea behind this benchmark is that if a cohort contains several complete trios (i.e., father, mother, and child), then phasing the cohort excluding all trio children will produce allele length estimates for all parental haplotypes (without using any information from within-family haplotype-sharing). Each child’s diploid VNTR estimate can then be compared to the sum of the phased allele length estimates for the two transmitted haplotypes, and assuming independence of parental haplotypes, this correlation satisfies the relationship:

$$"\mathrm{child} R^{2}"=R^{2}\left( diploid estimates, sum of transmitted phased estimates \right)=R^{2}\left( diploid estimates, truth \right){\cdot R}^{2}(phased estimates, truth)$$

We can thus estimate phasing accuracy by dividing the quantity on the left (obtained in trio children) by the estimated accuracy (i.e., $R^{2}$ vs. truth) of the diploid VNTR content estimates we obtain from read-depth. Assuming unbiased error in diploid VNTR estimates, this latter quantity can in turn be estimated as the correlation between diploid estimates obtained from independent sequencing of monozygotic twins, or more generally, “IBD2” sib-pairs who share both haplotypes IBD (i.e., inherited the same allele from their mother and inherited the same allele from their father):

$$\hat{R^{2}}\left( diploid estimates, truth \right)=R\left( diploid estimate in sib 1, diploid estimate in sib 2 \right)="IBD2 R"$$

Putting these two relationships together, we obtain:

$$\hat{R^{2}}\left( phased estimates, truth \right)=\frac{\mathrm{child} R^{2}}{IBD2 R}$$

The exome-sequenced UK Biobank cohort that we analyzed contained 613 sib-pairs, one-quarter of which are expected to be IBD2 at a given locus. We readily identified high-confidence IBD2 sib-pairs based on at most 3 mismatching SNP-array genotypes within ±1 Mb of each VNTR (computed using plink^18^ 1.9), from which we obtained $IBD2 R$ estimates with reasonably low noise.

The cohort contained far fewer complete trios, such that we needed to adapt the above approach to instead use “pseudo-trios”: i.e., “pseudo-children” each of whose haplotypes had a long IBD match with a “surrogate parent” in the data set. We identified pseudo-trios by starting with long IBD matches among related individuals previously identified by UK Biobank^19^ (specifically requiring either >3 cM IBD between 1^st^-degree relatives or >5 cM IBD between more-distant relatives) and then identifying the best surrogate for the opposite haplotype of either member of the related pair, determining that a pseudo-trio had been completed if a surrogate with >3 cM IBD could be found. (In these computations, we approximated IBD length with the average of no-error and 1-error IBS, requiring this average to be >0.5 cM on either side of the VNTR.) This approach identified roughly 600 pseudo-trios per locus, which sufficed to obtain $"\mathrm{child}R^{2}"$estimates with low noise.

Benchmarking imputation accuracy was much simpler using the standard cross-validation approach: we left out 5% of individuals in the cohort, phased the remainder of the cohort, and then imputed into the left-out individuals, estimating imputation accuracy as:

$$\hat{R^{2}}\left( imputed estimates, truth \right)=\frac{R^{2}(imputed estimates, left out estimates)}{IBD2 R}$$

One important caveat of the above benchmarks is that they assume unbiasedness of errors in read-depth-based estimates of diploid VNTR content. In reality, exome sequencing coverage depths at VNTRs can be biased by the presence of paralogous sequence variants (PSVs) within repeat units (that can subtly affect exome capture) or by read-mapping biases for very short alleles. While these biases did not produce first-order errors in either our allele length estimates or our estimates of phasing or imputation accuracy, we also performed imputation accuracy benchmarks by comparing imputed allele lengths derived from WES-read-depth to diploid VNTR content estimated from WGS-read-depth (which was available to us for 48 UK Biobank participants included in a WGS pilot study). Our analyses of these benchmarks are described in the next supplementary note.

**Optimization of phasing and imputation parameters**

Our phasing algorithm included five constant parameters ($\mathcal{l}_{0}$, $K_{\mathrm{top}}$, $p_{\mathrm{reg}}$, $\sigma_{\mathrm{mut}}$, and $\sigma_{\mathrm{err}}$) that determined relative copying weights of reference haplotypes with different lengths of IBS-sharing and determined the relative weighting of information derived from haplotype-sharing vs. direct read-depth-based estimates. The optimal choices of these parameters are expected to differ among VNTRs depending on their mutational characteristics (e.g., mutation rate) and sequencing properties (e.g., average number of VNTR-derived fragments sequenced). We therefore incorporated parameter-tuning into our phasing algorithm to enable each VNTR to be analyzed using parameters tailored for the VNTR.

The simplest approach to tuning would be to employ a comprehensive search across the reasonable parameter space for each parameter, benchmarking phasing accuracy (as described above) for each parameter configuration and then selecting the choice of parameters that yielded the highest accuracy. However, this brute-force approach was computationally intractable given the five-dimensional parameter space and nontrivial amount of computation required to perform each phasing analysis.

To optimize parameters in a computationally tractable way, we instead employed a simulated annealing strategy^20^ in which we explored the parameter space by iteratively: (1) proposing a random step (i.e., change to a single parameter); (2) evaluating phasing accuracy after one more full iteration (leaving out and updating each individual’s allele length estimates in turn) using either the current or proposed new parameter configuration; and (3) accepting the proposed parameter change if it achieved acceptable accuracy (relative to the accuracy obtained by running the full iteration using the current parameter configuration). This stochastic search technique was intended to improve robustness to local optima.

In detail, our algorithm began by initializing the parameters to:

$\mathcal{l}_{0}=2 \mathrm{cM}$, $K_{\mathrm{top}}=50$, $p_{\mathrm{reg}}=0.05$, $\sigma_{\mathrm{err}}=std.dev.\left( diploid estimates \right)\cdot\sqrt{IBD2 R}$, $\sigma_{\mathrm{mut}}=\frac{\sigma_{\mathrm{err}}}{2}$

and initializing both allele lengths of each individual to half the diploid estimate for that individual. We then ran $T$ = 500 phasing iterations, each of which updated each individual’s phased allele lengths once, proceeding through the individuals in random order. We designated the first 20 iterations as burn-in iterations in which no parameter modification was proposed. In each subsequent iteration, we proposed updating one parameter (cycling through the five parameters in turn) by multiplying it by a random factor drawn from the exponential of a uniform distribution: $\exp\left( U\left( -c,c \right) \right)$, where $c=1-\frac{t}{T}$ at iteration $t$ of $T$ for parameters $\mathcal{l}_{0}$, $K_{\mathrm{top}}$, $p_{\mathrm{reg}}$ and $c=\frac{1}{4}\left( 1-\frac{t}{T} \right)$ for parameters $\sigma_{\mathrm{mut}}$ and $\sigma_{\mathrm{err}}$ (which had a smaller range of interest). We further cropped proposed values of $K_{\mathrm{top}}$ to stay within the interval $[25, 100]$ and proposed values of $p_{\mathrm{reg}}$ to at most 0.9. We then computed the $"\mathrm{child} R^{2}"$ benchmark after one full iteration using either the current parameter configuration or the new proposal. We always accepted the proposed step if it achieved a higher $"\mathrm{child} R^{2}"$ benchmark and always rejected if it reduced $"\mathrm{child} R^{2}"$ by >0.001; if it only slightly reduced accuracy, we probabilistically rejected it with increasing probability as iterations progressed:

$$P_{\mathrm{reject}}={min(1, 10}^{4}\cdot\left( \mathrm{child} R^{2} \mathrm{reduction} \right)\cdot max(0.1, t/T))$$

After 500 such iterations, we reverted to the “phasing-optimized” configuration of parameters and allele length estimates that achieved the highest phasing accuracy across all iterations. Using these allele length estimates, we computed “imputation-optimized” values of the parameters $\mathcal{l}_{0}$, $K_{\mathrm{top}}$, $p_{\mathrm{reg}}$ by benchmarking imputation accuracy in a grid search over these three parameters. Separately, we ran 10 final iterations using the “phasing-optimized” parameters to obtain final phased allele length estimates for all individuals in the cohort (no longer leaving any individuals out for the purpose of computing accuracy benchmarks).

To minimize the effects of stochasticity, we ran the entire algorithm described above 5 times for each VNTR using 5 different random seeds (used to randomize the update order within each iteration and to randomize parameter-update proposals) and chose the run that produced the highest estimated phasing accuracy.

### 4. Optimizing genotyping of VNTRs with potential phenotypic effects

For each VNTR for which our association analysis and fine-mapping pipeline identified a potentially-causal phenotype association, we examined the VNTR allele length estimates produced by our default genotyping and phasing approach to identify potential improvements and to calibrate allele lengths to the absolute scale of repeat copies. (As described above, our initial analysis of normalized read-depths only produced unscaled allele length estimates relative to the mean.) For most VNTRs, we found that we could modestly improve the accuracy of our allele length estimates by optimizing the VNTR boundaries used to select aligning reads, separately genotyping and phasing repeat subtypes distinguished by paralogous sequence variants (PSVs) within the VNTR, or directly genotyping very short alleles using spanning reads. These improvements reduced both noise and bias in our estimates.

While the overall changes in allele length estimates between our initial genotyping (used in the initial analyses reported in Supplementary Tables 1 and 2) and optimized genotyping (used elsewhere in the paper) were modest ($R^{2}(initial, optimized)$ = 0.97 (*LPA*), 0.88 (*ACAN*), 0.93 (*TENT5A*), 1.00 (*MUC1*), 0.82 (*TCHH*)), these optimizations were important to ensure the accuracy of our deeper fine-mapping analyses identifying nearby SNPs likely to also exert causal effects on phenotypes.

***LPA***

*Sequence variation within the LPA KIV-2 VNTR.*

*LPA* alleles typically contain 10-30 repeat units of the KIV-2 VNTR, and different repeats within an allele often harbor sequence variation. In particular, the first exon (“exon 1”) of the KIV-2 repeat carries three common synonymous variants that distinguish three frequently-observed repeat types labeled A, B, and C in the literature^21^ (Extended Data Fig. 2a). Repeat type A is the most common type and accounts for 5 of the 6 KIV-2 repeats that appear in the GRCh38 reference sequence; repeat type B is also common and accounts for 1 of the 6 KIV-2 repeats in the GRCh38 reference (cf. Fig. 2 of Schmidt et al.^21^); while repeat type C is rare, accounting for only ~1% of all repeats^22^.

*Bias in IDT xGen v1 exome capture of KIV-2 repeat types.*

This common variation within the coding region of the KIV-2 VNTR created a challenge for accurate estimation of VNTR lengths from exome sequencing read counts that was compounded by the design of the IDT xGen v1 exome capture panel used to perform exome-sequencing of UK Biobank samples. The xGen v1 panel did not contain any probes targeting any of the 6 KIV-2 repeats in the GRCh38 reference (presumably because this region had been filtered as repetitive). However, it did contain probes targeting each exon of the adjacent KIV-3 repeat, which share moderate-to-high sequence similarity with KIV-2 exons 1 and 2. Specifically, exon 1 of KIV-3 is identical to the type-B exon 1 (“1B”) of KIV-2, while exon 2 of KIV-3 is similar to but somewhat diverged from exon 2 of KIV-2.

This capture design had two main effects on counts of read alignments in the KIV-2 region:

1. Reads aligning to exon 1B repeats in the GRCh38 reference (namely, the single KIV-2 repeat of type B which exactly matches exon 1 of KIV-3) represented “on-target” capture by the pair of 120bp probes that had been designed for exon 1 of KIV-3. In contrast, reads aligning to exon 1A repeats in the GRCh38 reference (i.e., the five KIV-2 repeats of type A) represented “off-target” capture, which resulted in generation of fewer reads per exon 1A unit within an allele than per exon 1B unit within the same allele.
2. Reads aligning to KIV-2 exon 2 repeats in the GRCh38 reference (namely, the six exon 2 repeats of KIV-2 as well as the exactly-matching exon 2 of KIV-1) all represented “off-target” capture from probes that had been intended to capture the somewhat-diverged exon 2 of KIV-3. Consequently, KIV-2 exon 2 read counts were unbiased with respect to repeat types but were greatly diminished (and thus noisier) due to reduced off-target capture efficiency.

Our initial analysis pipeline had simply counted reads mapping to the full repeat region (i.e., KIV-1 exon 2 + six repeats of KIV-2 + KIV-3 exon 1) and thus incurred downward bias of allele length estimates for alleles with larger fractions of type-A repeat units (due to the less-efficient, off-target capture of exon 1A).

*Correction of capture bias by separate estimation of type-A and type-B repeat copy numbers.*

To circumvent the biased capture of type-A vs. type-B repeats, we devised a modified genotyping strategy that independently estimated the copy number of each repeat type within each individual (and after phasing, within each allele). Specifically, we considered only read pairs that mapped to the seven repeats of exon 1 and its intronic flanks (i.e., the five exon 1A repeats in the KIV-2 region of the GRCh38 reference and the two exon 1B repeats, one in KIV-2 and one in KIV-3) and subdivided these read pairs into (i) those that mapped unambiguously to the 1A sequence (restricting to bases with quality ≥25); (ii) those that mapped unambiguously to the 1B sequence; and (iii) those that mapped equally well to 1A and 1B. We then performed our read-depth normalization procedure on (i) and (ii) separately (not counting reads in category (iii) or reads aligning to exon 2).

This strategy avoided the problem of capture bias because counts of reads generated from 1A vs. 1B sequence were no longer being compared to one another. As such, “1A read-depths” scaled linearly with KIV-2 type-A content and “1B read-depths” scaled linearly with KIV-2 type-B content, albeit with different constant factors due to capture bias.

*Calibrating type-A and type-B read-depths to estimate absolute KIV-2 copy numbers.*

Our modified genotyping procedure left one remaining task: determining the appropriate scaling factors to convert normalized 1A and 1B read-depths (that our pipeline normalized to a mean of 1 per diploid individual) to absolute estimates of repeat counts of types A and B and ultimately KIV-2. We accomplished this task by making use of the following observations:

- Absolute calibration of 1B read-depth. After phasing, the distribution of phase-resolved 1B read-depths exhibited discrete peaks with constant spacing, indicating that the appropriate scaling factor (~5.2) to use to obtain absolute type-B repeat counts could simply be selected to make the peaks become integer-valued (Extended Data Fig. 2b).
- Relative calibration of 1A vs. 1B read-depth. Phased 1A read-depths were insufficiently precise to resolve integer-spaced peaks (Extended Data Fig. 2b), so we instead leveraged reads aligning to exon 2 (which were equally affected by off-target capture across all KIV-2 repeat types) to calibrate 1A and 1B read-depth against one another (by regressing exon 2 read-depth on 1A and 1B read-depth). This computation yielded an estimated ratio of ~6.7 for the contribution of 1A normalized read-depth vs. 1B normalized read-depth (which corroborated with estimates of 1A vs. 1B copy number from whole-genome sequencing of 48 UK Biobank participants included in a WGS pilot study); combining this information with the 1B scaling factor of ~5.2 produced a scaling factor of ~34.9 to convert 1A read-depth to type-A repeat counts.
- Subtraction of 1 repeat corresponding to KIV-1 exon 2 + KIV-3 exon 1. Finally, we subtracted 1 from the total type-A + type-B count to adjust for our read counts including one neighboring exon on either side of KIV-2 (KIV-1 exon 2 = KIV-2 exon 2, and KIV-3 exon 1 = KIV-2 exon 1B).

Combining the above gave the calibration formula:

$$\mathrm{KIV}\text{-}2 copy number= 34.9\times\left( 1A phased read\text{-}\mathrm{depth} \right)+5.2\times\left( 1B phased read\text{-}\mathrm{depth} \right)-1$$

for obtaining absolute KIV-2 VNTR haploid copy number estimates from phased, normalized 1A and 1B read-depths. We verified that the absolute KIV-2 copy number distribution we obtained in this way was in close agreement to literature (cf. Fig. 2 of Kraft et al.^23^; note that this figure counts the total number of KIV repeats, which is 9 more than the number of KIV-2 repeats).

*Further minor optimization of 1A read-depth measurements.*

We performed one final optimization that slightly improved the accuracy of type-A repeat count estimates by making use of an additional paralogous sequence variant (119bp into the intron downstream of KIV-2 exon 1A, chr6:160616998:T>C in hg38; Extended Data Fig. 2a) that marked a repeat expansion constituting ~30% of all KIV-2 repeat units (averaged across all European alleles), but present only in a minority of alleles (Extended Data Fig. 2b). We labeled the type-A repeats carrying the reference allele “A1” and those carrying the alternate allele “A2”.

Subdividing reads derived from type-A repeats into A1 vs. A2 (vs. ambiguous) groups offered the opportunity to separately estimate A1 and A2 repeat content and then tailor the parameters of our phasing and imputation algorithm separately for the A2 repeat expansion vs. the A1 repeats, potentially improving accuracy. This task was complex but achievable with some care:

- Most read pairs that aligned uniquely to exon 1A do not span the 119bp-downstream variant, so directly assigning reads to A1 or A2 was not feasible.
- However, we could use reads that span the PSV to approximately partition type-A-derived reads according to the observed ratio of T:C bases at the PSV. This approach worked well because ~40% of participants were homozygous for zero copies of A2, such that the partitioning lost no accuracy for such individuals.
- We still needed to restrict reads that contain the REF allele (T) to those that arise from type-A repeats (and discard those that come from type-B repeats). We accomplished this task by separating the PSV-spanning reads into those that aligned unambiguously to 1A sequence (which we counted), those that aligned unambiguously to 1B sequence (which we discarded), and those that did not align uniquely to either (which we pro-rated by the estimated fraction of type-A repeats in the individual).
- This approach produced an approximately unbiased partition of type-A reads into A1 vs. A2 measurements that was fairly noisy (because a typical sample only had ~100 reads that spanned the PSV and could be used for partitioning, in contrast to the ~1000 reads that mapped to type-A sequence). However, both the A1 and A2 measurements phased very well, enabling an improvement in accuracy by repartitioning 1A reads according to post-phasing (“refined”) A1 and A2 estimates and then running another iteration of phasing.

In our final KIV-2 length estimates, we used an average of “1A phased read-depth” obtained with and without this additional optimization (to balance the trade-off between better phasing accuracy after partitioning A1 vs. A2 but more noise in read-depth estimates).

Finally, we note that our genotyping procedure did not explicitly treat type-C repeats. As such, reads generated from type-C repeats contributed to a mixture of the other repeat type counts depending on which type-informative paralogous sequence variants they spanned (which determined whether they aligned best to A1, A2, or B). We expect that any biases introduced by this behavior are negligible given the rarity of type-C repeats (only ~1% of all KIV-2 repeat units^22^).

*Benchmarking accuracy of KIV-2 allele length estimates.*

Our cross-validation-based benchmarks of phasing accuracy (based on pseudo-trios) estimated high accuracy of phased estimates of KIV-2 repeat types A ($\hat{R^{2}}=0.95$) and B ($\hat{R^{2}}=0.98$) and subtypes A1 ($\hat{R^{2}}=0.99$) and A2 ($\hat{R^{2}}=0.99$), with similarly-accurate imputation ($\hat{R^{2}}=0.92, 0.98, 0.97, 0.99$, respectively). Assuming approximate independence of errors across different repeat types, we estimated the RMSE of total KIV-2 length created by combining A+B estimates or by combining A1+A2+B estimates (with appropriate weighting) to be ~1.1 and ~0.9, respectively (using the formula $RMSE\approx\sqrt{\sum_{types} \hat{R^{2}}\cdot\left( allele length variance \right)}$), suggesting that our final KIV-2 length estimates (which averaged estimates from these two methods) likewise achieved RMSE of approximately 1 repeat unit.

We corroborated these benchmarks by separately comparing total KIV-2 allele lengths (diploid VNTR content) derived from our WES-based pipeline to WGS read-depth-based estimates we generated for 48 UK Biobank participants for whom WGS data had independently been generated by sequencing centers at the Broad Institute and BGI. Of the 48 participants, 6 had been exome-sequenced, and we imputed WES-based KIV-2 lengths into the remaining 42 participants (by separately imputing counts of each repeat type).

The WGS read-depth-based KIV-2 length estimates were highly concordant between the Broad and BGI sequencing experiments ($R^{2}=0.99$). Our WES-derived (mostly imputed) estimates also closely matched the WGS-based estimates ($R^{2}=0.95$; Extended Data Fig. 1b).

***ACAN***

*Sequence variation within the ACAN VNTR.*

The 57bp repeat units of the *ACAN* VNTR harbor several common paralogous sequence variants (PSVs) that distinguish repeat units from one another. Two of the most common PSVs affect adjacent bases of the repeat unit which belong to codons for adjacent amino acids; Doege et al. previously used these “diagnostic codons” to classify *ACAN* VNTR repeats into four types^24^ (numbered 1-4; Extended Data Fig. 8a). Examination of the *ACAN* VNTR sequence in the GRCh38 reference and in other long read-based human sequence assemblies available from NCBI (mostly generated by the Human Genome Structural Variation Consortium) showed that repeat type 4 could be further subdivided into four common or low-frequency subtypes based on three additional PSVs 17bp, 20bp, and 26bp downstream of the two “diagnostic” bases distinguishing repeat types 1-4 (Extended Data Fig. 8a).

*Bias in IDT xGen v1 exome capture of ACAN VNTR repeat subtypes.*

The IDT xGen v1 exome capture panel tiled the *ACAN* VNTR GRCh38 reference sequence with 120bp capture probes, with 12 probes falling fully within the VNTR region. Similar to *LPA*, this capture design created biases in counts of reads generated from fragments of *ACAN* VNTR alleles containing repeats of different types. Most of these biases were more subtle than at *LPA* because (i) different repeat types usually differed by only 1-2bp out of 57bp; (ii) the GRCh38 reference contained similar numbers of type-1 repeats (6), type-2 repeats (7), and type-3 repeats (9); and (iii) most European alleles contained few type-4 repeats (e.g., 3 in GRCh38). However, some alleles (and in particular, a 22-repeat allele observed at low frequency in Europeans^24^) carried an expansion of a subtype of repeat 4 not contained in GRCh38 and therefore not targeted by any capture probes, which led to under-counting of this repeat4 subtype by ~20% in our initial analyses.

*Correction of capture bias by separate estimation of copy numbers of repeat subtypes.*

To address this bias as well as subtler biases in counts of other repeat types, we again adopted a strategy of separately genotyping each repeat subtype. In contrast to *LPA*, the *ACAN* VNTR repeat unit (57bp) was shorter than the length of an exome-sequencing read (76bp), so instead of counting reads mapping to repeat subtypes, we counted occurrences of PSV “barcodes” carried by each subtype. For repeats 1-3, these “barcodes” consisted of only the two adjacent diagnostic bases. When we observed a read alignment containing the bases (TA) indicative of repeat 4 at these positions, we further examined the bases 17bp, 20bp, and/or 26bp downstream (if present within the read) to determine whether the read could be assigned to one of the four repeat 4 subtypes. (If the read contained the REF=G base at +17bp, we assigned it to either “rep4_00x” or “rep4_01x” based on only the base at +20bp, and if it contained the ALT=T base at +17bp, we assigned it to either “rep4_1x0” or “rep4_1x1” based on only the base at +26bp (Extended Data Fig. 8a).) We considered reads that either fully aligned or aligned with soft-clipping on one side (CIGAR = 76M or #M#S or #S#M) and required base quality ≥25 at all PSVs used in analysis.

For each the seven repeat subtypes (repeats 1-3 and the four subtypes of repeat 4), we applied our normalization procedure to the “barcode counts” obtained in this manner, producing an analogue of “normalized read-depth” for each repeat subtype. We then applied our phasing algorithm to these normalized barcode-depths to partition these (unscaled) estimates of copy number into their haploid components. (For rep4_00x, we slightly modified the phasing algorithm to account for extreme heteroskedasticity—as the vast majority of alleles carried 0 or 1 copies of this subtype whereas the low-frequency 22-allele carried 8 copies—by adjusting the model to assume that variance in count-based estimates increased linearly with the estimates, basing the parameters of the linear fit on observed differences between estimates in IBD2 sibs.) For all seven repeat subtypes, the phased estimates exhibited multimodal distributions with discrete peaks that scaled to integers upon identifying a suitable scaling transformation, enabling easy calibration of these estimates to absolute copy numbers (Extended Data Fig. 8b).

In addition to genotyping these seven common and low-frequency repeat subtypes, we also genotyped an eighth VNTR variant that amounted to a deletion of the ending repeat of the VNTR (the latter half of which is moderately diverged from all of the other repeat types; Extended Data Fig. 8a). This ending repeat is found in nearly all *ACAN* alleles but deleted in a low-frequency African allele (AF=1% in UKB participants of African ancestry). To genotype loss of this repeat, we simply searched for reads mapping to the *ACAN* VNTR region that contained a unique sequence (AGGACATCAGCGGGCTTCCTTCTGGAGGAGAA) generated by loss of the ending repeat. Carriers were then easily identified as individuals with >10 reads containing this unique sequence.

Combining all of these components (which we independently phased and calibrated to absolute copy-number counts) produced final VNTR allelic copy number estimates given by the formula:

$$ACAN VNTR copy number (haploid)=6.36\cdot rep1+4.88\cdot rep2+31.18\cdot rep3+1.63\cdot\mathrm{rep}4_{00x}+1.78\cdot\mathrm{rep}4_{01x}$$

$$+ 0.08\cdot\mathrm{rep}4_{1x0}+1.89\cdot\mathrm{rep}4_{1x1}-\mathrm{rep}E_{\mathrm{loss}}+2.91$$

The +2.91 constant term at the end mostly arises from the literature canonically considering two additional, more-diverged repeats to be part of the VNTR (despite their sequence divergence); we therefore added 2 repeats to the count for consistency with previous work. The remaining +0.91 constant adjustment corresponds to offsets of +0.49 and +0.42 that we needed to add to counts of repeat 1 and rep4_01x, respectively, to align the modes of their allele distributions to integers. These two repeat subtypes constitute the first and last full repeats of nearly all *ACAN* VNTR alleles (based on the long read assemblies available to us from NCBI), and these first and last repeats are expected to generate fewer reads than interior repeat units due to the pre-VNTR and post-VNTR unique sequences not benefiting from capture enrichment from multiple capture probes, probably explaining these offsets.

*Benchmarking accuracy of ACAN VNTR allele length estimates.*

Our cross-validation-based benchmarks of phasing accuracy (based on pseudo-trios) estimated high accuracy of phased estimates of all *ACAN* VNTR repeat subtypes (estimated $\hat{R^{2}}$ vs. truth of 1.00 (rep1), 1.00 (rep2), 0.97 (rep3), 1.00 (rep4_00x), 0.93 (rep4_01x), 1.00 (rep4_1x0), and 0.95 (rep4_1x1)), from which we estimated (following the same procedure we used with *LPA*) that our combined estimates of (haploid) *ACAN* VNTR copy numbers had an RMSE of ~0.9 repeat units.

Unlike with *LPA*, WGS data from the UK Biobank WGS pilot study was not useful for corroborating our allele length estimates using a different technology (because WGS read coverage at *ACAN* was much lower—and therefore noisier—than WES read coverage at ACAN, which had been greatly boosted by the tiling of the VNTR with 12 exome capture probes). However, we verified that the overall distribution of allele lengths was very consistent with literature (cf. Table 1 of Doege et al.^24^ and Fig. 3 of Horton et al.^25^).

In addition to recovering the previously reported distribution of common alleles, we also identified rare (combined European allele frequency ~0.06%) very long alleles with ~40-45 copies of the VNTR that had not been reported in previous literature. We verified that these alleles belonged to IBD clusters (including a 16-haplotype cluster with ~42 repeats and 7-haplotype and 3-haplotype clusters with ~44 repeats), consistent with true inherited variation.

***TENT5A***

*Short size of TENT5A VNTR alleles produces bias in read-depth measurements.*

The *TENT5A* VNTR was the shortest repeat we studied in detail, consisting of a 15bp repeat unit repeated only 2-7 times per allele^26^. Phased allele length estimates from our initial genotyping of this VNTR produced six clearly distinguishable modes, but the spacing of the modes was uneven, and in particular, the 2-allele appeared to generate much lower read-depth than expected. This behavior was probably driven by mapping bias, as reads spanning the full 30bp of the 2-allele do not map directly to the GRCh38 reference sequence (which contains a 4-allele).

*Improved TENT5A VNTR genotyping by combining spanning-read and read-depth information.*

The above observation suggested an alternative genotyping strategy incorporating read-level information, as smaller alleles (specifically, the 2-allele, 3-allele, and 4-allele) of the *TENT5A* VNTR were consistently spanned by several 76bp reads indicating their presence. (We also searched for evidence of 0-alleles or 1-alleles but did not find any evidence that such alleles existed.) Additionally, 76bp reads that partially overlapped the VNTR could also inform of the presence of an allele of at least 5 repeats or an allele of at least 6 repeats (depending on the precise location of the read alignment relative to the VNTR).

We therefore implemented a hybrid genotyping strategy that combined direct read-level information (used to identify short alleles) with read-depth information (used to estimate the lengths of longer alleles). Specifically, for each individual, we applied the following procedure:

- Identify the minimum- and maximum-length allele indicated by direct read-level evidence (which could be the same allele, indicating a homozygote).
- If the maximum-length allele indicated has length ≤5, set the individual’s (unphased) genotype to be the minimum-length and maximum-length allele.
- Otherwise:
  - If the minimum-length allele has length ≤4, then estimate the length of the longer allele as 5 + 2.5 * (# reads indicating ≥6 repeats) / (# reads indicating ≥5 repeats)
  - Otherwise, estimate the sum of the lengths of the two alleles as twice the above quantity: 10 + 5 * (# reads indicating ≥6 repeats) / (# reads indicating ≥5 repeats).

The intuition behind the ratio (# reads indicating ≥6 repeats) / (# reads indicating ≥5 repeats) was that the 76bp read length almost exactly corresponds to the length of a 5-allele, such that longer alleles (specifically, 6-alleles and 7-alleles) result in generation of additional reads mapping fully within the VNTR (and indicating ≥6 repeats) in addition to reads partially overlapping the VNTR on either side (indicating ≥5 repeats). Dividing (# reads indicating ≥6 repeats) by (# reads indicating ≥5 repeats) allowed an easy adjustment for variation in sequencing depth, leaving just the need for a calibration factor which we empirically estimated to be ~2.5.

*Further minor improvements to TENT5A VNTR genotyping.*

The above strategy provided quite accurate, well-calibrated diploid genotype estimates that we could analyze using our standard phasing and imputation algorithm (which treated all genotype estimates as continuous, real-valued measurements). To obtain a further slight improvement in accuracy, we optimized *TENT5A* VNTR genotyping in two additional ways.

First, we adapted the above approach of estimating lengths of longer alleles to make use of our read-depth normalization pipeline (normalizing read-depths of each individual against read-depths of other individuals who exhibited the closest-matching exome-wide coverage profiles). Specifically, we ran a first round of phasing analysis on the diploid genotype estimates computed as above and identified the subset of individuals deemed to be heterozygous for a 6-allele and a shorter allele. We then calibrated counts of reads indicating ≥6 repeats (normalized for exome-wide sequencing coverage) in each individual against the mean of the corresponding quantity across a matched panel consisting of the (6-allele, shorter allele)-heterozygotes among the 500 individuals with closest-matching exome-wide coverage profiles. This strategy removed the need to calibrate counts of reads indicating ≥6 repeats against counts of reads indicating ≥5 repeats.

Second, we adapted our phasing algorithm to account for the (partially) discrete nature of genotypes we obtained from our hybrid genotyping pipeline. Specifically, to leverage the hard-called allele lengths (derived from read-level information) that our hybrid approach produced in individuals who carried shorter alleles, we modified our phasing algorithm to overwrite allele length estimates with hard-called alleles as appropriate. In detail, during each update of an individual’s phasing, we ran our algorithm as usual but then post-processed the estimated allele lengths as follows: (i) if exactly one hard-called allele was available, we overwrote the shorter of the two estimated allele lengths with the hard-call; (ii) if hard-calls were available for both alleles (such that the only uncertainty was their phase), we overwrote both estimated allele lengths with the hard-calls in sorted order.

*Benchmarking accuracy of TENT5A VNTR allele length estimates.*

Our cross-validation-based benchmarks of phasing accuracy (based on pseudo-trios) and imputation accuracy (based on left-out samples) estimated $\hat{R^{2}}$ vs. truth of 0.98 and 0.95, respectively.

We also benchmarked the accuracy of our allele length estimates in the 48 UK Biobank participants with available WGS pilot data (6 exome-sequenced and the other 42 genotyped at *TENT5A* via our imputation pipeline). *TENT5A* VNTR alleles of all lengths could be directly genotyped from 151bp reads from WGS, so we simply compared diploid genotypes from WGS to those from our WES-based pipeline, observing very high accuracy consistent with our cross-validation results ($R^{2}=0.97$; Extended Data Fig. 1b).

***MUC1***

*Sequence variation within the MUC1 VNTR.*

Alleles of the *MUC1* VNTR contain abundant intra-allelic variation at multiple positions within the 60bp repeat unit^27^. Repeat units within the GRCh38 assembly of the *MUC1* VNTR contain numerous variations of the canonical repeat sequence, partly reflecting this variation and partly reflecting the difficulty of correctly assembling this repetitive sequence: in addition to likely-true PSV variation, the *MUC1* VNTR sequence in GRCh38 also contains several erroneous frameshift-inducing insertions and deletions that prevent the full VNTR from being annotated as being contained within a large exon. For this reason, the IDT xGen v1 capture panel only contained two 120bp probes within the *MUC1* VNTR (because most of the VNTR sequence had been annotated as intronic).

Based on the effects of within-VNTR sequence variation on read-depth-based estimates of diploid VNTR content that we observed at *LPA* and *ACAN*, we expect that the *MUC1* allele lengths estimated by our genotyping, phasing, and imputation pipeline were probably also modestly biased, with repeat units more closely matching the targeted repeats producing more efficient capture (and thereby being slightly overcounted). However, in light of the complexity of intra-allelic variation at *MUC1* and the fact that our analyses of *MUC1* only explored the first-order effects of VNTR length variation (e.g., examining phenotypic effects at the resolution of short vs. long vs. very long alleles, which were clearly distinguished in our allele length estimates), we did not attempt to optimize our read-depth-based genotyping approach beyond slightly adjusting the right endpoint of the VNTR (initially called at chr1:155192051, which we revised to chr1:155192006 after examining the GRCh38 sequence). This change had a negligible effect on allele length estimates.

*Calibrating MUC1 VNTR allele length estimates.*

We did still need to calibrate the allele lengths produced by our genotyping and phasing algorithm (which provided only relative length estimates). To do so, we compared our estimated distribution of phased allele lengths to allele length histograms obtained by previous studies that directly measured *MUC1* allele lengths using gel electrophoresis^28,29^. Previous studies placed the short-allele mode at ~37 repeats and the difference between the short- and long-allele modes at ~40 repeats, so we applied a suitable affine transformation (i.e., scaling and shift) to our estimates to match these values.

*Benchmarking accuracy of MUC1 VNTR allele length estimates.*

Our cross-validation-based benchmarks of phasing and imputation accuracy were very high ($\hat{R^{2}}\geq0.98$), but we expected that these estimates were somewhat inflated because they were unable to assess error introduced by consistent biases from varying exome capture efficiency for alleles carrying different repeat variants.

The *MUC1* VNTR was sufficiently long and variable to be accurately measured using WGS read-depth, so we estimated diploid VNTR content in the 48 individuals in the UK Biobank WGS pilot study by analyzing read alignments using the Genome STRiP pipeline^6^. Comparing these estimates to estimates from our WES-based pipeline for these 48 individuals (6 exome-sequenced, 42 with imputed *MUC1* alleles) showed high accuracy ($R^{2}=0.90$; Extended Data Fig. 1b). This $R^{2}$ value measures both error in our WES-based estimates and error in the WGS-based estimates and thus serves as a likely lower bound on the true accuracy of allele length estimates from our WES pipeline.

***TCHH***

*Optimizing the boundary of the 18bp TCHH VNTR.*

Our initial genotyping of the *TCHH* VNTR counted reads aligning to the repeat region defined by chr1:1521119170-152112225. However, closer inspection of this region showed that it contains two distinct repeats, as previously described^30^: the 18bp (6 amino acid) repeat we wished to genotype, and a 39bp (13 amino acid) repeat to its right (upstream in transcription). The latter repeat appears to exhibit very little variation (and we did not detect association of such variation with male pattern baldness in UK Biobank). In our optimized genotyping, we therefore restricted read-counting to reads containing only repeats of the 18bp sequence, which corresponded to mapping within the region chr1:152111930-152112103. Doing so improved concordance of estimates in IBD2 sib-pairs to $IBD2 R=0.91$.

*Calibrating TCHH VNTR allele length estimates.*

To obtain absolute *TCHH* VNTR length estimates from the relative allele length estimates produced by our genotyping and phasing pipeline, we first estimated the constant offset corresponding to the allele length that would generate zero observed reads aligning within the VNTR. We did so using the following reasoning:

- The GRCh38 reference allele for TCHH contains 8 copies of the 18bp repeat unit (followed by 1 additional copy of the same 18bp repeat that is considered to belong to the adjacent 39bp repeat).
- To align to the region chr1:152111930-152112103, a 76bp read generated from the 8- allele in the reference must start between 152111930 and 152112103 – 75 (inclusive).
- For an allele carrying a different number of repeats $n$, the size of the region on the allele from which such reads can be generated increases by $18(n-8)$.
- Solving for $152111929=152112103-75+18(n-8)$ gives $n=2.5$ as the offset corresponding to the allele length that corresponds to zero observed reads.

Second, we roughly estimated the mean European *TCHH* VNTR allele length as ~8.5 repeat copies based on the long read assemblies available to us from NCBI and a corroborating analysis of insert sizes of paired-end reads from high-coverage WGS of 1000 Genomes EUR samples.

Combining these two pieces of information yielded the affine transformation we applied to obtain absolute estimates of *TCHH* VNTR allelic copy numbers.

*Benchmarking accuracy of TCHH VNTR allele length estimates.*

Our cross-validation-based benchmarks of phasing accuracy (based on pseudo-trios) and imputation accuracy (based on left-out samples) estimated $\hat{R^{2}}$ vs. truth of 0.78 and 0.71, respectively, indicating the difficulty of imputing this VNTR, which is poorly tagged by all nearby SNPs. We expect that these cross-validation estimates of $\hat{R^{2}}$ vs. truth are probably fairly accurate given that alleles of the 18bp *TCHH* VNTR harbor very little intra-allelic variation (and are thus unlikely to suffer exome capture bias); however, data from long-read sequencing will be needed to verify these estimates and to determine the accuracy of our absolute allele length calibration (which is not measured by $R^{2}$).

***RRBP1***

Our VNTR ascertainment pipeline identified a 30bp repeat in *RRBP1* with 4 copies in GRCh38 spanning chr20:17659123-17659247 that our initial analysis indicated exhibited heritable variation ($IBD2 R=0.47$, phasing $\hat{R^{2}}=0.41$) that associated potentially-causally with height. Upon closer inspection of the GRCh38 reference sequence in this region, we observed that the flanking sequence contained a total of ~40 approximate repeats (with varying levels of divergence from the 30bp repeat unit appearing in 4 consecutive perfect copies); however, expanding the region used for read-counting to include the full approximate repeat region reduced sib-pair correlation, suggesting that the repeat variation might be more local.

We attempted to optimize the boundaries of the read-counting region to improve genotyping (as measured by sib-pair correlation) and found that counting reads aligning fully within the region chr20:17659101-17659347 and containing sequence distinguishing the main repeat unit from nearby repeats (TTGCCCTGGTTCTGGGCCCCC) appeared to somewhat improve genotyping and phasing accuracy ($IBD2 R=0.51$, phasing $\hat{R^{2}}=0.56$). However, rerunning the association and fine-mapping analysis using the updated allele length estimates reduced confidence that the association with height was causal (FINEMAP posterior probability = 0.33), so we did not perform further follow-up analyses at this locus.

Further work will be required to elucidate the nature of VNTR variation in *RRBP1* and whether it influences height. A recent study using long-read sequencing indicated that the GRCh38 assembly may be inaccurate at this locus, with *RRBP1* alleles carrying insertions (relative to GRCh38) of 15-16 repeats of a 30bp unit presumably corresponding to this VNTR^31^.

### 5. Genotyping paralogous sequence variants within the *LPA* KIV-2 VNTR

The two coding exons and surrounding splice regions of the *LPA* KIV-2 VNTR contain several common variants and numerous rare variants that have been technically challenging to study because such “paralogous sequence variants” (PSVs) typically only affect 1 out of the 10-30 KIV-2 copies within an *LPA* allele, complicating variant calling from high-throughput sequencing^22,32^. Previous studies have identified three such variants with clear effects of Lp(a) (a stop gain variant in exon 1 [ref.^33^], a canonical splice variant at the +1 donor site of exon 1 [ref.^32^], and a variant in the last base of exon 2 [ref.^34^] that impairs splicing) as well as a variant in the splice region preceding exon 2 that was observed to associate with reduced Lp(a) (*P*=0.0052 in *N*=23 carriers^22^).

We sought to leverage whole-exome sequencing in 49,959 UK Biobank participants (together with phasing and imputation into the remainder of the UK Biobank cohort) to comprehensively explore the landscape of variation within and surrounding the KIV-2 exons and determine the effects of such variation on Lp(a) levels. These analyses, which we describe in this note and the following note, uncovered three additional rarer variants within the region that likely result in Lp(a)-null alleles (Supplementary Table 3), in addition to recovering the effects of the four previously-identified KIV-2 PSV variants (strengthening the evidence for causality of the fourth variant and quantifying the extent to which both non-LoF splice variants reduce Lp(a)). (We also applied similar analyses to the *ACAN* locus for height but did not find any PSVs in the *ACAN* VNTR with strong statistical evidence for causality.)

**Ascertaining and genotyping PSVs from exome-sequencing data**

To ascertain and estimate (diploid) copy numbers of PSVs from exome-sequencing data, we merged read alignments across the seven paralogous copies of KIV-2 exons 1 and 2 and their intronic flanks in the GRCh38 reference (i.e., the six copies formally belonging to the KIV-2 repeat as well as KIV-1 exon 2 and KIV-3 exon 1). We then identified reads supporting non-reference variants, and for each non-reference variant identified within an individual, we estimated its diploid copy number—i.e., the number of KIV-2 repeats containing the variant, summed across the maternally- and paternally-inherited *LPA* alleles—using the formula:

$$diploid PSV copy number=\frac{\# ALT reads}{\# ALT reads+\# REF reads}\cdot(diploid KIV\text{-}2 copy number)$$

where diploid KIV-2 copy number had been estimated according to the genotyping and phasing algorithm described above. Intuitively, this formula allocates the estimated total number of KIV-2 repeats across those carrying the reference vs. non-reference allele at the site in question according to the ratio of the numbers of reads observed. PSV copy number estimates obtained in this way were sometimes downward-biased by capture bias or mapping bias favoring the reference allele, but such biases could be corrected in downstream analysis.

**Phasing and imputing PSV genotypes**

Analysis of KIV-2 PSVs using exome-sequencing data from UK Biobank was particularly technically challenging because the off-target capture of exon 2 and type-A versions of exon 1 (due to the exclusion of these exons from the IDT xGen v1 panel) resulted in generation of only ~1 read per repeat unit at some sites, such that our estimates of diploid PSV copy number tended to be very noisy. Fortunately, the large size of the UKB cohort (49,959 exome-sequenced participants) enabled us to employ haplotype-sharing analysis to simultaneously phase-resolve and refine PSV copy number estimates: intuitively, PSV measurements from individuals who shared long haplotypes (and therefore were likely to share *LPA* alleles) could be analyzed together to help inform estimates of one another’s genotypes.

For 80 common and low-frequency PSVs, the same phasing algorithm that we had applied to VNTRs also accurately phased and imputed PSV genotypes (mean$\hat{R^{2}}$= 0.91 for phasing and 0.88 for imputation).

Rarer PSVs could not be phased using the same algorithm because it relied on measures of concordance of copy number estimates in IBD2 sib-pairs and in pseudo-trios to perform parameter estimation, such that if none of the 133 sib-pairs we determined to be IBD2 at *LPA* carried a rare PSV, then the parameter optimization routine broke down. We therefore implemented a separate pipeline that we applied to phase 85 rare PSVs estimated to have 2 to 400 carriers among the exome-sequenced participants. This pipeline analyzed each such PSV as follows:

- Identify any individual with at least 1 read carrying the ALT allele as a potential carrier.
- For each potential carrier:
  - For each of the individual’s two haplotypes, count the number of other potential carriers among the 200 haplotypes sharing longest IBS (at *LPA*).
  - Call the individual as a homozygous carrier if both haplotypes share long IBS with several other potential carriers (total carrier count >10 across 2 x 200 “haplotype neighbors” of the two haplotypes, with reasonably balanced counts from each neighbor, i.e., <5:1 imbalance).
  - Otherwise, call the individual as a heterozygous carrier as long as one haplotype shares long IBS strictly more potential carriers than the other; if so, assign the PSV to the haplotype with more potential carriers among its IBS neighbors.

After using the above pipeline to phase the 85 rare PSVs in exome-sequenced individuals, we then imputed them into the remainder of the UK Biobank cohort using Minimac4^35^ (treating these rare PSVs as biallelic).

**Optimizing genotyping of common and low-frequency PSVs with large effects on Lp(a)**

An initial round of association and fine-mapping analysis using phased and imputed PSV genotypes from the pipelines above identified five common and low-frequency PSVs that appeared to greatly (or entirely) reduce Lp(a), including the four previously-reported variants as well as a new Tyr>Asp (Y>D) missense variant in exon 1 with MAF=0.3% in European alleles that appeared to produce an Lp(a)-null allele. Given the large effects of these variants and their influence on Lp(a) levels in a sizable fraction of the cohort, we further optimized our genotyping of these five variants to ensure accurate fine-mapping of rarer variants.

We implemented two optimizations within the copy number estimation framework described above, both aimed at increasing the fidelity of initial genotype estimates that were often based on very few reads supporting the ALT allele. First, we applied stringent filters on base qualities and within-read positions (e.g., dropping the last base of each read) that we manually optimized for each of the five PSVs by using samtools^13^ mpileup to examine distributions of these parameters among ALT base calls from confident carriers (with multiple reads containing the ALT base) vs. potential non-carriers (with only one ALT read). Second, we identified and dropped potential duplicate reads that had not been filtered by standard duplicate read analysis (which checks for pairs of paired-end reads that map to exactly the same pair of positions) because multi-mapping reads generated from the KIV-2 region had been randomly mapped to one of the paralogous KIV-2 repeats in GRCh38.

Our cross-validation benchmarks within our phasing and imputation pipeline indicated that the optimized copy number estimates of all five of these PSVs imputed accurately ($\hat{R^{2}}>0.75$). Phased allelic copy numbers of the two most common non-LoF splice-altering mutations (MAF=20% and MAF=13% in Europeans) were sufficiently accurate to identify rarer alleles likely to carry the PSV on two KIV-2 repeats (CN=2; MAF~1%) or three KIV-2 repeats (CN=3; MAF~0.2%; Extended Data Fig. 5).

### 6. Fine-mapping associations of *LPA* variants with lipoprotein(a)

KIV-2 size variation alone explains roughly half of population variation in Lp(a) levels, such that remaining variation in Lp(a) among individuals is largely explained by other sequence variants in *LPA* (that affect Lp(a) levels together with KIV-2 size)^21^. Previous studies have identified several coding and splice variants in *LPA* that create null alleles^32,33,36–39^ (producing no detectable serum Lp(a)), and a few other coding or splice region variants have been observed to associate with substantially reduced Lp(a) (with less certain causality)^22,40^. Additionally, two variants in the 5’ UTR of *LPA* have been observed to associate with more moderate changes in Lp(a) with support from *in vitro* (reporter assay) analyses of translational activity^41,42^. However, a comprehensive list of the *LPA* variants that influence Lp(a) levels in most individuals has remained elusive (as has strong statistical evidence for causality of the variants mentioned above, aside from protein-truncating variants) because of the challenges of genotyping KIV-2 variation and accounting for nonlinear effects on Lp(a).

To comprehensively fine-map *LPA* variant associations to variants likely to causally influence Lp(a) levels, we performed a stepwise conditional analysis using a sequence of statistical tests tailored to iteratively identify likely-causal variants with different characteristics (detailed in the following paragraphs) and condition on these variants in subsequent steps. In Supplementary Table 3, we list each of the 17 steps of this analysis, the likely-causal variant(s) identified, the statistical test performed, the number of alleles tested, the association statistic of the putative causal variant(s), and a description of the association statistics of other *LPA* variants. At each step of the analysis, we performed an association test for all 37,208 variants within 1Mb of *LPA* for which we could obtain or impute genotype information, including:

- SNPs and indels directly genotyped on the UK Biobank SNP-arrays;
- SNPs that we re-imputed (to obtain phased genotype estimates) from the Haplotype Reference Consortium panel r1.1 into the UK Biobank cohort using Minimac4^35^ v1.0.1;
- SNPs and indels that we previously imputed from the WES cohort into the remainder of the UKB cohort^43^; and
- SNP and indel PSVs in the KIV-2 region that we ascertained and genotyped from exome-sequencing read alignments and imputed into the rest of the UKB cohort (as described in the previous note).

We performed stepwise association analyses on subsets of the 337,466 stringently-filtered White British, unrelated UKB participants in all steps except Step 16, in which we analyzed exome-sequenced participants of self-reported African-ancestry (*N*=893).

At each step of this analysis, we tested for association of a variant on one haplotype controlling for the lipoprotein(a) contributed by the allele on the opposite haplotype (on the homologous chromosome) and controlling for the effects of variants discovered in prior steps. In the initial steps (Stage 1; Steps 1-12), we required the allele on the opposite, homologous chromosome to carry a previously identified null allele (carrying rs41272114 or rs41259144 (combined European MAF=0.049), both of which are known to create alleles that produce no detectable serum Lp(a)^36,38^), allowing us to perform association analyses within an effective haploid model of Lp(a). To increase power to detect effects of rarer variants (Stage 2; Step 13) and variants with weaker effects (Stage 3; Steps 14, 15, and 17), we then expanded our analysis to allow the allele on the opposite, homologous chromosome to carry any of the variants identified in Steps 1-12 whose carriers appeared to produce low Lp(a); this criterion was satisfied for 30% of all alleles (see legend of Supplementary Table 3 for precise definition). For variant discovery in African-ancestry samples (Stage 3; Step 16), we placed no restriction on the homologous alleles, and instead modeled Lp(a) from KIV-2 length (non-parametrically) and the variants discovered in the prior steps of analysis as described below.

At each step, we removed alleles carrying large-effect variants discovered in prior steps (except at Step 17, where we specifically restricted to carriers of KIV-2.2 +0G>A (G4925A) to search for sub-haplotypes carrying causal variants). We modeled the effects of small-effect common variants identified in prior steps by including them as covariates in linear regression association tests on log(Lp(a)) (Stage 3; Steps 15-17).

Our stepwise analysis targeted variants in three stages:

*Stage 1, Steps 1-12: Large-effect, common and low-frequency variants*. We first sought to identify large-effect variants that drastically reduced Lp(a) despite occurring on alleles with short or medium KIV-2 lengths (that would usually be expected to produce moderate-to-high levels of Lp(a)). We performed Fisher’s exact tests on variants for a binarized Lp(a) phenotype (low vs. high), restricting to alleles opposite null alleles (see above) and whose KIV-2 lengths were restricted to a particular range. The Lp(a) cutoff and maximum KIV-2 length varied between steps (Supplementary Table 3).

In 12 consecutive steps of analysis, this procedure identified 12 variants each of which was predicted to be protein-altering and achieved the top association statistic in a step of analysis. The variants identified in the first two steps were noteworthy both for their high allele frequencies (European MAF=13% and 20%, respectively) and their genomic locations, which were within the KIV-2 repeat on either end of KIV-2 exon 2 (Supplementary Table 3). The first variant alters the final base of KIV-2 exon 2 and was recently demonstrated to impair splicing at the adjacent splice acceptor site^34^. The second variant is in the splice region preceding KIV-2 exon 2 and was recently identified and observed to associate with reduced Lp(a)^22^ (*P*=0.0052 in *N*=23 carriers). Our analysis provided strong statistical evidence of the causality of this association, and we further observed that this variant was computationally predicted to impair splicing of the nearby splice donor site (SpliceAI^44^ donor loss $\Delta$ score = 0.79).

Interestingly, the computational prediction indicated that the KIV-2 exon 2 -11G>A variant might either simply impair the splice donor site or also create a cryptic splice donor 9bp upstream (causing an inframe insertion introducing an IleSerSer amino acid sequence). To determine if this PSV created an alternative splice site, we examined RNA-seq reads from the GTEx project^45^ (v8), across all donors and tissues, to look for any reads showing evidence of the use of a different splice site at the location of the PSV, but we found no evidence that the PSV created an alternative splice site, suggesting that it simply impairs the canonical splice donor site. (The Genotype-Tissue Expression (GTEx) Project was supported by the Common Fund of the Office of the Director of the National Institutes of Health, and by NCI, NHGRI, NHLBI, NIDA, NIMH, and NINDS. The data used for the analyses described above were obtained from the AnVIL GTEx portal for GTEx v8 on 01/07/21.)

*Stage 2, Step 13: Rare variants producing potential null alleles*. Next, we sought rare variants likely to create additional null alleles (producing undetectable Lp(a)). To do so, we performed Fisher’s exact tests against Lp(a) binarized at the lower limit of the reportable range (3.8 nmol/L), restricting association analysis to alleles opposite the 30% of alleles expected to produce low Lp(a) based on Steps 1-12 (see above). We considered only variants that approximately matched the profile of known null alleles in terms of the frequency of Lp(a) measurements below the reportable range (<3.8 nmol/L) in this setup. (More precisely, 85% of Lp(a) measurements were below 3.8 nmol/L among null allele carriers for which the opposite, homologous allele was expected to produce low Lp(a), so we only considered variants for which the 95% CI of this frequency overlapped 0.85) This analysis identified 11 rare variants with top associations, of which five variants (four protein-altering and one in the 5’ UTR of *LPA*) appeared likely to be causal. Two additional variants were in LD with the protein-altering variants and one variant was in LD with the 5’ UTR variant. The remaining three variants were >100kb away from *LPA* and we were unable to identify plausible causal variants underlying these associations (Supplementary Table 3).

*Stage 3, Steps 14-17: Moderate-effect, common variants.* After excluding alleles carrying large-effect variants (Steps 1-13), we sought to identify common variants with smaller effects on Lp(a). In Steps 14 and 15, we restricted to alleles with KIV-2 length in the interval 11 to 18 opposite alleles expected to produce low Lp(a). Among these alleles, the relationship between KIV-2 length and log(Lp(a)) appeared to be linear. We therefore ran linear regression analyses to test log(Lp(a)) for variant associations including KIV-2 length and KIV-2 length squared as covariates. These analyses identified two additional variants, one in the 5’ UTR (with prior support from *in vitro* studies^42^) and a missense SNP, both of which appeared to exert a moderate and consistent influence on Lp(a) across the KIV-2 spectrum.

We next ran a similar analysis using data from African-ancestry participants, hoping to identify additional Lp(a)-modifying variants that may be less common among Europeans. To maximize power, we created an interim model for Lp(a), fit using measurements in Europeans, so we could use data from all exome-sequenced participants of African-ancestry (and place no restriction on the opposite allele). We first estimated the contribution of a “clean” allele (i.e., one without any previously identified large effect variant) to Lp(a) by averaging Lp(a) measurements for all (European and exome-sequenced) individuals with a clean allele of similar KIV-2 length (within 3 repeat units) whose opposite allele carried a null variant (rs41272114 or rs41259144). We then estimated multiplicative factors for Lp(a)-modifying variants by linear regression on log(Lp(a)) – log(Lp(a) estimated from KIV-2 length). Finally, we computed Lp(a) predictions in African-ancestry participants as the sum of the contributions of the two alleles. In Step 16, we ran linear regression analyses on log(measured Lp(a)) – log(predicted Lp(a)), identifying an additional 5’ UTR SNP (also with prior support from *in vitro* studies^41,42^). Minor allele carriers of this SNP appeared to have modestly increased Lp(a) across the KIV-2 spectrum in both Africans and Europeans (Fig. 2a).

Finally, in Step 17, we performed an analysis similar to Steps 14 and 15 among carriers of the splice modifier variant KIV-2.2 +0G>A (G4925A) and identified a missense SNP (mostly contained on alleles carrying the splice modifier variant) that appeared to reduce Lp(a) (Extended Data Fig. 5).

The variants identified by this multi-stage analysis account for almost all (>97%) of the short and medium European *LPA* alleles (<19 KIV-2 repeat units) that produce small amounts of serum lipoprotein(a) (<6 nmol/L when opposite null alleles carrying rs41272114 or rs41259144). This list of variants appears to be comprehensive of all large-effect, Lp(a)-reducing variants polymorphic in Europeans except those that are very rare. We expect that additional low-frequency and rare null alleles remain to be found in other populations.

### 7. Predictive model of Lp(a) from KIV-2 and fine-mapped *LPA* coding and 5’ UTR variants

We modeled lipoprotein(a) concentration as the sum of the contributions of maternally- and paternally-inherited *LPA* alleles consisting of phased KIV-2 lengths and SNP haplotypes for the coding and UTR SNPs identified by our fine-mapping pipeline (Supplementary Table 3):

$$\mathrm{Lp}\left( a \right)=f\left( {KIV2}_{\mathrm{mat}}, SNPs_{\mathrm{mat}} \right)+f\left( \mathrm{KIV}2_{\mathrm{pat}}, SNPs_{\mathrm{pat}} \right).$$

We did not include other variables in our predictive model because genetic variation at *LPA* explained the vast majority of Lp(a) variance (>80%), whereas other covariates explained very small amounts of variance (e.g., sex (0.2%), age (0.1%), and other loci in the genome (<0.1%)).

The contribution of each (haploid) *LPA* allele appeared to be well-modeled as a “baseline” function of KIV-2 length alone, on top of which coding and UTR SNPs on the same haplotype act via multiplicative factors (Fig. 2a). As such, we modeled each haplotype’s contribution as:

$$f\left( \mathrm{KIV}2_{\mathrm{hap}}, SNPs_{\mathrm{hap}} \right)=f_{\mathrm{baseline}}\left( \mathrm{KIV}2_{\mathrm{hap}} \right) \prod_{\begin{aligned} SNPs with ALT \\ allele on hap \end{aligned}} c_{\mathrm{SNP}}$$

where $c_{\mathrm{SNP}}$ denotes a constant factor indicating the magnitude of the effect of each Lp(a)-modifying SNP that our fine-mapping pipeline identified (Supplementary Table 3).

Empirically, the baseline curve $f_{\mathrm{baseline}}\left( \mathrm{KIV}2_{\mathrm{hap}} \right)$ appeared to have two key properties: (i) an exponential decay that governs the relationship between KIV-2 length and Lp(a) for medium-length and longer alleles; and (ii) an inflection point that causes this relationship to break down for short alleles, with the shortest alleles apparently contributing less to Lp(a) than less-short alleles (Fig. 2a). Property (i) is equivalent to the logarithm of the baseline curve $\log(f_{\mathrm{baseline}})$having a linear asymptote. To fit these behaviors while avoiding overparameterization of the baseline curve, we therefore modeled the logarithm of the baseline curve using a conic section (specifically, a hyperbola), which could produce properties (i) and (ii) using only five parameters:

$$\log\left( f_{\mathrm{baseline}}\left( x \right) \right)=c\sqrt{\left( x-a \right)^{2}+b}+dx+e$$

where $x=\mathrm{KIV}2_{\mathrm{hap}}$. We adopted this functional form only to approximately capture the empirical behavior of the relationship between KIV-2 length and Lp(a) while avoiding overfitting; we do not expect that the form itself or its parameters directly correspond to any particular feature of the underlying biology.

**Model-fitting**

Our main challenge in fitting the parameters of this model from UK Biobank data was the “cropping” of Lp(a) measurements provided by UK Biobank. Specifically, the Randox AU5800 instruments that UK Biobank used to perform immunoturbidimetric assays measuring Lp(a) had a reportable range of 3.8–189 nmol/L. Values of Lp(a) that fell outside this reportable range were indicated “Not reportable at assay (too low)” or “Not reportable at assay (too high)” and not made available to researchers. (While the UK Biobank Data Showcase suggested that these values were available in the Biomarker Assay Extended Dataset, we were unable to find such values in the extended dataset.)

To overcome this obstacle, we restricted our model-fitting analyses to a subset of participants whose Lp(a) measurements were largely unaffected by the cropping issue based on the combinations of *LPA* alleles they carried. Specifically, we began with the set of 40,587 exome-sequenced participants who self-reported European ancestry, passed PC-filtering (Methods) and relatedness pruning (which dropped one member of each ≤2^nd^-degree related pair, prioritizing retaining those with Lp(a) measurements), and had not withdrawn from UK Biobank. We then fit the coefficients of our haploid model $f\left( \mathrm{KIV}2_{\mathrm{hap}}, SNPs_{\mathrm{hap}} \right)$ on the subset of haplotypes for which the “opposite haplotype” (i.e., the *LPA* allele inherited from the other parent) was predicted to produce little or no Lp(a) (<2 nmol/L). This criterion essentially isolated the effect of one *LPA* allele, which both reduced noise in Lp(a) measurements and substantially ameliorated the issue of Lp(a) measurements exceeding the reportable range (as only ~2% of European haplotypes produced >189 nmol/L of Lp(a) on their own, in contrast to ~7% of all (diploid) measurements from European-ancestry participants exceeding 189 nmol/L). This approach was a slight relaxation of the “effective-haploid” model (considering haplotypes opposite an Lp(a)-null allele) that we utilized during fine-mapping; we relaxed the criterion on the other haplotype to “predicted Lp(a) <2 nmol/L” to increase sample size (from ~5% of alleles opposite an Lp(a)-null allele to ~19% of alleles opposite an allele predicted to contribute <2 nmol/L to Lp(a)). While this approach incurred the slight inconvenience of needing a predictive model of Lp(a) to apply to opposite haplotypes during model-fitting, we overcame this difficulty by iteratively fitting the model and updating opposite-haplotype predictions until convergence.

Within each model-fitting iteration, we fit model parameters in two steps:

1. Estimate parameters of the baseline curve (5 parameters) and moderate-effect SNPs (3 multiplicative parameters for the 5’ UTR SNPs rs1800769 and rs1853021 and the missense SNP rs3124784).

   This step estimated the way in which KIV-2 length and common, moderate-effect SNPs determined Lp(a) in the majority of European haplotypes that did not carry a large-effect SNP that greatly reduced Lp(a). We fit the 8 parameters in question (5 parameters for the $f_{\mathrm{baseline}}\left( \mathrm{KIV}2_{\mathrm{hap}} \right)$ curve and 3 parameters for the SNPs) by using Matlab’s fminunc optimization routine to minimize squared error between measured Lp(a) (adjusted for the small predicted contribution of the opposite haplotype) vs. Lp(a) predicted by the model fit, with two additional subtleties.

   First, we restricted model-fitting to KIV-2 length ranges that were largely unaffected by the cropping of Lp(a) measurements to 3.8–189 nmol/L, adjusting the KIV-2 length range according to the alleles of the 3 moderate-effect SNPs. Specifically, for *LPA* alleles carrying the Lp(a)-increasing rs1800769:T minor allele (but not carrying the Lp(a)-reducing rs3124784:A minor allele), we restricted to KIV-2 in the range [14, 24]; for *LPA* alleles carrying neither of these minor alleles nor the rs1853021:A minor allele, we restricted to KIV-2 in the range [14, 23]; and for all other LPA alleles, we restricted to KIV-2 in the range [0, 23].

   Second, for the rare remaining instances in which an allele satisfying the above criteria (and opposite an allele predicted to contribute <2 nmol/L to Lp(a)) still produced measured Lp(a) exceeding the reportable range, we set “measured Lp(a)” to its posterior mean assuming Lp(a) was drawn from a log-normal distribution with $\sigma=0.304$ (which appeared empirically to approximately capture variance in Lp(a) measurements after controlling for genetic effects) and with $\mu$ set to match the fraction of Lp(a) measurements exceeding the reportable range (among alleles with the same Lp(a)-modifying SNP haplotypes and with similar KIV-2 lengths, i.e., in the same 2-percentile bin of KIV-2 length).
2. Estimate multiplicative parameters for effects of large-effect SNPs.

   Most of the large-effect coding SNPs identified by our fine-mapping procedure (including all canonical splice site variants and stop gain variants) produced no detectable Lp(a) in nearly all individuals who carried an Lp(a)-null allele on the opposite haplotype; as such, we set the multiplicative factor $c_{\mathrm{SNP}}=0$ for these SNPs. Six large-effect SNPs appeared to each reduce Lp(a) by >70% but clearly did not completely abolish Lp(a) production: 3 cryptic splice SNPs (two near the exon-intron boundary of KIV-2 exon 2 and rs41270998) and 3 missense SNPs (rs41267807, rs41267809, and rs76144756). We estimated the multiplicative effects of these SNPs by comparing mean measured Lp(a) to mean predicted Lp(a) (predicted based on the 8 parameters fit in step 1) in carriers of these SNPs with KIV-2 length <16 (restricting to shorter KIV-2 alleles to improve signal-to-noise in these estimates.

The parameter values we obtained in this way for the 5 parameters of the baseline curve $f_{\mathrm{baseline}}\left( \mathrm{KIV}2_{\mathrm{hap}} \right)$ were:

$$a=10.11, b=30.18, c=-0.227, d= -0.087, e=7.437$$

The multiplicative factors $c_{\mathrm{SNP}}$ we estimated for the Lp(a)-modifying SNPs are provided in Supplementary Table 3.

**Predicting Lp(a)**

Given phased KIV-2 lengths and *LPA* SNP haplotypes for an individual, our model provides a prediction of Lp(a) as the sum of predicted contributions of each *LPA* allele. These genetically-predicted values of Lp(a) were not subject to cropping to the reportable range of 3.8–189 nmol/L, offering an opportunity to examine the cardiovascular effects of extremely high Lp(a) (e.g., >400 nmol/L). However, for the purpose of assessing the accuracy of our model, we needed to compare predicted Lp(a) to measured Lp(a), so in benchmarking analyses we cropped predicted Lp(a) (after summing across each individual’s two haplotypes). Stratifying by self-reported ethnicity among exome-sequenced individuals, we obtained

$$R^{2}\left( cropped pred. Lp\left( a \right), measured Lp\left( a \right) \right)=0.83 \left( \mathrm{EUR} \right), 0.61 \left( \mathrm{SA} \right), 0.54 \left( \mathrm{AFR} \right), 0.62 \left( \mathrm{EAS} \right).$$

(While model-fitting was performed in a subset of the same European-ancestry participants, overfitting is not a concern given the very small number of parameters fit.) The decreased prediction accuracy in non-European ancestries likely arises from a combination of unmodeled Lp(a)-decreasing variants (with higher non-European allele frequencies) and lower accuracy of phased KIV-2 length estimates in the small fraction of non-European UK Biobank participants.
